# A systematic map of soil invertebrate biodiversity in Canada: Identifying spatial and taxonomic knowledge gaps

**DOI:** 10.64898/2026.09.22.753449

**Authors:** Maheshi E. Dharmasiri, Zoë Lindo, Erin K. Cameron

## Abstract

Soil is one of the most biodiverse habitats on the planet, yet the extent of soil biodiversity knowledge in Canada is unknown. To better understand our knowledge, we systematically mapped soil invertebrate biodiversity knowledge from peer-reviewed literature to identify spatial and taxonomic gaps and to assess whether areas projected to experience different magnitudes of climate change are adequately sampled. A total of 1,176 Canadian sampling locations were identified from 427 articles. We found biases in sampling coverage spatially and in relation to current and projected climatic conditions. The geographic region above 60 degrees North is sparsely sampled. Colder regions (< -0.12 °C) and regions receiving the least (< 646.47 mm), moderately high (2268.67 to 2809.40 mm), and high (2809.40 to 3350.14 mm) amounts of precipitation were under sampled. Regions projected to have the smallest (< 0.86 °C) and greatest (> 1.59 °C) magnitudes in temperature change and regions projected to have moderate (-2.06 to +2.69 mm) to large changes (+2.69 to +7.44 mm) in precipitation were underrepresented. The sampled locations also suffer from taxonomic biases which were skewed toward Nematoda (19%), Annelida (18%), Acari (15%), and Collembola (14%). Our results suggest that methods such as stratified sampling is essential to capture underrepresented regions, taxonomic groups, and an array of current and projected climatic conditions. Additionally, shifting to a multispecies sampling approach is essential for broader soil ecosystem knowledge. Importantly, data from well-sampled areas should be examined to determine how it can be leveraged to advance conservation management of soil invertebrates.

## Introduction

Soil is one of the most biodiverse habitats on earth and it is home to an estimated 59% of terrestrial species (e.g., bacteria, mites, nematodes) (Anthony et al. 2023). Among these organisms, soil invertebrates represent a key group that underpins key ecosystem functions such as nutrient cycling, soil formation, and carbon sequestration and storage (Lavelle et al. 2006; Bardgett and Van Der Putten 2014; Pascual et al. 2015; Eisenhauer et al. 2024). Some of these invertebrates are ecosystem engineers (e.g., ants, earthworms, termites) (Orgiazzi et al. 2016) which can manipulate the soil structure and influence biogeochemical cycles (Wu et al. 2025). The diversity and abundance of soil invertebrates are integral to ecosystem processes; consequently, declines in soil biodiversity can directly impact ecosystem function (Wagg et al. 2014; Zeng et al. 2025). Soil invertebrates are affected by a range of anthropogenic pressures, including intensive agricultural practices, chemical inputs, land use change, invasive species, and climate change, among others (FAO et al. 2020; Beaumelle et al. 2023; Phillips et al. 2024a). These stressors not only alter soil biological communities but also contribute to broader processes of soil degradation that threaten long-term ecosystem resilience and function.

The state of the world’s soils is declining with approximately 33% of soils globally classified as moderately to highly degraded (Smith et al. 2024), with about 25% of agriculture soils being highly degraded (Van Vuuren et al. 2011), which directly drives soil biodiversity losses (Phillips et al. 2024a). However, despite the importance of soils to biodiversity, climate regulation, food production, and ecosystem functioning, soil ecosystems often remain poorly represented within broader biodiversity policy and conservation frameworks. International initiatives such as the Kunming-Montreal Global Biodiversity Framework (KMGBF) (COP and CBD 2022a) have established targets intended to halt and reverse biodiversity losses, outlining key pathways for protecting ecological systems. However, these targets are often framed broadly around terrestrial, marine, and freshwater ecosystems, leaving nations to determine how such priorities are translated into national policy. In Canada, for example, the 2030 Nature Strategy (ECCC 2024) was developed in alignment with KMGBF targets. Unfortunately, soil biodiversity is not directly addressed in Canada’s 2030 Nature Strategy limiting meaningful protections and management strategies for belowground biodiversity.

To date, macroecological studies at the continental scale on soil biodiversity remain somewhat limited, and based largely on soil biodiversity in temperate biomes and a few taxonomic groups (Cameron et al. 2018; Guerra et al. 2020a). Globally, only 37% of areas were identified to overlap with the highest aboveground and belowground biodiversity (top 25%) (Cameron et al. 2019a). This demonstrates that aboveground biodiversity can be a poor predictor of belowground biodiversity (Cameron et al. 2019a). For example, belowground biodiversity may be high in areas where aboveground biodiversity is low, like in the Arctic. Therefore, protection targets that address habitats (e.g., 30x30 (ECCC 2024)) may not capture soil biodiversity if they are interpreted from an aboveground lens, reinforcing the importance of developing targeted approaches for conserving soil biodiversity.

Recent efforts to close key data gaps that impede development of effective soil conservation strategies (Parnell et al. 2025) include global scale harmonized sampling efforts such as Soil BON (Guerra et al. 2021) and the sOilFauna (Mathieu et al. 2022), while soil biodiversity was added as a property to monitor in the LUCAS soil monitoring program that extends across the EU (Orgiazzi et al. 2018). Furthermore, the European Union (EU) has implemented the Soil Monitoring and Resilience Law, a cohesive and integrated framework to address soil as a habitat and conserve its biodiversity (Arias-Navarro et al. 2024; Panagos et al. 2025). This recognition of soil biodiversity including its legal protection is only occurring in the EU (Montanarella 2008; Köninger et al. 2022; Kotschik et al. 2024; Ristok et al. 2025), and in contrast to the EU’s recent efforts, the recent Senate of Canada report on the status of soil health (Senate Canada 2024) fails to directly acknowledge the importance of soil biodiversity, and the direct role it plays in ecosystem services provided by soil (e.g., food production, climate and water regulation, and pest control) (Turbé et al. 2010).

Despite Canada’s early leadership in global biodiversity agreements (Minister of Supply and Services Canada 1995) biodiversity continues to decline (Ray et al. 2021), with particularly severe gaps in belowground knowledge (Cameron et al. 2018). In order to prioritize and protect soil biodiversity in Canada, we urgently need to understand the existing evidence base on soil invertebrate knowledge and then use what we discover to formulate a roadmap for future research, policy, and practice.

Canada is geographically the second largest country in the world, with almost 10 million km of total area (Statistics Canada, Geography Division 2018) representing a diverse landscape, and identifying the sampling coverage of soil organisms across this vast environment is critical to obtain an overview of areas that require additional sampling to detect biological community shifts in the context of global change. Further, if the same taxonomic biases (e.g., soil macrofauna) identified globally are present in Canada (Cameron et al. 2018; Guerra et al. 2020b; Beaumelle et al. 2021, 2023; Phillips et al. 2024a), we will be able to narrow these knowledge gaps by increasing sampling effort on understudied soil invertebrates.

We also need to assess how well the current sampling of soil invertebrates captures regions projected to experience varying magnitudes of climate change across the country. The vast geographic extent results in climate change impacts widely varying across the country (Bush and Lemmen 2019). Changes in precipitation and warming can alter soil invertebrate abundances (Briones et al. 2007; Goncharov et al. 2023), reduce diversity (Gruss et al. 2025), alter compositions of feeding guilds (A’Bear et al. 2014) which can have implications on turnover of older soil carbon pools, decomposition and nutrient cycling, soil biodiversity composition and overall ecosystem functioning (Briones et al. 2007; A’Bear et al. 2014; Goncharov et al. 2023; Bicho et al. 2025).

Here, we conducted the first systematic mapping of soil invertebrate biodiversity metadata across Canada from primary peer-reviewed literature, capturing the extent of our current understanding of belowground data including spatial and taxonomic biases within the data. Our objectives were to: 1) identify spatial gaps, 2) compare current and projected climate change impacts with sampling coverage (i.e., are areas projected to experience different magnitudes of climate change adequately sampled?), and 3) identify taxonomic gaps.

## Methods

### Search strategy

A systematic map was developed prior to starting the review (S1 ROSES for Systematic Protocol) following the reporting standards for systematic evidence syntheses (ROSES) protocol (Haddaway et al. 2017). A systematic map is a tool developed in the social sciences that is now often used in environmental science, which can be used to understand current knowledge or the evidence base in a particular field (Haddaway et al. 2016). The ROSES protocol is a set of standard guidelines to keep the systematic map reliable and transparent. The search string used was adapted from previous soil biodiversity meta-analyses (Thakur et al. 2020; Phillips et al. 2024a), and has three components all joined by AND. The first component included terms used to refer to the soil habitat. The second part included soil organisms, with the specific taxa adopted from past reviews of soil ecology (Phillips et al. 2019, 2024b; Thakur et al. 2020; Beaumelle et al. 2021). Both invertebrate and microbe terms were included because this study is part of a larger project, but for this systematic map only invertebrate data were included. The third part included Canada as a topic or as an address, in order to retrieve all relevant studies conducted in Canada. The address field was included to retrieve studies that did not report the location of the study in the title, abstract, or keywords. In most cases, fieldwork conducted in Canada would have a researcher with a Canadian research affiliation and thus this should capture those articles.

The Boolean search term in Web of Science format (Topic – TS, Address – AD) was as follows:

TS = (soil OR belowground OR below-ground OR peat* OR “litter bags” OR detrit*) AND TS = (biodiversity OR divers* OR organisms OR conservation OR biota OR fauna OR microfauna OR micro-fauna OR macrofauna OR macro-fauna OR mesofauna OR meso-fauna OR animal* OR arthropod* OR invertebrate* OR decomposer* OR detritivore* OR rotifer* OR nematod* OR hypha* OR oligochaet* OR protozoa* OR protist* OR microb* OR bacteria OR fung* OR mite* OR acari OR amoeba* OR tardigrad* OR isopod* OR flagellate* OR archae* OR annelid* OR collembol* OR springtail* OR earthworm* OR micro-arthropod* OR microarthropod* OR mycorrhiza* OR enchytrae* OR ciliat* OR termit* OR lumbricid* OR protura OR diplura OR pseudoscorpions OR ants OR myriapod* OR coleoptera OR larva* OR arachnida OR gastropod*) AND (TS = (canad*) OR AD = (canad*))

We conducted the search on January 9 , 2024, using Web of Science and Scopus. The Web of Science Core Collection was searched using the Science Citation Index Expanded (SCIE) and Present and Emerging Sources Citation Index (ESCI). For Web of Science, the document type was limited to Article, Review Article, Early Access, Book Chapters, Data Paper, and Correction. For Scopus, it was limited to Article, Review, Book Chapter, Book, Erratum, and Data Paper. However, deviating from the original ROSES protocol, no reviews or book chapters were ultimately included. No date restrictions were set, therefore, all indexed articles within the databases were searched. Data was extracted from papers in English only. A total of 35,503 articles were obtained from the search term (S2 ROSES Flow Diagram).

### Screening process

We used Rayyan software (Ouzzani et al. 2016) to screen the articles retrieved from the Web of Science and Scopus. A total of 12,350 were screened by title and abstract. A screening protocol (S3 Screening Protocol) was created to ensure inclusion and exclusion decisions at this stage were consistent among the reviewers (M.D., R.C., R.K., K.J.). To calibrate the screening decisions among the researchers, we calculated Kappa scores using epiR package version 2.0.85 (Stevenson and Sergeant 2025) after screening several hundred of the same articles. Kappa scores were greater than 0.6, which is generally the accepted threshold for screening agreement in systematic reviews and systematic maps (see Haddaway 2019; Castle et al. 2022; Llopis et al. 2023; Li 2024; Oswald et al. 2025). Following title and abstract screening, we reviewed the full text of 1,436 articles (M.D. and K.J.). A justification was provided for each study that was excluded at the full text stage, and a list of articles are provided (10.5281/zenodo.20847097). There were no instances where reviewers had to make a decision with respect to inclusion or exclusion of a study for which they were an author.

### Inclusion criteria

There were three main inclusion criteria for this systematic map. First, the study must be conducted in Canada with a sampling location or description provided. Second, the study must report soil organism data either as occurrence, richness, diversity, abundance, or biomass data at the taxa level (e.g., nematodes, mites etc.) and not by body size (microfauna, mesofauna etc.). Third, the soil organism sampling methodology must capture naturally occurring soil organisms belowground at a given time. The methodologies included were soil cores or rhizosphere sampling locations, litter bags, and emergence cages. Pitfall traps were only included when they were combined with any of the prior mentioned methods to sample a specific taxon because pitfall traps measured organism activity on the soil surface rather than in the soil itself (Beaumelle et al. 2021).

### Data extraction

We extracted the following metadata from each study, information on study timeframe, study type, methods used for identification of soil organisms, organism extraction methods (e.g., Tullgren funnel, Baermann pan etc.), measured biodiversity metrics (diversity, abundance etc.), taxonomic resolution of soil biota studied, soil organism(s) studied, geographic location, and sampling habitat. A data extraction protocol (S4 Full-text data extraction protocol) was created to maintain consistency between reviewers.

The taxonomic groupings used when extracting taxonomic information from the literature were primarily based on the Global Soil Biodiversity Atlas classification (Orgiazzi et al. 2016). Ants, termites, and Coleoptera were recorded separately while all other insects were grouped together. The soil taxonomic groups included were Acari, Formicidae, Coleoptera, Diplura, Annelida, Enchytraeidae, Platyhelminthes, Gastropoda, Opiliones, Isopoda, Myriapoda, Nematoda, Protura, Pseudoscorpiones, Rotifera, Scorpiones, Araneae, Tardigrada, Isoptera, and other insects. Additional categories were added as needed during the extraction if the reported soil invertebrates did not fit the above categories.

Geographic data was extracted at the finest possible scale for each study (e.g., extracted sampled point over study site). For papers that provided a location but no GPS coordinates, the coordinates were obtained from Google maps. At this stage, any habitat that did not fit the existing habitat categories was classified as ‘Other’. After the extraction was completed, these were revisited and were added to existing categories by expanding the category descriptions (see S5: Full-text data extraction protocol for habitat category descriptors). For articles that did not provide habitat descriptions (n = 9), article objectives and context was reviewed to assign the most probable sampled habitat. Some studies did not distinguish between methods used to sample an organism (e.g., soil cores and aerial sampling ; n = 7), or specify which organisms were collected from which country in global studies, (e.g., Canada and United States; n = 4), or reported data from combined habitats (e.g., soil and moss ; n = 6). The sampling location data for these studies were extracted because it was conducted in Canada but the taxonomic data was too vague to be included for the taxonomic analyses (see Pooled data in S4: Full-text data extraction protocol). For the spatial analysis, we identified the unique sampling locations by filtering out locations that had the same latitude, longitude, habitat, and taxa.

### Statistical analysis

All statistical analyses were completed using R version 4.5.2 (R Core Team 2025) and RStudio (Posit Software 2025). The Canada Albers Equal Area Conic (ESRI 102001) was used for all area calculations.

We mapped the sampling locations in relation to the 15 terrestrial ecozones in Canada using GIS data from National Ecological Framework for Canada (Government of Canada 2017). Canada’s land can be classified into 15 broad terrestrial ecozones, characterized by geological, biological, physical and anthropogenic characteristics (Ecological Stratification Working Group 1995). These high level ecozones are frequently used to set conservation priorities and make resource management decisions (Deguise and Kerr 2006; Fyson et al. 2024a; Wotherspoon et al. 2024; McFarlane et al. 2025), and thus were chosen for our analysis. To determine whether there was spatial sampling bias of soil biodiversity across Canada, we conducted a chi-square test by comparing the observed number of sampling locations in each ecozone with the expected number of sampling locations based on the proportion of area of each ecoregion (Cameron et al. 2019b). All intervals used for analyses were created using ClassInt package version 0.4.11 (Bivand 2025). The sf package version 1.0.21 (Pebesma 2018; Pebesma and Bivand 2023) was used for ecozone shapefile projections and extracting the ecozone for each sampling location. We also examined the mean annual temperature and mean annual precipitation for the climate normal period (1991 – 2020) at a 1 km resolution (Wang et al. 2016; AdaptWest 2022), to assess spatial bias of the sampling locations for average climate conditions and again conducted a chi-squared test. In this case, we divided the range of values for temperature into five equal intervals with increments of 5.56°C and for precipitation into seven equal intervals with increments of 541 mm. These categories were chosen to ensure that the number of sampling locations in each category were not clustered in one category, and no expected category had values that were too low. We calculated the expected number of sampling locations per category based on the proportion of the land area in a given category. The terra package version 1.8.60 was used for coordinate reference system projections and for raster data extraction for current and projected temperature and precipitation analyses (Hijmans 2025).

Additionally, to determine whether areas which are projected to experience different magnitudes of impact due to climate change are adequately sampled, we used a Chi-square test to assess the spatial bias. Mean annual temperature and total precipitation anomaly data from 2025 to 2100 at a spatial resolution of 10 km was used under the moderate emissions scenario (RCP 4.5). The projected climate data was a multi-model ensemble and we chose the 50^th^ percentile (median) for all our projected climate change analyses with a reference period from 1986 to 2005 (Canadian Centre for Climate Services 2020). This is an intermediate scenario, requiring a medium level of adaptation and the global average warming is expected to plateau between 1.7 to 3.2°C by 2090 (Van Vuuren et al. 2011). We divided the ranges into five equal intervals with increments of 0.24 degrees and 4.75 mm for temperature (°C) and precipitation (mm) respectively. In the Chi-squared test, we compared the observed number of sampling locations in these categories with the expected number per category, which we calculated based on the proportion of the land area projected to change within that category.

We extracted 25 different soil taxonomic groups at different taxonomic levels (e.g., Collembola, mites, earthworms etc.). All these groups were collapsed to the phyla level to present descriptive results.

## Results

A total of 1,176 unique Canadian sampling locations were obtained from the 427 articles published between 1972 and 2024. We identified seven different soil phyla which included Annelida, Arthropoda, Mollusca, Nematoda, Platyhelminthes, Rotifera, and Tardigrada. The total number of publications that reported soil sampling on Arthropoda, Annelida and Nematoda increased steadily over the 42-year time span, while soil sampling related to other phyla grew only incrementally and added relatively few sampling locations. The largest number of publications that reported soil invertebrate sampling in a single year was 48 in 2018.

Sampling effort across ecozones was significantly biased (Table S5: □^2^ = 14,822; df = 14; P < 0.001). The northern ecozones were sparsely sampled in comparison to the southern ecozones. Mixedwood Plains in southern Ontario was most sampled ecozone (Figure 1). The three largest ecozones (the Boreal Shield, Northern Arctic, and Taiga Shield) make up nearly half of the terrestrial land mass in Canada, but only 9% of the sampling locations occurred within these ecozones.

**Figure 1.**
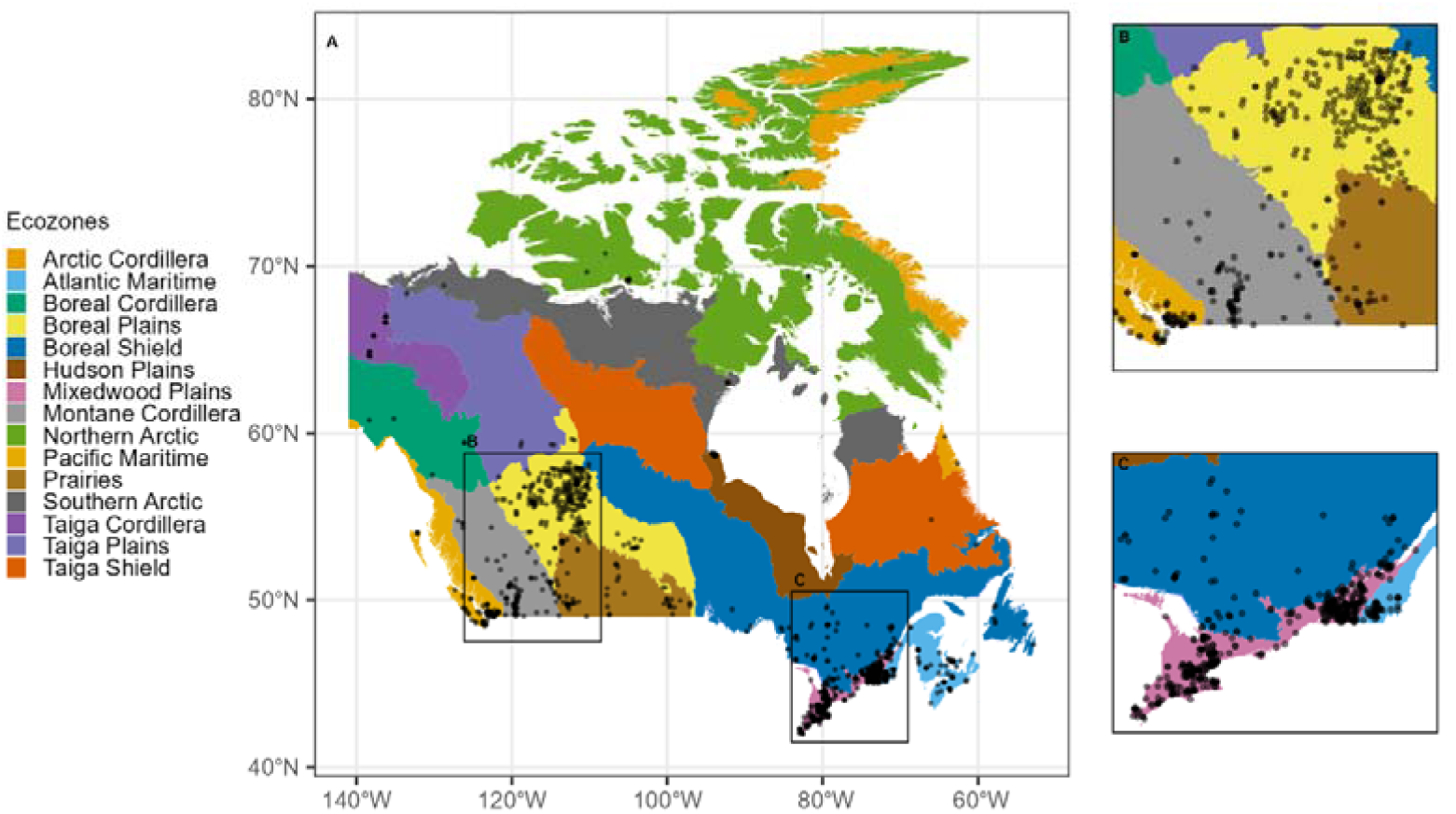
The number of sampling locations found in each of the 15 terrestrial ecozones of Canada. Mixedwood Plains was the most frequently sampled ecozone (n = 439) while Taiga Shield was the least sampled ecozone (n = 1). Ecozone base layer from (Agriculture and Agri-Food Canada 2017) was cropped to fit the cartographic boundaries of Canada from (Statistics Canada 2023) using ArcGIS Pro Software (2024) (version 3.4.0). The coordinate system is WGS 84 (World Geodetic System 1984) and the coordinates are in decimal degrees. The map was created in R Core Team (R Core Team 2025) and Posit Software, PBC (2025) by Maheshi Dharmasiri.

Sampling locations were significantly spatially biased with respect to mean annual climate (Table S6: Temperature: □^2^ = 8, 237.40; df = 4; P < 0.001; Precipitation: □^2^ = 528.79; df = 6; P < 0.001). Colder regions (> -0.11°C) were vastly underrepresented representing 67% of the total landmass (Figure 2). The hottest areas (mean annual >5.45°C), comprising approximately 4% of the land mass, were overrepresented with a total of 54% of sampling locations. Areas with the least precipitation (mean annual < 646.47 mm) representing 69% of total area was the second most sampled category but remains underrepresented in relation to the proportional area (Figure 3). Areas receiving moderately high (2268.67 to 2809.40 mm) to high amounts of precipitation (2809.40 to 3350.14 mm) were also underrepresented.

**Figure 2.**
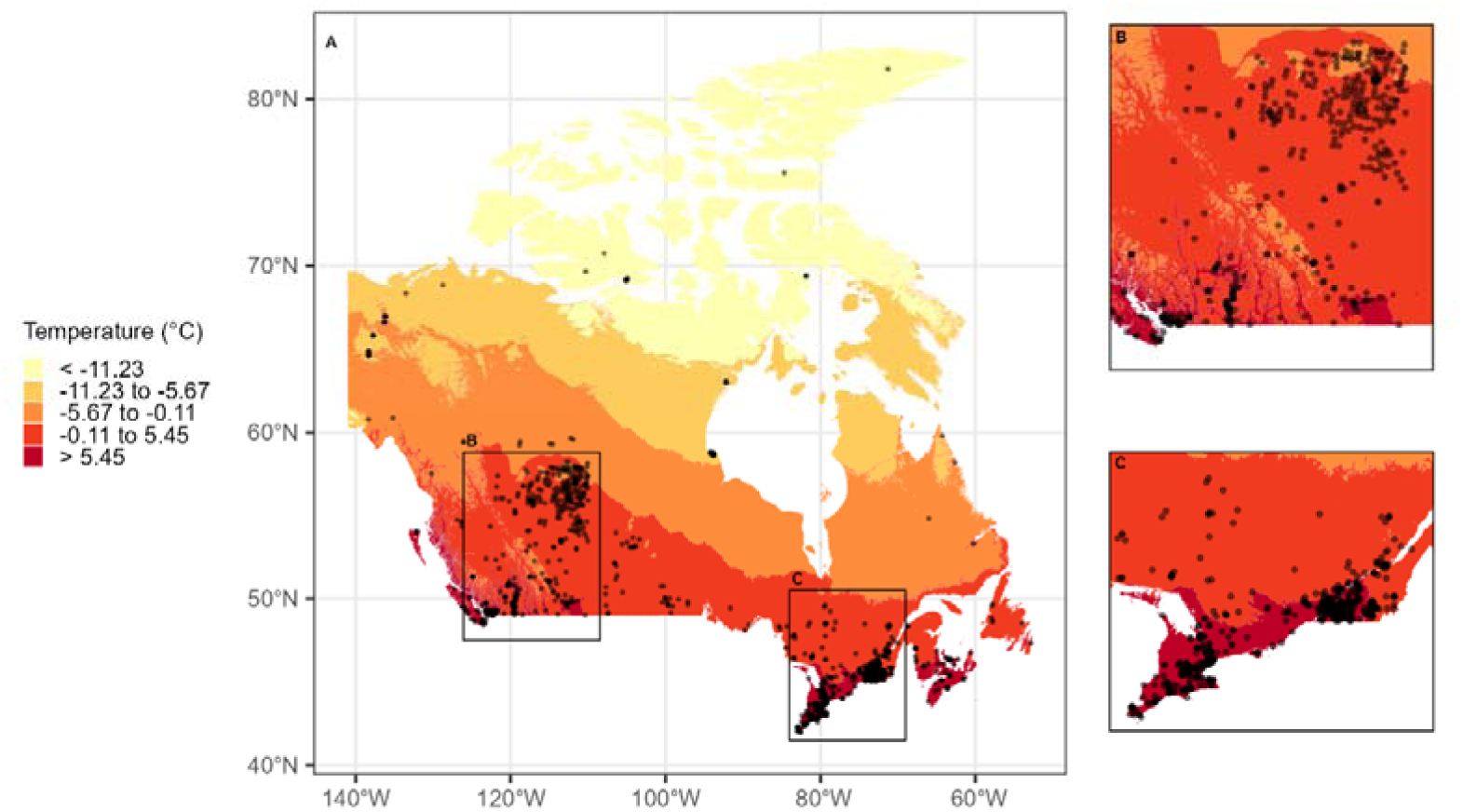
The annual temperature in degrees from the 1991 – 2020 climate normal period. The semi-transparent black circles are the sampling locations. These circles appear darker when overlapping with multiple sampling locations. The inset maps (B) and (C) show areas with the largest number of sampling locations. Current temperature base layer from Wang et al. (2022) was cropped to fit the cartographic boundaries of Canada from (Statistics Canada 2023) using ArcGIS Pro Software (2024) (version 3.4.0). The coordinate system is WGS 84 (World Geodetic System 1984) and the coordinates are in decimal degrees. The map was created in R Core Team (R Core Team 2025) and Posit Software, PBC (2025) by Maheshi Dharmasiri.

**Figure 3.**
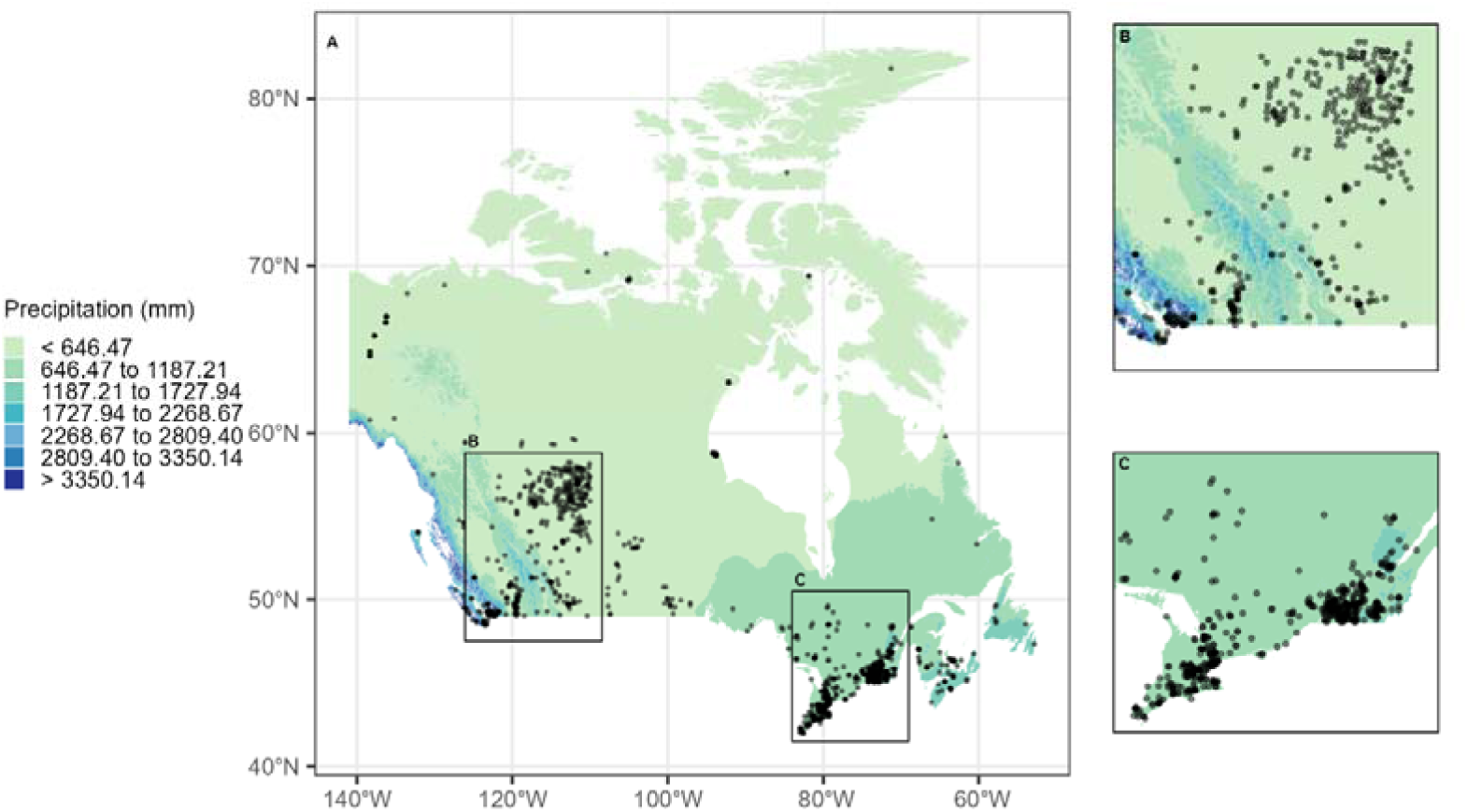
The annual precipitation in mm from the 1991 – 2020 climate normal period (A). The semi-transparent black circles are the sampling locations. These circles appear darker when overlapping with multiple sampling locations. The inset maps (A) and (B) show areas with the largest number of sampling locations. Current precipitation base layer from Wang et al. (2022) was cropped to fit the cartographic boundaries of Canada from (Statistics Canada 2023) using ArcGIS Pro Software (2024) (version 3.4.0). The coordinate system is WGS 84 (World Geodetic System 1984) and the coordinates are in decimal degrees. The map was created in R Core Team (R Core Team 2025) and Posit Software, PBC (2025) by Maheshi Dharmasiri.

Sampling effort in the context of projected mean annual climate change was significantly spatially biased for both temperature and precipitation (Table S7: Temperature: □^2^ = 64.24; df = 4; P < 0.001; Precipitation: □^2^ = 53.32; df = 4; P < 0.001). Areas projected to have a very small change (< 0.86 °C), moderate change (1.11 to 1.35 °C) or a very large change (> 1.59 °C) in temperature were underrepresented (Figure 4). These two categories collectively represent 47.14% of the total area. The two categories representing small change (0.86 to 1.11 °C) and large change (1.35 to 1.59 °C) together cover a spatial extent of 53% of the land mass and are both overrepresented, with a total of 62% of sampling locations. Areas projected to experience the most severe (< -6.82 mm), large declines (-6.82 to -2.06 mm) and greatest increases in precipitation (> 7.44 mm) received greater sampling attention (Figure 5). Categories capturing moderate (-2.06 to +2.69 mm) to large changes (2.69 to +7.44 mm) in precipitation represent 77% of land mass but were both underrepresented in sampling effort.

**Figure 4.**
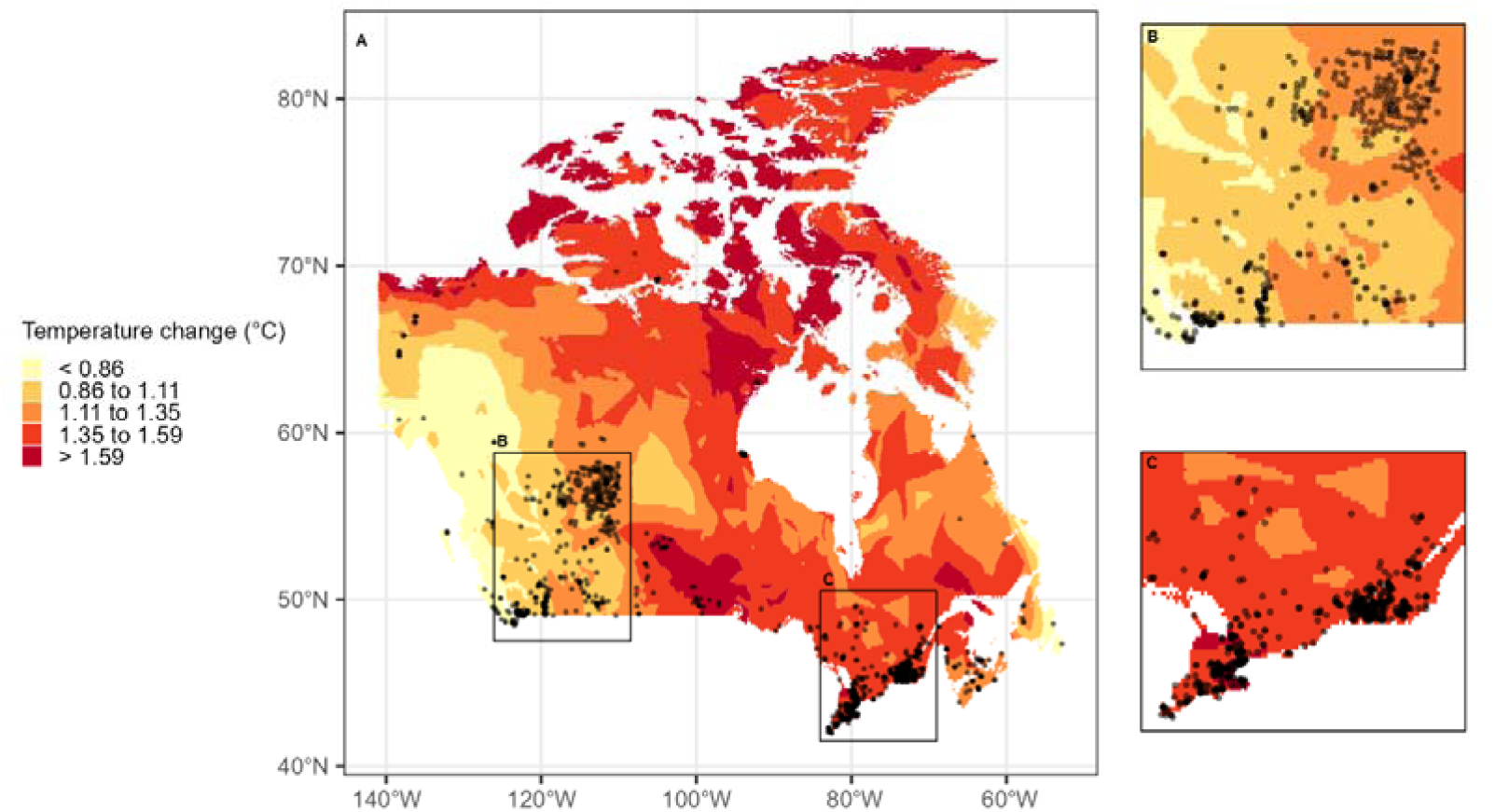
Annual projected temperature anomaly data for the 2025 to 2100 period under the RCP 4.5 scenario (Canadian Centre for Climate Services 2020). The reference period for the projected climate models was from 1986 to 2005. (A) The main map shows the sampling locations across Canada. The inset maps (B) and (C) show areas with the largest number of sampling locations. The semi-transparent black circles are the sampling locations with overlapping locations appearing darker. Projected mean annual temperature anomaly base layer from Canadian Centre for Climate Services (2020) was cropped to fit the cartographic boundaries of Canada from (Statistics Canada 2023) using ArcGIS Pro Software (2024) (version 3.4.0). The coordinate system is WGS 84 (World Geodetic System 1984) and the coordinates are in decimal degrees. The map was created in R Core Team (R Core Team 2025) and Posit Software, PBC (2025) by Maheshi Dharmasiri.

**Figure 5.**
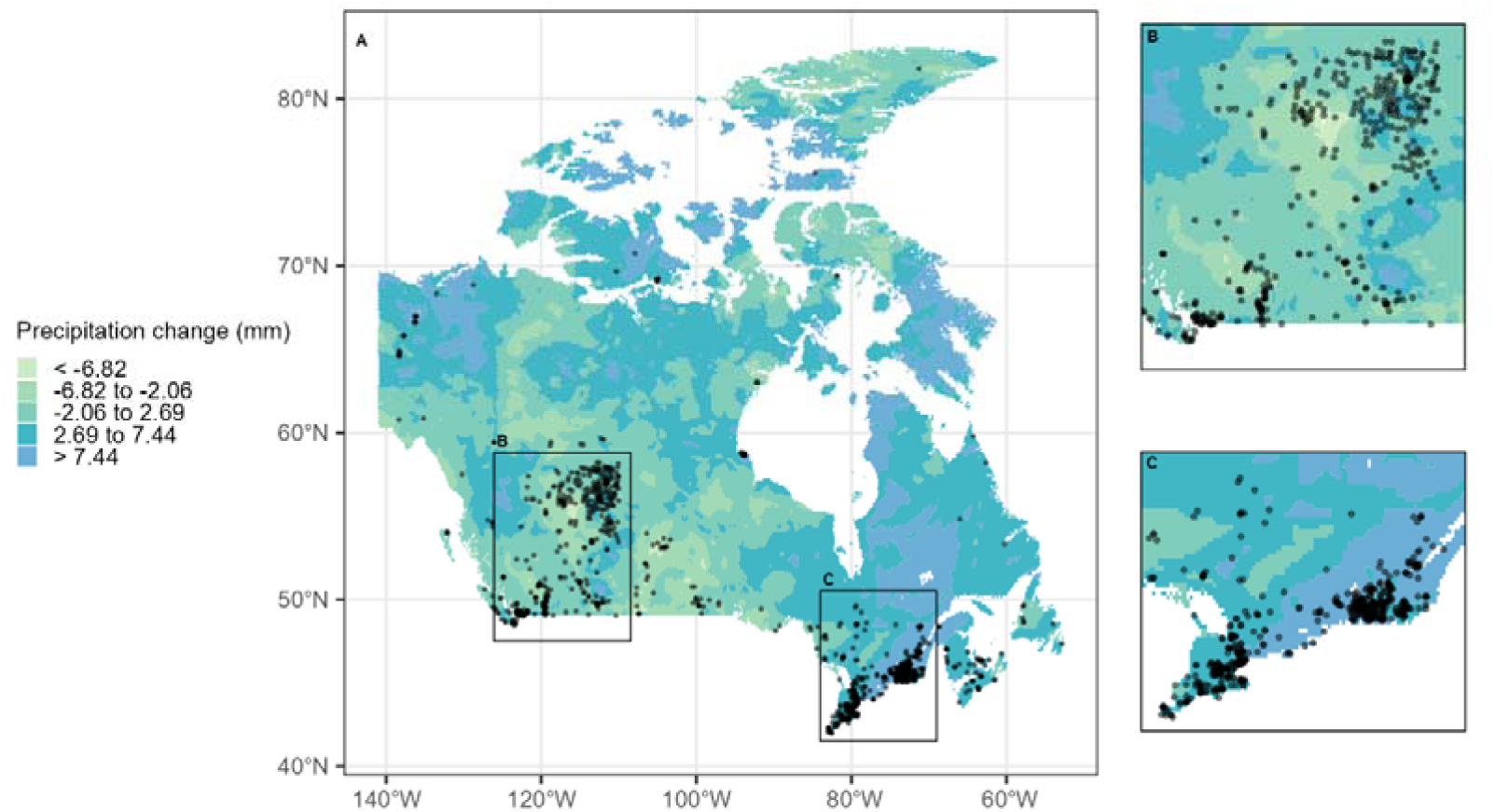
Annual projected precipitation anomaly data for the 2025 to 2100 period under the RCP 4.5 scenario (Canadian Centre for Climate Services 2020). The reference period for the projected climate models was from 1986 to 2005. (A) The main map shows the sampling locations across Canada. The inset maps (B) and (C) show areas with the largest number of sampling locations. The semi-transparent black circles are the sampling locations with overlapping locations appearing darker. Projected mean annual precipitation anomaly base layer from Canadian Centre for Climate Services (2020) was cropped to fit the cartographic boundaries of Canada from (Statistics Canada 2023) using ArcGIS Pro Software (2024) (version 3.4.0). The coordinate system is WGS 84 (World Geodetic System 1984) and the coordinates are in decimal degrees. The map was created in R Core Team (R Core Team 2025) and Posit Software, PBC (2025) by Maheshi Dharmasiri.

The most sampled phyla were Arthropoda (57%), Annelida (21%), Nematoda (19%), and while the least sampled were Tardigrada (1%), Mollusca (1%), Rotifera (0.81%), and Platyhelminthes (0.40%). The phylum Arthropoda was comprised of Acari (26%), Collembola (25%), Coleoptera (14%), other insects (11%), Arachnida, (7%), Myriapoda (6%), Pseudoscorpiones (3%), Protura (2%), Diplura (2%), ants (2%), Crustaceans (1%), Isopoda (1%), and less than 1% of studies were on Scorpiones, Isoptera, and Malacostraca. Within the phylum Annelida, there was more research related to earthworms (86%) than enchytraeids (11%).

Species (48%) was the most frequently used taxonomic resolution in the literature, primarily applicable to Arthropoda (44%), Annelida (31%) and Nematoda (24%). In Arthropoda, Acari (34%), Collembola (24%), and Coleoptera (22%) were the groups mostly identified to species. Within Annelida, earthworms were mostly identified to species (97%), while enchytraeids were not (3%).

Abundance was the most frequently measured variable (45%), followed by biodiversity metrics (e.g., diversity, richness) (32%) and biomass (9%). At the phyla level, abundance (55%) and diversity (60%) was mostly reported for Arthropoda while biomass was mostly reported for Annelida (59%). At more specific taxonomic resolutions, abundance was predominantly reported for Nematoda (23%) while diversity was reported for Acari (24%). Biomass was largely reported for earthworms (54%). Occurrence data was also published for many groups (14%).

## Discussion

Soil invertebrate biodiversity sampling in Canada is highly spatially biased, concentrated in southern and more densely populated provinces, with little to no sampling coverage across much of northern Canada. Consequently, current knowledge of Canadian soil invertebrate biodiversity remains limited, with approximately 4.28 × 10^-3^ articles published per 100 km^2^. These patterns reflect broader logistical, infrastructural, and historical biases in biodiversity research, where accessibility and proximity to research institutions strongly influence sampling effort, and the lack and decline of taxonomist expertise in Canada (Council of Canadian Academies 2010) . However, they also reveal a critical mismatch between sampling intensity and ecological importance.

Our ecozone analysis demonstrates that sampling coverage is not proportional to ecozone extent. The Boreal Shield, Canada’s largest ecozone and one of the world’s most extensive forested systems, remains substantially under-sampled relative to smaller and more accessible ecozones such as the Mixedwood Plains. While large portions of the boreal region remain remote and comparatively less disturbed by direct anthropogenic activity and human infrastructure (Andrew et al. 2012, 2014), these ecosystems are increasingly affected by climate change, resource extraction, altered disturbance regimes, and expanding northern development (Schindler and Lee 2010). The lack of baseline biodiversity data severely constrains our ability to detect ecological change, assess resilience, or understand how belowground communities respond to environmental disturbance across these landscapes.

The Arctic ecozones were also under represented in sampling effort, which is especially concerning given that these regions are warming at nearly three times the rate of the global average (Bush and Lemmen 2019). Unlike many aboveground taxa, soil biodiversity does not necessarily decline with latitude, suggesting that Arctic soils may support high levels of biodiversity that are at risk from this warming (Cameron et al. 2019a). This mismatch between above- and belowground diversity patterns highlights an important limitation of current conservation frameworks, where aboveground biodiversity conservation efforts may not adequately protect rare, unique, and high soil biodiverse hotspots and ecosystems. The lack of baseline soil biodiversity data across Arctic and Boreal ecozones therefore represents a major knowledge gap that constrains our ability to detect ecological change and to develop effective management strategies in regions experiencing rapid environmental transformation.

The Mixedwood Plains ecozones contained the highest concentration of sampling locations, likely reflecting well-known accessibility biases in biodiversity research (Hughes et al. 2021) towards agricultural areas (Geldmann et al. 2016), urban centres, and areas near existing infrastructure and population centres (Sobral-Souza et al. 2024). The Mixedwood Plains is Canada’s most densely populated ecozone and contains one of Canada’s major agricultural regions (Fyson et al. 2024a); therefore, the large number of sampling locations found in the Mixedwood Plains is likely due to accessibility and proximity to agricultural and urban centers. This is supported by our findings that 54% and 25% of the sampling locations from this ecozone are from agricultural and urban habitats, respectively. The population density and cultivation puts this ecozone at high risk from environmental degradation and pollution, especially with a shift towards practices such as row cropping which reduces biodiversity (Fyson et al. 2024b). This underscores the need to establish soil biodiversity baselines in this region; the sampling locations identified from our systematic map do not offer a precise roadmap to conservation management, but they likely provide more than sufficient baseline data to support management in this ecozone.

We also observed large biases in habitat sampling overall in our dataset with most of the sampling taking place in woody (34%) and agricultural (29%) habitats. Within these habitats, the most sampled invertebrates were Nematoda, Annelida, Acari, Collembola, Coleoptera, and other insects. Annelida (Pérès et al. 2011; Li et al. 2021), Acari (Manu et al. 2019), Nematoda (Yang et al. 2021), and Collembola (Ponge et al. 2003) are commonly studied taxa in part because they are used as environmental indicators. For example, the maturity index of nematodes classifies the type of nematodes occupying a habitat based on their reproduction strategy (colonizers employing r-strategies vs persisters with k-strategies), which is used as a biomonitoring tool to assess soil ecosystem health and quality (Bongers 1990). A review conducted by Potter and McKeown (2003) state most of the knowledge on Nematoda in Canada comes from agricultural contexts (Potter and McKeown 2003) with little surveying conducted outside of managed soils. Our findings also corroborate this, with a substantial number of Nematoda samples occurring in agricultural habitats (46%). Nematoda are also likely sampled most frequently in these habitats due to the fact that nematode plant-parasitic relationships can impact agricultural output (Subbotin and Chitambar 2018; Forge et al. 2022; Madani et al. 2024). Similarly, research on Coleoptera in Canada is predominantly focused on beetle-plant feeding associations, relating to economic costs in the agricultural sector (e.g., Goulet 2003). This is consistent with the sampling locations of Coleoptera primarily occurring in agricultural habitats as well (68%). While certain taxa are unsurprisingly studied more often in specific contexts, this highlights the need to broaden soil taxa sampling across diverse habitats and contexts, as responses can vary among organisms. The limited range of habitats sampled is problematic because soil organisms can be extremely localized (e.g., less cosmopolitan species and high endemicity) (Wu et al. 2011; Jorna et al. 2023), and soil organisms are not uniformly distributed, varying vertically and horizontally (Ettema 2002). The distribution of soil organisms can span spatial scales from a few millimeters to several hectares (Ettema 2002).

Our analyses also show that the sampling locations are spatially biased with respect to current and projected climate conditions. The unbalanced sampling across current conditions is concerning because it means that we lack knowledge of soil biodiversity baselines. This is an integral component of the Kunming-Montreal Global Biodiversity Framework (decision 15/5) (COP and CBD 2022b). These baselines are required to understand how soil biodiversity is shifting over time (e.g., phenology, ranges, biodiversity composition). Yet, in Canada there is very limited long-term soil biodiversity data, beyond that collected by the Alberta Biodiversity Monitoring Institute (ABMI); a single study conducted by this institute contributed the largest number of sampling locations, from a large range of Albertan habitats, to our systematic map. ABMI samples many taxa, however, it is restricted to only a few belowground taxa such as oribatid mites. A Canada-wide Biodiversity Observation Network (CAN BON) (Gonzalez et al. 2025) has been proposed which would collect baseline and long-term biodiversity data and therefore would increase the representation of soil habitats in biodiversity programming across the country.

Increased efforts also need to be directed towards sampling areas that will experience different magnitudes of climate change. Regions that are expected to have smaller magnitudes of change can serve as climate refugia (Keppel et al. 2024; Morelli et al. 2026). We identified that areas expected to have the least projected temperature changes were underrepresented. Identifying climate refugia that have intact soil ecosystems and minimal human stressors can serve as priority areas for conservation. Future sampling campaigns not only need to consider climate, but other key drivers of soil communities to fully understand the diversity and distribution of soil invertebrate communities across Canada. For instance, soil type is a key determinant of soil organism distributions, yet the full range across Canada is not captured under the current sampling locations (see S8). Stratified sampling should be used to capture a wide range of variables such as habitat diversity and current and projected climatic conditions at a study site. It can also be more a suitable approach to capture organisms that are unevenly distributed within an environment. For example, stratification facilitated easier distinction of nematode communities, including those found in in low abundances and only in a few samples (Arieira et al. 2016).

In addition to large spatial biases across Canada, our results also show taxonomic biases. The most sampled soil phyla were Arthropoda, Annelida, and Nematoda whereas the least studied were Mollusca, Rotifera, and Platyhelminthes. Multiple reviews focusing on various aspects of soil biodiversity (e.g., global change, chemical stressors, ecosystem function) at the global scale have similarly reported a bias towards Nematoda, Annelida, Collembola, Acari, and Coleoptera (e.g. Guerra et al. 2020b; Beaumelle et al. 2021; Phillips et al. 2024a). However, Martin et al. (2025) reports contrasting biases in the context of forest disturbances with Annelida, Enchytraeids, Isoptera, and Nematoda being substantially understudied relative to their global biomass, while numerous studies were reported with Collembola, Acari, and Coleoptera.

The taxonomic resolution and biodiversity metrices reported for the soil invertebrates also follow similar biases to the sampling data. More granular taxonomic levels and biodiversity data were available for Arthropoda, Annelida, and Nematoda phyla. However, within the Arthropoda phyla, we have insufficient data on many soil invertebrate groups. Future sampling needs to focus on collecting a combination of data including diversity, abundance, and biomass, when possible, to gain a more holistic understanding of the ecosystem at the sampling location. Additionally, most of the sampling locations reported on a single invertebrate taxon (74%), but a shift to multi-taxon sampling is needed to understand the communities present within these locations.

The habitat and taxonomic biases demonstrate that we need to shift away from the human benefit perspective when making conservation and policy decisions. The knowledge clusters we have identified predominantly come from habitats and taxa that have direct economic ties to humans, leading to large knowledge deficits in a diverse range of landscapes and soil invertebrates. Conservation of soil biodiversity should consider economic value in conjunction with intrinsic value, which can help focus conservation efforts on species that are in imminent danger from extinction as opposed to those with a known economic impact, leading to simpler and more equitable management (Decaëns et al. 2006; Phillips et al. 2020).

## Limitations of the map

We acknowledge that our review is not comprehensive because the literature search was limited to peer-reviewed articles and the English language. We recommend future reviews and systematic maps incorporate grey literature and French language literature in their searches. Despite these limitations, our systematic map provides strong evidence of knowledge gaps and clusters within the field of soil invertebrate biodiversity in Canada.

## Conclusion

This is the first systematic map conducted on knowledge gaps on soil invertebrate biodiversity in Canada. We describe the extent of the knowledge from a spatial, climate change, and taxonomic context. The knowledge gaps and clusters identified in this map will allow land managers, conservationists, policy makers, and research groups to formulate plans to narrow the identified knowledge gaps while simultaneously leveraging established knowledge clusters to advance belowground conservation management in Canada. The world’s soils are at risk, and Canada is no exception; it is critical we act now to protect and sustainably manage belowground organisms that form the foundation of healthy soils.

## Supporting information

supplementary files_S1_ROSES for Systematic Map Protocol

supplementary files_S2_ROSES flow diagram

supplementary files_S4_List of articles at full-text stage and extracted data_2

## Acknowledgements

We thank Roman Kisseljov, Rachel Corney, and Kaveen Jayawardane for article screening and data extraction, and Daniel Wilson for assistance with consolidation of sampling locations.

## Data availability

Data generated or analyzed during this study are available in the Zenodo repository, **[10.5281/zenodo.20847097,** https://zenodo.org/uploads/20847097?token=eyJhbGciOiJIUzUxMiJ9.eyJpZCI6IjM5NzNhNzlkLT NhOTMtNDdkYi1hYTEyLTAzZWE0ZTdjM2U3MyIsImRhdGEiOnt9LCJyYW5kb20iOiIzMzg1NDM4O DA1M2U5MTcwOWRkNjhmMWFmOGY0NzI1NyJ9.D6xt9mkS29qVGIWSKzxPBgfx1Kbyh5TwHj 9ISMebj9ZBl5kYg5gi3qm_nEwPsXcqfaVBYWqiEJI4VCwyme1dkw)].

## Author information

Author ORCIDs

Maheshi Dharmasiri https://orcid.org/0009-0009-2650-0648

Zoë Lindo https://orcid.org/0000-0001-9942-7204

Erin Cameron https://orcid.org/0000-0002-3374-9830

## Author contributions

### Funding information

This research was supported by Natural Sciences and Engineering Research Council of Canada (NSERC) (RGPIN-2019-05758 to EKC), a NSERC Postgraduate Scholarship – Doctoral (PGS D grant #601407) to MED and Saint Mary’s University Durland Scholarship in Graduate Research to MED.

## References

1. A’Bear, A.D., Jones, T.H., and Boddy, L. 2014. Potential impacts of climate change on interactions among saprotrophic cord-forming fungal mycelia and grazing soil invertebrates. Fungal Ecol. 10: 34–43. doi:10.1016/j.funeco.2013.01.009.

2. AdaptWest 2022. Gridded current and future climate data for North America at 1km resolution, interpolated using the ClimateNA v7.3 software. [Online] Available: adaptwest.databasin.org [2025 Oct. 2].

3. Agriculture and Agri-Food Canada 2017. A National Ecological Framework for Canada. [Online] Available: https://sis.agr.gc.ca/cansis/nsdb/ecostrat/gis_data.html [2025 Sept. 12].

4. Anthony, M.A., Bender, S.F., and Van Der Heijden, M.G.A. 2023. Enumerating soil biodiversity. Proc. Natl. Acad. Sci. 120: e2304663120. doi:10.1073/pnas.2304663120.

5. Arias-Navarro, C., Baritz, R., and Jones, A. (eds.) 2024. The state of soils in Europe: fully evidenced, spatially organised assessment of the pressures driving soil degradation. Publications Office of the European Union, Luxembourg. doi:10.2760/5897030.

6. Arieira, G. de O., Santiago, D.C., Franchini, J.C., and Guimarães, M. de F. 2016. Depth-stratified soil sampling for assessing nematode communities. Semina Ciênc. Agrár. 37: 715–728. doi:10.5433/1679-0359.2016v37n2p715.

7. Bardgett, R.D., and Van Der Putten, W.H. 2014. Belowground biodiversity and ecosystem functioning. Nature 515: 505–511. doi:10.1038/nature13855.

8. Beaumelle, L., Thouvenot, L., Hines, J., Jochum, M., Eisenhauer, N., and Phillips, H.R.P. 2021. Soil fauna diversity and chemical stressors: a review of knowledge gaps and roadmap for future research. Ecography 44: 845–859. doi:10.1111/ecog.05627.

9. Beaumelle, L., Tison, L., Eisenhauer, N., Hines, J., Malladi, S., Pelosi, C., Thouvenot, L., and Phillips, H.R.P. 2023. Pesticide effects on soil fauna communities—A meta-analysis. J. Appl. Ecol. 60: 1239–1253. doi:10.1111/1365-2664.14437.

10. Bicho, R.C., Scott-Fordsmand, J.J., and Amorim, M.J.B. 2025. Impacts of climate change–simulated flooding and drought events–on terrestrial invertebrates (Enchytraeids and Collembolans). Soil Ecol. Lett. 7: 240262. doi:10.1007/s42832-024-0262-2.

11. Bivand, R. 2025. classInt: Choose Univariate Class Intervals. doi:10.32614/CRAN.package.classInt.

12. Bongers, T. 1990. The maturity index: an ecological measure of environmental disturbance based on nematode species composition. Oecologia 83: 14–19. doi:10.1007/BF00324627.

13. Briones, M.J.I., Ostle, N.J., and Garnett, M.H. 2007. Invertebrates increase the sensitivity of non-labile soil carbon to climate change. Soil Biol. Biochem. 39: 816–818. doi:10.1016/j.soilbio.2006.09.007.

14. Bush, E., and Lemmen, D.S. 2019. Canada’s Changing Climate Report. Government of Canada, Ottawa, ON, Canada.

15. Cameron, E.K., Martins, I.S., Lavelle, P., Mathieu, J., Tedersoo, L., Bahram, M., Gottschall, F., Guerra, C.A., Hines, J., Patoine, G., Siebert, J., Winter, M., Cesarz, S., Ferlian, O., Kreft, H., Lovejoy, T.E., Montanarella, L., Orgiazzi, A., Pereira, H.M., Phillips, H.R.P., Settele, J., Wall, D.H., and Eisenhauer, N. 2019a. Global mismatches in aboveground and belowground biodiversity. Conserv. Biol. 33: 1187–1192. doi:10.1111/cobi.13311.

16. Cameron, E.K., Martins, I.S., Lavelle, P., Mathieu, J., Tedersoo, L., Gottschall, F., Guerra, C.A., Hines, J., Patoine, G., Siebert, J., Winter, M., Cesarz, S., Delgado-Baquerizo, M., Ferlian, O., Fierer, N., Kreft, H., Lovejoy, T.E., Montanarella, L., Orgiazzi, A., Pereira, H.M., Phillips, H.R.P., Settele, J., Wall, D.H., and Eisenhauer, N. 2018. Global gaps in soil biodiversity data. Nat. Ecol. Evol. 2: 1042–1043. doi:10.1038/s41559-018-0573-8.

17. Cameron, E.K., Sundqvist, M.K., Keith, S.A., CaraDonna, P.J., Mousing, E.A., Nilsson, K.A., Metcalfe, D.B., and Classen, A.T. 2019b. Uneven global distribution of food web studies under climate change. Ecosphere 10: e02645. doi:10.1002/ecs2.2645.

18. Canadian Centre for Climate Services 2020. Canadian statistically downscaled climate scenarios. Government. [Online] Available: https://climate-change.canada.ca/climate-data/#/downscaled-data [2025 Sept. 12].

19. Castle, S.E., Miller, D.C., Merten, N., Ordonez, P.J., and Baylis, K. 2022. Evidence for the impacts of agroforestry on ecosystem services and human well-being in high-income countries: a systematic map. Environ. Evid. 11. Springer Science and Business Media LLC. doi:10.1186/s13750-022-00260-4.

20. COP and CBD 2022a. Decision adopted by the Conference of the Parties to the Convention on Biological Diversity. 15/28. Biodiversity and Agriculture. [Online] Available: https://www.cbd.int/doc/decisions/cop-15/cop-15-dec-28-en.pdf.

21. COP and CBD 2022b. Decision Adopted by the Conference of the Parties to the Convention on Biological Diversity: 15/5. Monitoring framework for the Kunming-Montreal Global Biodiversity Framework. [Online] Available: https://www.cbd.int/doc/decisions/cop-15/cop-15-dec-05-en.pdf.

22. Council of Canadian Academies 2010. Canadian Taxonomy: Exploring Biodiversity, Creating Opportunity. The Expert Panel on Biodiversity Science. Ottawa, ON, Canada.

23. Decaëns, T., Jiménez, J.J., Gioia, C., Measey, G.J., and Lavelle, P. 2006. The values of soil animals for conservation biology. Eur. J. Soil Biol. 42: S23–S38. doi:10.1016/j.ejsobi.2006.07.001.

24. Deguise, I.E., and Kerr, J.T. 2006. Protected Areas and Prospects for Endangered Species Conservation in Canada. Conserv. Biol. 20: 48–55. doi:10.1111/j.1523-1739.2005.00274.x.

25. ECCC 2024. Canada’s 2030 Nature Strategy: Halting and Reversing Biodiversity Loss in Canada. Quebec, Canada. [Online] Available: https://publications.gc.ca/collections/collection_2024/eccc/en4/En4-539-1-2024-eng.pdf.

26. Ecological Stratification Working Group 1995. A National Ecological Framework for Canada. Agriculture and Agri-Food Canada, Research Branch, Centre for Land and Biological Resources Research and Environment Canada, State of the Environment Directorate, Ecozone Analysis, Branch, Ottawa/Hull.

27. Eisenhauer, N., Frank, K., Weigelt, A., Bartkowski, B., Beugnon, R., Liebal, K., Mahecha, M., Quaas, M., Al-Halbouni, D., Bastos, A., Bohn, F.J., De Brito, M.M., Denzler, J., Feilhauer, H., Fischer, R., Fritsche, I., Guimaraes-Steinicke, C., Hänsel, M., Haun, D.B.M., Herrmann, H., Huth, A., Kalesse-Los, H., Koetter, M., Kolleck, N., Krause, M., Kretschmer, M., Leitão, P.J., Masson, T., Mora, K., Müller, B., Peng, J., Pöhlker, M.L., Ratzke, L., Reichstein, M., Richter, S., Rüger, N., Sánchez-Parra, B., Shadaydeh, M., Sippel, S., Tegen, I., Thrän, D., Umlauft, J., Wendisch, M., Wolf, K., Wirth, C., Zacher, H., Zaehle, S., and Quaas, J. 2024. A belowground perspective on the nexus between biodiversity change, climate change, and human well-being. J. Sustain. Agric. Environ. 3: e212108. Wiley. doi:10.1002/sae2.12108.

28. Esri 2024. ArcGIS Pro. Redlands, California, USA.

29. Ettema, C. 2002. Spatial soil ecology. Trends Ecol. Evol. 17: 177–183. doi:10.1016/S0169-5347(02)02496-5.

30. FAO, ITPS, GSBI, and EC 2020. State of knowledge of soil biodiversity - Status, challenges and potentialities. FAO, Rome, Italy. doi:10.4060/cb1928en.

31. Forge, T., Munro, P., Wright, H., and Moreau, D. 2022. Plant-parasitic nematodes in Nova Scotia vineyards. Phytoprotection 102: 15. doi:10.7202/1088485ar.

32. Fyson, V.K., Wilmshurst, J.F., and Callaghan, C. 2024a. The changing agricultural landscape in Canada’s Mixedwood Plains Ecozone (2011–2022) and the implications for biodiversity. FACETS 9: 1–11. doi:10.1139/facets-2024-0019.

33. Fyson, V.K., Wilmshurst, J.F., and Callaghan, C. 2024b. The changing agricultural landscape in Canada’s Mixedwood Plains Ecozone (2011–2022) and the implications for biodiversity. FACETS 9: 1–11. doi:10.1139/facets-2024-0019.

34. Geldmann, J., Heilmann-Clausen, J., Holm, T.E., Levinsky, I., Markussen, B., Olsen, K., Rahbek, C., and Tøttrup, A.P. 2016. What determines spatial bias in citizen science? Exploring four recording schemes with different proficiency requirements. Divers. Distrib. 22: 1139–1149. doi:10.1111/ddi.12477.

35. Goncharov, A.A., Leonov, V.D., Rozanova, O.L., Semenina, E.E., Tsurikov, S.M., Uvarov, A.V., Zuev, A.G., and Tiunov, A.V. 2023. A meta-analysis suggests climate change shifts structure of regional communities of soil invertebrates. Soil Biol. Biochem. 181: 109014. doi:10.1016/j.soilbio.2023.109014.

36. Gonzalez, A., O’Connor, M.I., Bates, A.E., Bobiwash, K., Burton, A.C., Van Dam-Bates, P., Eckert, I., Gravel, D., Idrobo, C.J., Pollock, L., Simon, A.D.F., Slein, M.A., Sólymos, P., Starzomski, B.M., Sunday, J., and Tekwa, E. 2025. A Biodiversity Observation Network to support conservation action and mainstream knowledge in Canada. FACETS 10: 1–19. doi:10.1139/facets-2024-0206.

37. Goulet, H. 2003. Biodiversity of ground beetles (Coleoptera: Carabidae) in Canadian agricultural soils. Can. J. Soil Sci. 83: 259–264. doi:10.4141/S01-061.

38. Government of Canada 2017.May 3. GIS Data. [Online] Available: https://sis.agr.gc.ca/cansis/nsdb/ecostrat/gis_data.html [2025 Sept. 12].

39. Gruss, I., Czarniecka-Wiera, M., Świerszcz, S., Szymura, M., Szymura, T., and Raduła, M.W. 2025. Responses of grassland soil mesofauna to induced climate change. Sci. Rep. 15: 16532. doi:10.1038/s41598-025-01445-w.

40. Guerra, C.A., Bardgett, R.D., Caon, L., Crowther, T.W., Delgado-Baquerizo, M., Montanarella, L., Navarro, L.M., Orgiazzi, A., Singh, B.K., Tedersoo, L., Vargas-Rojas, R., Briones, M.J.I., Buscot, F., Cameron, E.K., Cesarz, S., Chatzinotas, A., Cowan, D.A., Djukic, I., Van Den Hoogen, J., Lehmann, A., Maestre, F.T., Marín, C., Reitz, T., Rillig, M.C., Smith, L.C., De Vries, F.T., Weigelt, A., Wall, D.H., and Eisenhauer, N. 2021. Tracking, targeting, and conserving soil biodiversity. Science 371: 239–241. doi:10.1126/science.abd7926.

41. Guerra, C.A., Heintz-Buschart, A., Sikorski, J., Chatzinotas, A., Guerrero-Ramírez, N., Cesarz, S., Beaumelle, L., Rillig, M.C., Maestre, F.T., Delgado-Baquerizo, M., Buscot, F., Overmann, J., Patoine, G., Phillips, H.R.P., Winter, M., Wubet, T., Küsel, K., Bardgett, R.D., Cameron, E.K., Cowan, D., Grebenc, T., Marín, C., Orgiazzi, A., Singh, B.K., Wall, D.H., and Eisenhauer, N. 2020a. Blind spots in global soil biodiversity and ecosystem function research. Nat. Commun. 11: 3870. doi:10.1038/s41467-020-17688-2.

42. Guerra, C.A., Heintz-Buschart, A., Sikorski, J., Chatzinotas, A., Guerrero-Ramírez, N., Cesarz, S., Beaumelle, L., Rillig, M.C., Maestre, F.T., Delgado-Baquerizo, M., Buscot, F., Overmann, J., Patoine, G., Phillips, H.R.P., Winter, M., Wubet, T., Küsel, K., Bardgett, R.D., Cameron, E.K., Cowan, D., Grebenc, T., Marín, C., Orgiazzi, A., Singh, B.K., Wall, D.H., and Eisenhauer, N. 2020b. Blind spots in global soil biodiversity and ecosystem function research. Nat. Commun. 11. Springer Science and Business Media LLC. doi:10.1038/s41467-020-17688-2.

43. Haddaway, N.R. 2019.Evidence of the impacts of metal mining and the effectiveness of mining mitigation measures on social–ecological systems in Arctic and boreal regions: a systematic map protocol.

44. Haddaway, N.R., Bernes, C., Jonsson, B.-G., and Hedlund, K. 2016. The benefits of systematic mapping to evidence-based environmental management. Ambio 45: 613–620. Springer Science and Business Media LLC. doi:10.1007/s13280-016-0773-x.

45. Haddaway, N.R., Macura, B., and Whaley, P. 2017. ROSES for Systematic Map Protocols.

46. Hijmans, R.J. 2025. terra: Spatial Data Analysis. doi:10.32614/CRAN.package.terra.

47. Jorna, J., Vandenbrink, B., Hogg, I.D., Wall, D.H., and Adams, B.J. 2023. Metabarcoding inventory of an arctic tundra soil ecosystem reveals highly heterogeneous communities at a small scale. Polar Biol. 46: 461–471. doi:10.1007/s00300-023-03131-x.

48. Keppel, G., Stralberg, D., Morelli, T.L., and Bátori, Z. 2024. Managing climate-change refugia to prevent extinctions. Trends Ecol. Evol. 39: 800–808. doi:10.1016/j.tree.2024.05.002.

49. Köninger, J., Panagos, P., Jones, A., Briones, M.J.I., and Orgiazzi, A. 2022. In defence of soil biodiversity: Towards an inclusive protection in the European Union. Biol. Conserv. 268: 109475. Elsevier BV. doi:10.1016/j.biocon.2022.109475.

50. Kotschik, P., Princz, J., Silva, C.D.L.E., Renaud, M., Marti-Roura, M., Brooks, B., Pieper, S., Rijk, I., Simini, M., Andres, S., Scholz-Starke, B., and Grenni, P. 2024. The upcoming European Soil Monitoring Law: An effective instrument for the protection of terrestrial ecosystems? Integr. Environ. Assess. Manag. 20: 316–321. Oxford University Press (OUP). doi:10.1002/ieam.4834.

51. Lavelle, P., Decaëns, T., Aubert, M., Barot, S., Blouin, M., Bureau, F., Margerie, P., Mora, P., and Rossi, J.-P. 2006. Soil invertebrates and ecosystem services. Eur. J. Soil Biol. 42: S3–S15. Elsevier BV. doi:10.1016/j.ejsobi.2006.10.002.

52. Li, H. 2024. What evidence exists regarding the impact of biodiversity on human health and well-being? A systematic map protocol.

53. Li, Y., Wang, J., and Shao, M. 2021. Assessment of earthworms as an indicator of soil degradation: A case-study on loess soils. Land Degrad. Dev. 32: 2606–2617. doi:10.1002/ldr.3928.

54. Llopis, J.C., Haddaway, N.R., Omirbek, N., Simmons, B.A., Garrett, R., and Jones, J.P.G. 2023. Evidence of anticipatory forest use behaviours under policy introduction: a systematic map protocol. Environ. Evid. 12: 20. doi:10.1186/s13750-023-00307-0.

55. Madani, M., Tenuta, M., and Castillo, P. 2024. The occurrence of cyst nematodes in agricultural fields of Southern Manitoba. Can. J. Plant Pathol. 46: 306–317. doi:10.1080/07060661.2023.2293128.

56. Manu, M., Honciuc, V., Neagoe, A., Băncilă, R.I., Iordache, V., and Onete, M. 2019. Soil mite communities (Acari: Mesostigmata, Oribatida) as bioindicators for environmental conditions from polluted soils. Sci. Rep. 9: 20250. doi:10.1038/s41598-019-56700-8.

57. Martin, P.A., Izquierdo, L.P., Biryol, C., Brasseur, S., Menival, C., Li, Q., Aviles, I.L., Petritan, I.C., Luyssaert, S., Geunet, B., Manzoni, S., Spake, R., Santonja, M., and Curiel, J. 2025. Forest disturbance impacts on soil and litter fauna: knowledge gaps and a roadmap for future research. Ecoevorxiv. [Online] Available: 10.32942/X2R929.

58. Mathieu, J., Antunes, A.C., Barot, S., Bonato Asato, A.E., Bartz, M.L.C., Brown, G.G., Calderon-Sanou, I., Decaëns, T., Fonte, S.J., Ganault, P., Gauzens, B., Gongalsky, K.B., Guerra, C.A., Hengl, T., Lavelle, P., Marichal, R., Mehring, H., Peña-Venegas, C.P., Castro, D., Potapov, A., Thébault, E., Thuiller, W., Witjes, M., Zhang, C., and Eisenhauer, N. 2022. sOilFauna - a global synthesis effort on the drivers of soil macrofauna communities and functioning: WORKSHOP REPORT. SOIL ORGANISMS 94(2). doi:10.25674/SO94ISS2ID282.

59. McFarlane, S., Van Mierlo, V., Manseau, M., Kroeze, A., Eberhardt, E., and Girard, J. 2025. Bioclimatic, terrain, and specific peatland composition are major drivers of woodland caribou winter habitat suitability in northern Ontario. Can. J. Zool. 103: 1–18. doi:10.1139/cjz-2024-0121.

60. Minister of Supply and Services Canada (ed.) 1995. Canadian biodiversity strategy: Canada’s response to the Convention on Biological Diversity. Biodiversity Convention Office, Environment Canada, Hull, Quebec, Canada.

61. Montanarella, L. 2008. Towards protecting soil biodiversity in Europe: The EU thematic strategy for soil protection. Biodiversity 9: 75–77. doi:10.1080/14888386.2008.9712889.

62. Morelli, T.L., Mozelewski, T., Cavalieri, C.N., Caven, A.J., Dreiss, L.M., Hovel, R.A., Hua, M., Jennings, M.K., John, A., Kehm, G., Keppel, G., Krawchuk, M.A., Langdon, S.F., Lawler, J.J., Lyon, L.M., Meigs, G.W., Mora-Gonzalez, M., Nadeau, C.P., Słowińska, S., Stolar, J., Thorne, J.H., Tourville, J.C., Zuckerberg, B., and Stralberg, D. 2026. Conserving climate-change refugia: Insights from research and practice. Conserv. Sci. Pract. 8: e70160. doi:10.1111/csp2.70160.

63. Orgiazzi, A., Ballabio, C., Panagos, P., Jones, A., and Fernández-Ugalde, O. 2018. LUCAS Soil, the largest expandable soil dataset for Europe: a review. Eur. J. Soil Sci. 69: 140–153. doi:10.1111/ejss.12499.

64. Orgiazzi, A., Bardgett, R.D., Barrios, E., Behan-Pelletier, V., Briones, M.J.I., Chotte, J.-L., De Deyn, G.B., Eggleton, P., Fierer, N., Fraser, T., Hedlund, K., Jeffery, S., Johnson, N.C., Jones, A., Kandeler, E., Kaneko, N., Lavelle, P., Lemanceau, P., Miko, L., Montanarella, L., Moreira, F.M.S., Ramirez, K.S., Scheu, S., Singh, B.K., Six, J., Putten, W.H. van der, Wall, D.H., and European Commission (eds.) 2016. Global soil biodiversity atlas. Publications Office, Luxembourg. doi:10.2788/799182.

65. Oswald, A.M., Mannion, N., Willis, S.G., Stephens, P.A., and McGowan, P.J.K. 2025. What is favourable conservation status?: A systematic map protocol.

66. Ouzzani, M., Hammady, H., Fedorowicz, Z., and Elmagarmid, A. 2016. Rayyan—a web and mobile app for systematic reviews. Syst. Rev. 5: 210. doi:10.1186/s13643-016-0384-4.

67. Panagos, P., Jones, A., Lugato, E., and Ballabio, C. 2025. A Soil Monitoring Law for Europe. Glob. Chall. 9: 2400336. doi:10.1002/gch2.202400336.

68. Parnell, J., Brown, G., deRuiter, P., Guerra, C., Montanarella, L., Lindo, Z., Bagnara, G.L., Battigelli, J., González, R., Poch, R.M., Wall, D.H., Malorgio, G., Vargas, R., Cuevas-Corona, R., and Rodriguez Eugenio, N. 2025. GLOSOB: The Global Soil Biodiversity Observatory. Soil Org. 97: 143–148. Soil Organisms. doi:10.25674/426.

69. Pascual, U., Termansen, M., Hedlund, K., Brussaard, L., Faber, J.H., Foudi, S., Lemanceau, P., and Jørgensen, S.L. 2015. On the value of soil biodiversity and ecosystem services. Ecosyst. Serv. 15: 11–18. Elsevier BV. doi:10.1016/j.ecoser.2015.06.002.

70. Pebesma, E. 2018. Simple Features for R: Standardized Support for Spatial Vector Data. R J. 10: 439. doi:10.32614/RJ-2018-009.

71. Pebesma, E., and Bivand, R. 2023. Spatial Data Science: With Applications in R. Chapman and Hall/CRC, New York. doi:10.1201/9780429459016.

72. Pérès, G., Vandenbulcke, F., Guernion, M., Hedde, M., Beguiristain, T., Douay, F., Houot, S., Piron, D., Richard, A., Bispo, A., Grand, C., Galsomies, L., and Cluzeau, D. 2011. Earthworm indicators as tools for soil monitoring, characterization and risk assessment. An example from the national Bioindicator programme (France). Pedobiologia 54: S77–S87. doi:10.1016/j.pedobi.2011.09.015.

73. Phillips, H., Beaumelle, L., Tyndall, K., Burton, V., Cameron, E., Eisenhauer, N., and Ferlian, O. 2019. The effects of global change on soil faunal communities: a meta-analytic approach. Res. Ideas Outcomes 5: e36427. doi:10.3897/rio.5.e36427.

74. Phillips, H.R.P., Beaumelle, L., Eisenhauer, N., Hines, J., and Smith, L.C. 2020. Lessons from the WBF2020: extrinsic and intrinsic value of soil organisms. SOIL ORGANISMS 92(2). doi:10.25674/SO92ISS2PP121.

75. Phillips, H.R.P., Cameron, E.K., Eisenhauer, N., Burton, V.J., Ferlian, O., Jin, Y., Kanabar, S., Malladi, S., Murphy, R.E., Peter, A., Petrocelli, I., Ristok, C., Tyndall, K., Van Der Putten, W., and Beaumelle, L. 2024a. Global changes and their environmental stressors have a significant impact on soil biodiversity—A meta-analysis. iScience 27: 110540. Elsevier BV. doi:10.1016/j.isci.2024.110540.

76. Phillips, H.R.P., Cameron, E.K., Eisenhauer, N., Burton, V.J., Ferlian, O., Jin, Y., Kanabar, S., Malladi, S., Murphy, R.E., Peter, A., Petrocelli, I., Ristok, C., Tyndall, K., Van Der Putten, W., and Beaumelle, L. 2024b. Global changes and their environmental stressors have a significant impact on soil biodiversity—A meta-analysis. iScience 27: 110540. doi:10.1016/j.isci.2024.110540.

77. Ponge, J.F., Gillet, S., Dubs, F., Fedoroff, E., Haese, L., Sousa, J.P., and Lavelle, P. 2003. Collembolan communities as bioindicators of land use intensification. Soil Biol. Biochem. 35: 813–826. doi:10.1016/S0038-0717(03)00108-1.

78. Posit Software 2025. RStudio.

79. Potter, J.W., and McKeown, A.W. 2003. Nematode biodiversity in Canadian agricultural soils. Can. J. Soil Sci. 83: 289–302. doi:10.4141/S01-064.

80. R Core Team 2025. R: A Language and Environment for Statistical Computing. R Foundation for Statistical Computing, Vienna, Austria. [Online] Available: https://www.R-project.org/.

81. Ray, J.C., Grimm, J., and Olive, A. 2021. The biodiversity crisis in Canada: failures and challenges of federal and sub-national strategic and legal frameworks. FACETS 6: 1044–1068. Canadian Science Publishing. doi:10.1139/facets-2020-0075.

82. Ristok, C., Babin, D., Bartkowski, B., Burkhard, B., Filser, J., Hohberg, K., Kleemann, J., Kolb, S., Lehmitz, R., Rillig, M.C., Römbke, J., Ruess, L., Scheu, S., Scheunemann, N., Schmidt, A., Steinhoff-Knopp, B., Eisenhauer, N., Tebbe, C.C., and Xylander, W.E.R. 2025. Towards a comprehensive assessment of soil biodiversity in Germany: status quo, challenges, and policy implications.

83. Senate Canada 2024. Critical Ground - Why Soil is Essential to Canada’s Economic, Environmental, Human, and Social Health. Ottawa.

84. Smith, P., Poch, R.M., Lobb, D.A., Bhattacharyya, R., Alloush, G., Eudoxie, G.D., Anjos, L.H.C., Castellano, M., Ndzana, G.M., Chenu, C., Naidu, R., Vijayanathan, J., Muscolo, A.M., Studdert, G.A., Eugenio, N.R., Calzolari, M.C., Amuri, N., and Hallett, P. 2024. Status of the World’s Soils. Annu. Rev. Environ. Resour. 49: 73–104. doi:10.1146/annurev-environ-030323-075629.

85. Sobral-Souza, T., Bosco, N.S., Candelária, L.P., Collevatti, R.G., Layme, V.M.G., and Rodrigues, D.D.J. 2024. Spatial bias in sampling small rodents in the Atlantic Forest: A landscape and accessibility perspective. Perspect. Ecol. Conserv. 22: 297–305. Elsevier BV. doi:10.1016/j.pecon.2024.07.004.

86. Statistics Canada 2023. 2021 Census – Boundary files. [Online] Available: https://www12.statcan.gc.ca/census-recensement/2021/geo/sip-pis/boundary-limites/index2021-eng.cfm?year=21 [2026 Mar. 3].

87. Statistics Canada, Geography Division 2018. Geography. [Online] Available: https://www150.statcan.gc.ca/n1/pub/11-402-x/2011000/chap/geo/geo-eng.htm.

88. Stevenson, M., and Sergeant, E. 2025. epiR: Tools for the Analysis of Epidemiological Data. doi:10.32614/CRAN.package.epiR.

89. Subbotin, S.A., and Chitambar, J.J. (eds.) 2018. Plant Parasitic Nematodes in Sustainable Agriculture of North America: Vol.1 - Canada, Mexico and Western USA. Springer International Publishing, Cham. doi:10.1007/978-3-319-99585-4.

90. Thakur, M.P., Phillips, H.R.P., Brose, U., De Vries, F.T., Lavelle, P., Loreau, M., Mathieu, J., Mulder, C., Van Der Putten, W.H., Rillig, M.C., Wardle, D.A., Bach, E.M., Bartz, M.L.C., Bennett, J.M., Briones, M.J.I., Brown, G., Decaëns, T., Eisenhauer, N., Ferlian, O., Guerra, C.A., König-Ries, B., Orgiazzi, A., Ramirez, K.S., Russell, D.J., Rutgers, M., Wall, D.H., and Cameron, E.K. 2020. Towards an integrative understanding of soil biodiversity. Biol. Rev. 95: 350–364. Wiley. doi:10.1111/brv.12567.

91. Turbé, A., Toni, A.D., Benito, P., Lavelle, P., Lavelle, P., Ruiz, N., Putten, W.H.V. der, Labouze, E., and Mudgal, S. 2010. Soil biodiversity: functions, threats and tools for policy makers. Bio Intelligence Service, IRD, and NIOO, Report for European Commission (DG Environment).

92. Van Vuuren, D.P., Edmonds, J., Kainuma, M., Riahi, K., Thomson, A., Hibbard, K., Hurtt, G.C., Kram, T., Krey, V., Lamarque, J.-F., Masui, T., Meinshausen, M., Nakicenovic, N., Smith, S.J., and Rose, S.K. 2011. The representative concentration pathways: an overview. Clim. Change 109: 5–31. doi:10.1007/s10584-011-0148-z.

93. Wagg, C., Bender, S.F., Widmer, F., and Van Der Heijden, M.G.A. 2014. Soil biodiversity and soil community composition determine ecosystem multifunctionality. Proc. Natl. Acad. Sci. 111: 5266–5270. Proceedings of the National Academy of Sciences. doi:10.1073/pnas.1320054111.

94. Wang, T., Hamann, A., Spittlehouse, D., and Carroll, C. 2016. Locally Downscaled and Spatially Customizable Climate Data for Historical and Future Periods for North America. PLOS ONE 11: e0156720. doi:10.1371/journal.pone.0156720.

95. Wang, T., Hamann, A., Spittlehouse, D., and Carroll, C. 2022. Gridded current and future climate data for North America at 1km resolution, interpolated using the ClimateNA v7.3 software. AdaptWest. [Online] Available: adaptwest.databasin.org [2025 Oct. 2].

96. Wotherspoon, A.R., Achim, A., and Coops, N.C. 2024. Assessing future climate trends and implications for managed forests across Canadian ecozones. Can. J. For. Res. 54: 278–289. doi:10.1139/cjfr-2023-0058.

97. Wu, D., Du, E., Eisenhauer, N., Mathieu, J., and Chu, C. 2025. Global engineering effects of soil invertebrates on ecosystem functions. Nature 640: 120–129. doi:10.1038/s41586-025-08594-y.

98. Wu, T., Ayres, E., Bardgett, R.D., Wall, D.H., and Garey, J.R. 2011. Molecular study of worldwide distribution and diversity of soil animals. Proc. Natl. Acad. Sci. 108: 17720–17725. doi:10.1073/pnas.1103824108.

99. Yang, J., Wu, X., Chen, Y., Yang, Z., Liu, J., Wu, D., and Wang, D. 2021. Combined attributes of soil nematode communities as indicators of grassland degradation. Ecol. Indic. 131: 108215. doi:10.1016/j.ecolind.2021.108215.

100. Zeng, X.-M., Delgado-Baquerizo, M., Wen, S., Feng, J., Zhang, W., Zhang, Q., and Liu, Y.-R. 2025. Soil invertebrate diversity supports ecosystem multifunctionality along elevation gradients. Plant Soil 512: 1243–1260. doi:10.1007/s11104-024-07142-3.

