## supplementary files for "A systematic map of soil invertebrate biodiversity in Canada: Identifying spatial and taxonomic knowledge gaps"

S1: ROSES for Systematic Map Protocol (*included separately)*

S2: ROSES flow diagram (*included separately)*

S3: Abstract and title screening protocol

S4: Full-text data extraction protocol

S5: Ecozone chi-squared table

S6: Current mean annual climate period chi-squared table

S7: Projected mean annual climate chi-squared table

S8: Sampled locations in relation to the soil types in Canada

S9: References for articles included in the study

**S3: Abstract and title screening protocol**

**Inclusion Criteria**
For inclusion at this stage, the article should contain data on soil biodiversity AND have taken place in Canada.

Soil Biodiversity Data

- Include data on soil biodiversity, including diversity, abundance, richness, or biomass data.

Geographic Location

- Include a study that was conducted in Canada. However, if no geographic location is provided but the study reports soil biodiversity data then include it. The geographic location can be checked at the full text screening stage.

Methodologies

Include any of the following:

- Soil/fen cores samples
- Peat cores
- Litter bags, either on the surface or belowground
- Hand sorting litter
- Digging or plowing
- Sampling related to the rhizosphere (e.g., mycorrhizal fungi)
- A combination of endosphere and rhizosphere sampling
- A study that uses pitfall traps in combination with other methods to determine biodiversity/abundance/relative abundance
- Emergence cages
- Lab reared organisms Microorganisms or soil invertebrates that were manipulated/inoculated/added to artificial/autoclaved/sterilized environments,
- If the soil sampling methodology is not explained, include the study for full text screening.

We are excluding laboratory reared the soil organisms, store bought organisms and any study that only uses pitfall traps to sample organisms. Buried wood soil for colonization over time are not included.

Study Type

- Include a study (e.g., field, laboratory, mesocosm) that involves soil organisms, soil invertebrates, and/or soil microorganisms. We are excluding Paleoecology studies, species description papers and review papers.

Soil Life Stages

- Include any organism that completes one or more stages of their life cycle in soil.

Habitat

Include any of the following habitats:

- Grassland
- Agriculture
- Woody
- Cold/dry system
- Wetland
- Tree plantation
- Anthropogenic disturbed sites
- Coastal
- Urban

We are excluding aquatic habitats, aquifers (underground water sources), and intertidal zones.

**S4: Full-text data extraction protocol**

For the study to be appropriate for data extraction it should have clear location details, data on soil organisms, and an appropriate soil sampling methodology. To determine if the study is suitable for data extraction, read the objectives, aims, and/or hypothesis paragraph(s) in the study.

1. Identify the location of sampling. If it was conducted outside of Canada, it can be excluded.
2. Data on soil biodiversity at the taxa level should be provided either as diversity, richness, abundance, or a list of organisms from which diversity can be calculated. This information may not always be provided in the body of the article but as supplementary material. This should be checked before excluding at the full text stage.
3. Identify if an appropriate soil sampling methodology is used.

If these criteria are met proceed to data extraction.

**Data Extraction**

1. **Methods**

- All the methodologies listed below are included. This is a more detailed list to assist with inclusion/exclusion at the full text screening stage.
  - Any type of soil core including fen, peat, or litter. The core can be of any size.
  - Hand sorting, sifting soil/litter and or digging for soil samples without a soil corer.
  - Litter bags or mesh bags, which can either be on the surface or buried below-ground.
  - Any form of rhizosphere sampling. The soil must still be adhered to roots (e.g. root zone, closely bound to roots). Exclude the study, if the soil is washed off before measuring a variable.
  - Pitfall traps in combination with other methods to determine biodiversity/abundance. This means a single taxon must be sampled with both pitfall trapping and soil cores (or another accepted method). Exclude studies that use pitfall traps as their only method of determining biodiversity/abundance.
  - Emergence cages/traps
  - Ants or termites sampled from termite mounds or ant hills
  - Source population information for microbes or soil invertebrates must be provided for studies that use exposed/inoculated/added to artificial/autoclaved/sterilized**.**
  - For mesocosm/microcosm/greenhouse/laboratory experiment studies, control or time at zero soil community profiles/abundances must be provided to reflect the original community sampled.
  - Any study that simply stated collection/sampling, but does not explicitly state or describe an accepted soil sampling methodology, then only include it if the soil organism’s full life cycle takes place in soil.

2. **Exclusion Justification**

An article excluded at the full text stage requires a reason.

- Select one of the following exclusion reasons. If the reason does not fit the following, then provide a justification.
  - Not in Canada
  - Lab cultures/store bought
  - No sampling information – This is when no information is provided on how an organism was collected from their environment.
  - No soil biodiversity data – This is when a study reports no diversity, abundance, richness, or soil organism data or if the reported data are at a coarse level (e.g., soil microbes, soil fauna etc.). This category can also include studies reporting below-ground physical or chemical properties.
  - Sampling methodology not accepted – A study that may report soil organism data but were sampled using alternative methods not accepted in this systematic map (i.e. sticky traps, Bovian traps, etc.)
  - Pitfall trap only – A study that used pitfall traps only to sample soil organisms.
  - Paleoecology studies
  - Language – If the study is not in English
  - Habitat not accepted – A study conducted above-ground or below-ground but not specifically from the soil (i.e., roots only, garlic bulbs, carcasses, dung pats, marine or aquatic environments).

3. **Meta Data Extraction**

Sampling time-frame column

- The timeframe for each study was broken down into two groups; as either a one-year study or a multi-year study. For a study reporting more than one taxon, each taxon should be considered individually for the timeframe. If no information on when the fieldwork/data collection was reported (i.e. no year, month) then indicate not provided.

Study type:

- Reading the objectives of the study will help clarify the type of study being conducted (Field, Laboratory, Mesocosm/Microcosm/Greenhouse, Modeling). A study can have multiple types as there may be a field and laboratory component that included soil organisms.

Taxa identification method

- The type of method used for organism identification (morphological, molecular, biochemical). A study can use a combination of these methods.

Extraction Type

- This is applicable to soil invertebrates. This is intended to capture the methodology used to extract soil invertebrates from soil. A study can use more than one of these forms.
  - Wet: Oostenbrink elutriators, Baermann funnels, Baermann pan, O’Connor wet-funnel method, Townshend wet method, mist chamber method, cold/heat wet funnel extraction, Cobb’s decanting and sieving method, wet and dry extraction
  - Heat: Berlese funnels, Tullgren funnels, MacFadyen extractors, Berlese-Tullgren funnels, high gradient extraction, Lussenhop extractor, Merchant-Crossley extractor, modified (Hill 1969) Kempson, Lloyd & Gelhardi (1963) apparatus, a high efficiency canister-type soil arthropod extractor, wet and dry extraction
  - Chemical: 4% formalin, 0.5% formaldehyde solution, mustard solution extraction/hot mustard, allyl isothiocyanate (AITC) solution extraction, allium cepa
  - Not Provided: A study does not report any form of extraction.

Methods open comments

- Write the specific type of sampling that was used to obtain or extract other soil organisms that do not fit the above-mentioned categories. This includes hand sorting (includes hand collected, hand sifting, hand sorted from under logs), flotation, centrifugation, litterbags, or emergence cages.

Variable measured

- A study can report more than one variable measured, in which case the reviewer should select all applicable options.
  - Abundance: Include any numerical count such as density, absolute abundance, relative abundance, etc. For molecular studies, only report abundance when qPCR/RT- qPCR or ddPCR is used.
  - Taxonomic diversity metrices: Reports taxonomic diversity indices (e.g., Shannon, Simpsons, etc.) or species richness (or at any other taxonomic level). This is a measure of the number of different species at a specific site and their evenness. This number can be provided at other levels such as genera, orders, families etc. If species richness is not provided but occurrence data is provided, then check if species richness can be calculated. A study that provides different types of organisms found at the site (occurrence data) can be used to determine richness.
  - Occurrence: This is when no abundance, diversity, or biomass information is provided but a location is provided showing where the soil organism was found.
  - Biomass: All forms of biomass are included (e.g., fresh biomass, dry biomass, ash-free dry biomass). If biomass is calculated for that taxon, indicate it by selecting this option as well.

Taxonomic resolution

- Record the level each soil organism was identified to in the study (e.g., species, genus, family, etc.). Taxa reported as ‘earthworm,’ or ‘mites’, etc. should be reported as NA for this column.

Life stage

- A study that reports data on different life stages must be recorded here (e.g., juveniles, cocoons, larvae, pupae, etc.)

Pooled Data

The data will be considered pooled if following conditions apply:

- A study that does not distinguish between methods used to sample soil organisms (e.g., soil cores and aerial sampling) and one of the methods used is not an accepted soil methodology (e.g. aerial sampling).
- The sampled countries (e.g., Canada and United States) do not distinguish the soil organisms found from each country but reports the pooled data in the analyses.
- The sampled habitats provide a combination of habitats some of which are not accepted but only the pooled data is reported (e.g., soil habitat and moss).

4. **Location and Habitat Extraction**

**Location**

- Enter all sampled site locations. If the study provides precise GPS coordinates for the sampling locations, then enter these instead of coordinates provided for the general sample site.
- For a study that does not report GPS coordinates, read the description of the sampled site and use google maps to extract GPS coordinates that match this location description. Make a note that these are vague locations.

**Habitat**

If the article provides only one GPS location but provides two different types of habitats, then select both types of habitats.

a. **Agriculture:** Orchards (e.g., cherry, apple), cornfields, cultivated land, tobacco
b. **Woody:** Coniferous, deciduous, mixed forest, shrubs, reclaimed sites that are now planted with woody trees (not for forestry/harvesting), woody hedgerows, sampling of woody sites near roadsides/campsites (make a note indicating adjacent/near roadsides or campsites
c. **Grassland:** Grazed, managed, mown, mixture, meadows, sward
d. **Cold Dry System:** Alpine, tundra, desert, drylands
e. **Wetland:** Peatland, saltmarsh, swamps, inland ponds sites
f. **Tree Plantation:** This can be for any forestry purpose, e.g., timber plantation, tree farm, forest plantation, groves (if these trees are for harvesting fruit, nuts, etc. then select Agriculture option)
g. **Anthropogenic Disturbance Sites:** Mining waste sites, mining sites that are not reclaimed for any other purpose, sour gas plants, landfills, clear cuts areas
h. **Urban:** City centers, city parks, car parks, recreational areas, railroads, golf course, gardens, campground, near highways (rocky outcrops/moss carpets), roadsides
i. **Coastal:** Beaches, riparian zones, riverbanks
j. **Other:** Any other habitat not fitting the above-mentioned habitat description. Examples include weeds, grass with weeds, infested fields.

**Data Extracted**

This is the information that was extracted into the data sheet and the data extracted.

| Column Name | Description |
| --- | --- |
| ID | Each study obtained from the search was assigned a unique ID (numeric) |
| Title | Title of article |
| Year | Year the article was published |
| Journal | The journal the article was published in |
| Issue | Journal issue |
| Pages | Article pages |
| Authors | Last name, First name initial of all authors of the article |
| DOI | DOI of the article |
| Reviewer | The researcher who reviewed the article at the full-text screen stage |
| Excluded full text | No = Full text was included; Yes = full text was excluded |
| Excluded full text reason | The reason why the article was excluded at the full-text stage. |
| Sampling timeframe | The duration of the study |
| Sampling time frame comments | Additional comments about sampling time frame |
| Study type | The type of study conducted |
| Taxa identification method | The method in which the taxa were identified |
| Extraction type | The method soil organisms were extracted from soil |
| Methods Comments | Additional methodologies used to sample soil invertebrates. |
| Variable measured | The types of soil organism data reported |
| Taxonomic resolution | The taxonomic extent the soil organism was identified to |
| Soil organism | The soil organism(s) studied |
| Harmonised name | The names for each taxon were standardized |
| Body size | The size was assigned according to the GSBA |
| Invertebrate | No = taxon is not an invertebrate; Yes = taxon is an invertebrate |
| Taxonomic comments | Additional comments about the taxa |
| Insect | No = taxon is not an insect; Yes = taxon is an insect |
| Life stage | What life stages were reported? |
| Pooled data | No = data was not pooled; Yes = data was pooled |
| Province/territory | The province or territory the study was conducted in |
| Location | The city/town/county/geographical region the study was conducted in |
| Location coordinates provided | No = coordinates not given in article; Yes = coordinates given in article |
| GPS coordinates | The latitude and longitude of the study site |
| Habitat | The environment the study was conducted in |
| Location comments | Additional comments about the location |

**S5: Ecozone chi-squared table**

**Table S6.** The number of soil biodiversity sampled locations obtained from the literature for each of the 15 terrestrial ecozones in Canada, and the expected number of sampled locations based on geographic area for each ecozone assuming a search effort proportional to the ecozone area.

| Ecozone | Area | Observed | Expected |
| --- | --- | --- | --- |
| Arctic Cordillera | 242,769.43 | 2 | 29 |
| Atlantic Maritimes | 200,764.35 | 104 | 24 |
| Boreal Cordillera | 470,696.55 | 5 | 56 |
| Boreal Plains | 740,670.97 | 249 | 88 |
| Boreal Shield | 1,880,136.82 | 91 | 224 |
| Hudson Plains | 376,240.40 | 18 | 45 |
| Mixedwood Plains | 112,555.95 | 439 | 13 |
| Montane Cordillera | 489,781.38 | 87 | 58 |
| Northern Arctic | 1,516,869.25 | 14 | 181 |
| Pacific Maritimes | 209,358.98 | 73 | 25 |
| Prairies | 466,618.61 | 57 | 56 |
| Southern Arctic | 848,262.07 | 6 | 101 |
| Taiga Cordillera | 266,971.67 | 25 | 32 |
| Taiga Plains | 657,777.95 | 5 | 78 |
| Taiga Shield | 1,393,701.29 | 1 | 166 |

**S6: Current mean annual climate period chi-squared table**

**Table S6.** Sampled locations obtained from the literature for mean annual temperature and precipitation values (1991 – 2020). The ranges are divided into five equal intervals with increments of approximately 5.56 degrees and 757 mm for temperature (˚ C) and precipitation (mm), respectively. The expected number of sampled locations are based on the proportion of area represented by each temperature and precipitation category, assuming a search effort proportional to the area of each temperature and precipitation range.

| Variable | Range | Area | Observed | Expected |
| --- | --- | --- | --- | --- |
| Temperature (C) | < -11.23 | 1,594,280.70 | 14 | 190 |
|  | -11.23 to -5.67 | 1,795,791.66 | 29 | 214 |
|  | -5.67 to -0.11 | 3,177,075.53 | 63 | 379 |
|  | -0.11 to 5.45 | 2,911,703.17 | 438 | 347 |
|  | > 5.45 | 378,489.85 | 632 | 45 |
| Precipitation (mm) |  |  |  |  |
|  | Range | Area | Observed | Expected |
|  | < 646.47 | 6,647,258.12 | 452 | 806 |
|  | 646.47 to 1187.21 | 2,594,970.79 | 606 | 315 |
|  | 1187.21 to 1727.94 | 298,629.85 | 94 | 36 |
|  | 1727.94 to 2268.67 | 56,059.62 | 11 | 7 |
|  | 2268.67 to 2809.40 | 43,229.30 | 4 | 5 |
|  | 2809.40 to 3350.14 | 35,165.62 | 2 | 4 |
|  | 3350.14 to 3890.87 | 21,630.82 | 7 | 3 |

**S7: Projected mean annual climate chi-squared table**

**Table S7.** Sampled locations obtained from the literature for future annual temperature and precipitation values (2025 to 2100) for the Moderate emissions scenario (RCP 4.5). The ranges are divided into five equal intervals with increments of approximately 0.24 degrees and 5.79 mm for temperature (˚ C) and precipitation (mm), respectively. With respect to precipitation, negative numbers represent projected declines while positive numbers represent projected increases in precipitation, in reference to 1986 to 2005 period. The expected number of sampled locations are based on the proportion of area represented by each temperature and precipitation category.

| Variable | Category | Area (km2) | No. of observed sampled locations | No. of expected sampled locations |
| --- | --- | --- | --- | --- |
| Temperature (C) | < 0.86 | 903,697.97 | 40 | 109 |
|  | 0.86 to 1.11 | 1,362,536.45 | 197 | 164 |
|  | 1.11 to 1.35 | 2,572,022.02 | 279 | 309 |
|  | 1.35 to 1.59 | 3,813,235.20 | 529 | 458 |
|  | > 1.59 | 1,139,972.07 | 131 | 137 |
| Precipitation (mm) | < -6.82 | 34,509.93 | 11 | 4 |
|  | -6.82 to -2.06 | 821,484.45 | 110 | 99 |
|  | -2.06 to + 2.69 | 3,320,178.48 | 325 | 400 |
|  | 2.69 to 7.44 | 4,167,018.09 | 491 | 502 |
|  | > 7.44 | 1,424,219.02 | 239 | 171 |

**S8: Sampled locations in relation to the soil types in Canada**

Figure S8: The eleven different orders of soil in Canada (A). The semi-transparent black circles are the sampled locations. These circles appear darker when overlapping with multiple sampled locations. The inset maps (B) and (C) show areas with the largest number of sampled locations. The soil orders base layer is from Agriculture and Agri-Food Canada (2013). The coordinate system is WGS 84 (World Geodetic System 1984) and the coordinates are in decimal degrees. The map was created in R Core Team (2025) and Posit Software, PBC (2025) (version 2026.01.1) by Maheshi Dharmasiri.

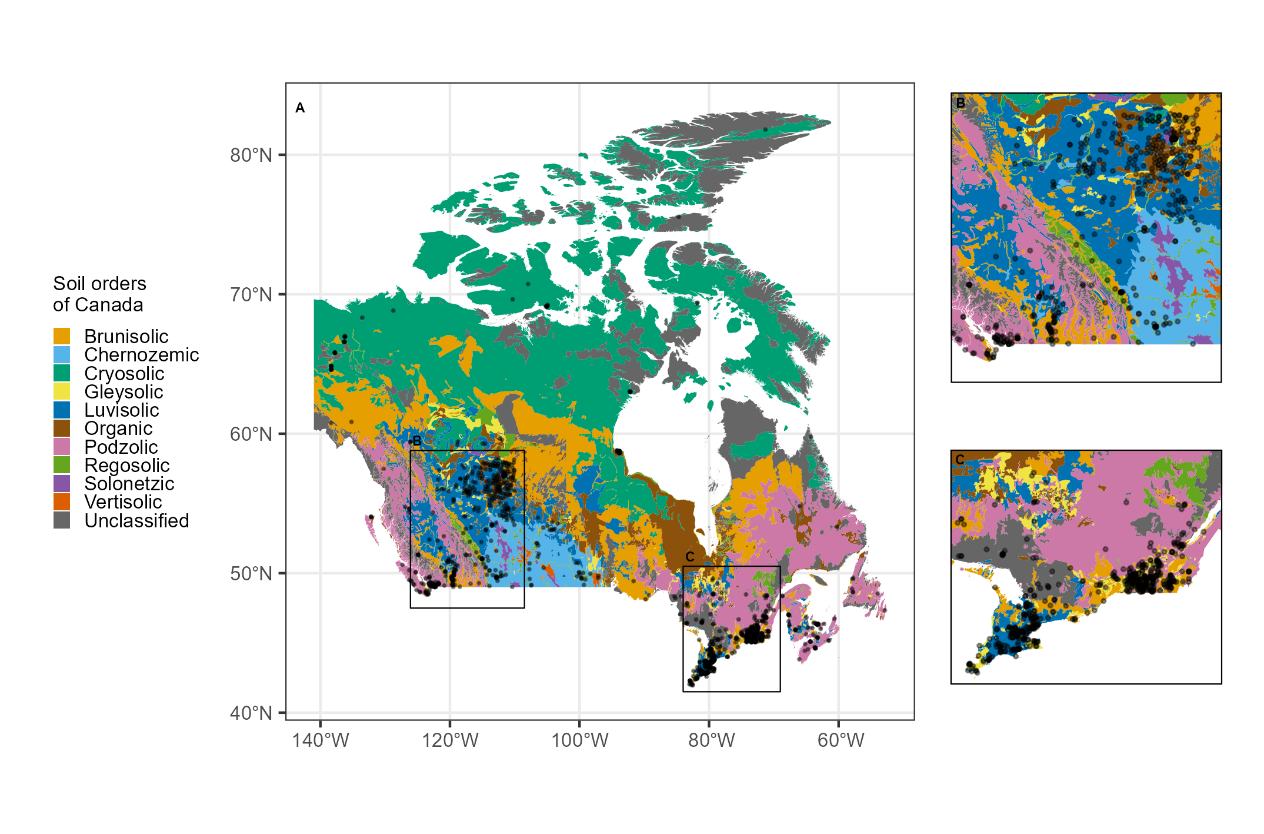

**S9: References for articles included in the study**

1. Abail, Z., and Whalen, J.K. 2018a. Selective ingestion contributes to the stoichiometric homeostasis in tissues of the endogeic earthworm Aporrectodea turgida. Soil Biology and Biochemistry 119: 121–127. doi:10.1016/j.soilbio.2018.01.014.
2. Abail, Z., and Whalen, J.K. 2018b. Corn residue inputs influence earthworm population dynamics in a no-till corn-soybean rotation. Applied Soil Ecology 127: 120–128. doi:10.1016/j.apsoil.2018.03.013.
3. Abail, Z., and Whalen, J.K. 2019. Nitrous oxide in vivo emission may regulate nitrogen stoichiometry in earthworm body tissues. European Journal of Soil Biology 91: 25–31. doi:10.1016/j.ejsobi.2019.01.002.
4. Abail, Z., and Whalen, J.K. 2021. Earthworm contributions to soil nitrogen supply in corn-soybean agroecosystems in Quebec, Canada. Pedosphere 31: 405–412. doi:10.1016/S1002-0160(20)60086-8.
5. Abbasi, P.A., Riga, E., Conn, K.L., and Lazarovits, G. 2005. Effect of neem cake soil amendment on reduction of damping-off severity and population densities of plant-parasitic nematodes and soilborne plant pathogens. Canadian Journal of Plant Pathology 27: 38–45. Taylor & Francis. doi:10.1080/07060660509507191.
6. Addison, J.A., and Holmes, S.B. 1995a. Comparison of forest soil microcosm and acute toxicity studies for determining effects of fenitrothion on earthworms. Ecotoxicology and Environmental Safety 30: 127–133. doi:10.1006/eesa.1995.1016.
7. Addison, J.A., and Holmes, S.B. 1995b. Effect of two commercial formulations of Bacillus thuringiensis subsp. kurstaki (Dipel 8L and Dipel 8AF) on the collembolan species folsomia candida in a soil microcosm study. Bull. Environ. Contam. Toxicol. 55. doi:10.1007/BF00203766.
8. Addison, J.A., Marshall, V.G., and Trofymow, J.A. 1998. Soil microarthropod abundance and species richness in successional Douglas-fir forests. Northwest Science 72: 96–97. Washington State University Press.
9. Addison, J.A., Otvos, I.S., Battigelli, J.P., and Conder, N. 2006. Does aerial spraying of Bacillus thuringiensis subsp. kurstaki (Btk) pose a risk to nontarget soil microarthropods? Can. J. For. Res. 36: 1610–1620. doi:10.1139/x06-048.
10. Addison, J.A., and Parkinson, D. 1978. Influence of Collembolan feeding activities on soil metabolism at a high Arctic site. Oikos 30: 529–538. Nordic Society Oikos, Wiley. doi:10.2307/3543348.
11. Addison, J.A., Trofymow, J.A., and Marshall, V.G. 2003a. Functional role of Collembola in successional coastal temperate forests on Vancouver Island, Canada. Applied Soil Ecology 24: 247–261. doi:10.1016/S0929-1393(03)00089-1.
12. Addison, J.A., Trofymow, J.A., and Marshall, V.G. 2003b. Abundance, species diversity, and community structure of Collembola in successional coastal temperate forests on Vancouver Island, Canada. Applied Soil Ecology 24: 233–246. doi:10.1016/S0929-1393(03)00090-8.
13. Adl, S.M., Ruiz‐Trillo, I., and Wilson, M.L. 2006. Observations on a Geocentrophora sp. (Lecithoepitheliata) flatworm from forest soils in Nova Scotia. Journal of Natural History 40: 1381–1387. Taylor & Francis. doi:10.1080/00222930600929285.
14. Adler, N., Bachmann, J., Blanckenhorn, W.U., Floate, K.D., Jensen, J., and Römbke, J. 2016. Effects of ivermectin application on the diversity and function of dung and soil fauna: Regulatory and scientific background information. Environ Toxicol Chem 35: 1914–1923. doi:10.1002/etc.3308.
15. Amankwa, G.A., White, A.D., McDowell, T.W., and Van Hooren, D.L. 2006. Pearl millet as a rotation crop with flue-cured tobacco for control of root-lesion nematodes in Ontario. Can. J. Plant Sci. 86: 1265–1271. NRC Research Press. doi:10.4141/P05-163.
16. Anas, O., and Reeleder, R.D. 1987. Recovery of fungi and arthropods from sclerotia of clerotinia clerotiorum in Quebec muck soils. Phytopathology 77: 327–331.
17. Angelone, S., and Bidochka, M.J. 2018. Diversity and abundance of entomopathogenic fungi at ant colonies. Journal of Invertebrate Pathology 156: 73–76. doi:10.1016/j.jip.2018.07.009.
18. Atul-Nayyar, Hamel, C., Forge, T., Selles, F., Jefferson, P.G., Hanson, K., and Germida, J. 2008. Arbuscular mycorrhizal fungi and nematodes are involved in negative feedback on a dual culture of alfalfa and Russian wildrye. Applied Soil Ecology 40: 30–36. doi:10.1016/j.apsoil.2008.03.004.
19. Ball-Coelho, B., Bruin, A.J., Roy, R.C., and Riga, E. 2003. Forage pearl millet and marigold as rotation crops for biological control of root-lesion nematodes in potato. Agronomy Journal 95: 282–292. doi:10.2134/agronj2003.2820.
20. Barreto, C., Branfireun, B.A., McLaughlin, J.W., and Lindo, Z. 2021. Responses of oribatid mites to warming in boreal peatlands depend on fen type. Pedobiologia 89: 150772. doi:10.1016/j.pedobi.2021.150772.
21. Barreto, C., and Lindo, Z. 2018. Drivers of decomposition and the detrital invertebrate community differ across a hummock-hollow microtopology in Boreal peatlands. Écoscience 25: 39–48. Taylor & Francis. doi:10.1080/11956860.2017.1412282.
22. Barreto, C., and Lindo, Z. 2021. Checklist of oribatid mites (Acari: Oribatida) from two contrasting boreal fens: An update on oribatid mites of Canadian peatlands. Systematic and Applied Acarology: 866–884. doi:10.11158/saa.26.5.4.
23. Battigelli, J.P., Berch, S.M., and Marshall, V.G. 1994. Soil fauna communities in two distinct but adjacent forest types on northern Vancouver Island, British Columbia. Can. J. For. Res. 24: 1557–1566. NRC Research Press. doi:10.1139/x94-203.
24. Battigelli, J.P., McIntyre, G.S., Broersma, K., and Krzic, M. 2003. Impact of cattle grazing on prostigmatid mite densities in grassland soils of southern interior British Columbia. Can. J. Soil. Sci. 83: 533–535. NRC Research Press. doi:10.4141/S02-064.
25. Battigelli, J.P., Spence, J.R., Langor, D.W., and Berch, S.M. 2004. Short-term impact of forest soil compaction and organic matter removal on soil mesofauna density and oribatid mite diversity. Can. J. For. Res. 34: 1136–1149. NRC Research Press. doi:10.1139/x03-267.
26. Bélair, G. 1991. Effects of preplant soil fumigation on nematode population densities, and on growth and yield of raspberry. phyto 72: 21–25. Société de protection des plantes du Québec (SPPQ)l. doi:10.7202/705999ar.
27. Bélair, G. 1998. Seasonal and vertical distribution of Meloidogyne hapla in organic soil. phyto 79: 1–8. Société de protection des plantes du Québec (SPPQ)l. doi:10.7202/706130ar.
28. Bélair, G., and Benoit, D.L. 1996a. Host suitability of 32 common weeds to Meloidogyne hapla in organic soils of southwestern Quebec. J Nematol 28: 643–647.
29. Bélair, G., and Benoit, D.L. 1996b. Host suitability of 32 common weeds to Meloidogyne hapla in organic soils of southwestern Quebec. J Nematol 28: 643–647. [Online] Available: https://pmc.ncbi.nlm.nih.gov/articles/PMC2619742/ [2026 May 1].
30. Bélair, G., Dauphinais, N., and Fournier, Y. 2019. Pathogenicity of Pratylenchus penetrans to dwarfing apple rootstocks. phyto 99: 12–14. Société de protection des plantes du Québec (SPPQ). doi:10.7202/1059304ar.
31. Bélair, G., Dauphinais, N., Fournier, Y., and Mauléon, H. 2001. Survey of plant-parasitic and entomopathogenic nematodes in vineyards of Quebec. phyto 82: 49–55. Société de protection des plantes du Québec (SPPQ)l. doi:10.7202/706215ar.
32. Bélair, G., Dauphinais, N., and Mimee, B. 2016. Evaluation of cultural methods for the management of the golden nematode (Globodera rostochiensis) in Quebec, Canada. Canadian Journal of Plant Pathology 38: 209–217. Taylor & Francis. doi:10.1080/07060661.2016.1189969.
33. Bélair, G., and Fournier, Y. 1997. Plant bed treatment with 1,3-dichloropropene for Meloidogyne hapla control in carrots grown in organic soil. phyto 78: 35–39. Société de protection des plantes du Québec (SPPQ)l. doi:10.7202/706117ar.
34. Belair, G., and Khanizadeh, S. 1994. Distribution of plant-parasitic nematodes in strawberry and raspberry fields in Quebec. Phytoprotection 75: 101–107. doi:10.7202/706056ar.
35. Bélair, G., and Parent, L.E. 1996. Using crop rotation to control Meloidogyne hapla Chitwood and improve marketable carrot yield. HortSci 31: 106–108. doi:10.21273/HORTSCI.31.1.106.
36. Bélair, G., and Tremblay, N. 1995. The influence of chitin-urea amendments applied to an organic soil on a Meloidogyne hapla population and on the growth of greenhouse tomato. phyto 76: 75–80. Société de protection des plantes du Québec (SPPQ)l. doi:10.7202/706087ar.
37. Belanger, G., Winch, J.E., and Townshed, J.L. 1985. Carbofuran effects on establishment of legumes in relation to plant growth, nitrogen fixation, and soil nematodes. Can. J. Plant Sci. 65: 423–433. NRC Research Press. doi:10.4141/cjps85-057.
38. Bell, A.J., Calladine, K.S., Wardle, D.A., and Phillips, I.D. 2022. Rapid colonization of the post-burn environment improves egg survival in pyrophilic ground beetles. Ecosphere 13: e4213. doi:10.1002/ecs2.4213.
39. Bendell-Young, L., Chouinard, J., and Pick, F.R. 1994. Metal concentrations in chrionomids in relation to peatland geochemistry. Arch. Environ. Contam. Toxicol. 27: 186–194. doi:10.1007/BF00214262.
40. Berch, S.M., Battigelli, J.P., and Hope, G.D. 2007. Responses of soil mesofauna communities and oribatid mite species to site preparation treatments in high-elevation cutblocks in southern British Columbia. Pedobiologia 51: 23–32. doi:10.1016/j.pedobi.2006.12.001.
41. Berch, S.M., Brockley, R.P., Battigelli, J., and Hagerman, S. 2009. Impacts of repeated fertilization on fine roots, mycorrhizas, mesofauna, and soil chemistry under young interior spruce in central British Columbia. Can. J. For. Res. 39: 889–896. NRC Research Press. doi:10.1139/X08-204.
42. Bird, G.A., and Chatarpaul, L. 1986. Effect of whole-tree and conventional forest harvest on soil microarthropods. Can. J. Zool. 64: 1986–1993. NRC Research Press. doi:10.1139/z86-299.
43. Bird, G.A., and Chatarpaul, L. 1988. Effect of forest harvest on decomposition and colonization of maple leaf litter by soil microarthropods. Can. J. Soil. Sci. 68: 29–40. NRC Research Press. doi:10.4141/cjss88-003.
44. Blauel, T., Celetti, M.J., Jordan, K.S., and Mcdonald, M.R. 2021. Optimizing methods to sample and quantify stem and bulb nematode, Ditylenchus dipsaci, in garlic, Allium sativum, field soil. Canadian Journal of Plant Pathology 43: 820–826. Taylor & Francis. doi:10.1080/07060661.2021.1928756.
45. Bodnaryk, R.P., Luo, M., and Kudryk, L. 1997. Effects of modifying the phytosterol profile of canola, Brassica napus L., on growth, development, and survival of the bertha armyworm, Mamestra configurata Walker (Lepidoptera: Noctuidae), the flea beetle, Phyllotreta cruciferae (Goeze) (Coleoptera: Chrysomelidae) and the aphids, Lipaphis erysimi (Kaltenbach) and Myzus persicae (Sulzer) (Homoptera: Aphididae). Can. J. Plant Sci. 77: 677–683. NRC Research Press. doi:10.4141/P97-011.
46. Boiteau, G., Goyer, C., Rees, H.W., and Zebarth, B.J. 2014. Differentiation of potato ecosystems on the basis of relationships among physical, chemical and biological soil parameters. Can. J. Soil. Sci. 94: 463–476. NRC Research Press. doi:10.4141/cjss2013-095.
47. Bradley, R.L., Whalen, J., Chagnon, P.-L., Lanoix, M., and Alves, M.C. 2011. Nitrous oxide production and potential denitrification in soils from riparian buffer strips: Influence of earthworms and plant litter. Applied Soil Ecology 47: 6–13. doi:10.1016/j.apsoil.2010.11.007.
48. Brandt, J.P., Smith, S.M., and Hubbes, M. 1996. Distribution and sampling of rood weevil larvae in young ornamental conifer plantations. The Canadian Entomologist 128: 1125–1133. doi:10.4039/Ent1281125-6.
49. Briar, S.S., Barker, C., Tenuta, M., and Entz, M.H. 2012. Soil nematode responses to crop management and conversion to native grasses. J Nematol 44: 245–254. [Online] Available: https://pmc.ncbi.nlm.nih.gov/articles/PMC3547337/ [2026 May 4].
50. Broadbent, A.B., and Tomlin, A.D. 1979. Species list of acari recovered from soil of a Guelph cornfield and a London pasture. Proceedings of the Entomological Society of Ontario 110: 101–103.
51. Broadbent, A.B., and Tomlin, A.D. 1982. Comparison of two methods for assessing the effects of Carbofuran on soil animal decomposers in cornfields. Environ Entomol 11: 1036–1042. doi:10.1093/ee/11.5.1036.
52. Broatch, J.S., Dosdall, L.M., Yang, R.-C., Harker, K.N., and Clayton, G.W. 2008. Emergence and seasonal activity of the entomophagous rove beetle Aleochara bilineata (Coleoptera: Staphylinidae) in canola in western Canada. Environ Entomol 37: 1451–1460. doi:10.1603/0046-225X-37.6.1451.
53. Button, M., Koch, I., and Reimer, K.J. 2012. Arsenic resistance and cycling in earthworms residing at a former gold mine in Canada. Environmental Pollution 169: 74–80. doi:10.1016/j.envpol.2012.04.031.
54. Button, M., Moriarty, M.M., Watts, M.J., Zhang, J., Koch, I., and Reimer, K.J. 2011. Arsenic speciation in field-collected and laboratory-exposed earthworms Lumbricus terrestris. Chemosphere 85: 1277–1283. doi:10.1016/j.chemosphere.2011.07.026.
55. Cameron, A., Boilard, G., Dubois, R., Bradley, R., Benetková, P., Józefowska, A., Thevathasan, N., Whalen, J., and Šimek, M. 2021. Distribution of earthworm communities in agroecosystems with forested riparian buffer strips: A multiscale study. Applied Soil Ecology 167: 104035. doi:10.1016/j.apsoil.2021.104035.
56. Cameron, E.K., and Bayne, E.M. 2011. An experimental test of facilitation between non-native earthworms. Can. J. Zool. 89: 1223–1230. NRC Research Press. doi:10.1139/z11-102.
57. Cameron, E.K., and Bayne, E.M. 2015. Spatial patterns and spread of exotic earthworms at local scales. Can. J. Zool. 93: 721–726. NRC Research Press. doi:10.1139/cjz-2014-0197.
58. Cameron, E.K., Knysh, K.M., Proctor, H.C., and Bayne, E.M. 2013a. Influence of two exotic earthworm species with different foraging strategies on abundance and composition of boreal microarthropods. Soil Biology and Biochemistry 57: 334–340. doi:10.1016/j.soilbio.2012.07.010.
59. Cameron, E.K., Proctor, H.C., and Bayne, E.M. 2013b. Effects of an ecosystem engineer on belowground movement of microarthropods. PLOS ONE 8: e62796. Public Library of Science. doi:10.1371/journal.pone.0062796.
60. Cameron, E.K., Zabrodski, M.W., Karst, J., and Bayne, E.M. 2012. Non-native earthworm influences on ectomycorrhizal colonization and growth of white spruce. Écoscience 19: 29–37. Taylor & Francis. doi:10.2980/19-1-3478.
61. Cárcamo, H.A., and Parkinson, D. 2001. Localized acidification near sour gas processing plants: are forest floor macro-invertebrates affected? Applied Soil Ecology 17: 199–213. doi:10.1016/S0929-1393(01)00140-8.
62. Cárcamo, H.A., Parkinson, D., and Bargshoon, D. 1998a. Distribution of earthworms along a sharp acidification gradient. Pedobiologia 42: 88–95. doi:10.1016/S0031-4056(24)00396-2.
63. Cárcamo, H.A., Parkinson, D., and Volney, J.W.A. 1998b. Effects of sulphur contamination on macroinvertebrates in Canadian pine forests. Applied Soil Ecology 9: 459–464. doi:10.1016/S0929-1393(98)00105-X.
64. Caron, A.-S., Koudji, E.G., Handa, I.T., Montoro Girona, M., and Despland, E. 2023. Forest tent caterpillar outbreaks drive change in ant communities in boreal forests. Forests 14: 1147. Multidisciplinary Digital Publishing Institute. doi:10.3390/f14061147.
65. Carter, A. 1983. Cadmium, copper, and zinc in soil animals and their food in a red clover system. Can. J. Zool. 61: 2751–2757. NRC Research Press. doi:10.1139/z83-361.
66. Carter, A., and Cragg, J.B. 1976. Concentrations and standing crops of calcium, magnesium, potassium and sodium in soil and litter arthropods and their food in an aspen woodland ecosystem in the Rocky Mountains (Canada). Pedobiologia 16: 379–388. doi:10.1016/S0031-4056(23)02192-3.
67. Carter, M.R., Kunelius, H.T., Sanderson, J.B., Kimpinski, J., Platt, H.W., and Bolinder, M.A. 2003. Productivity parameters and soil health dynamics under long-term 2-year potato rotations in Atlantic Canada. Soil and Tillage Research 72: 153–168. doi:10.1016/S0167-1987(03)00085-0.
68. Carter, M.R., and Noronha, C. 2007. Soil micro-arthropod communities and microbial parameters in the potato ridge under two field management systems on sandy loams in Atlantic Canada. Can. J. Soil. Sci. 87: 399–404. NRC Research Press. doi:10.4141/CJSS06013.
69. Carter, M.R., Noronha, C., Peters, R.D., and Kimpinski, J. 2009a. Influence of conservation tillage and crop rotation on the resilience of an intensive long-term potato cropping system: Restoration of soil biological properties after the potato phase. Agriculture, Ecosystems & Environment 133: 32–39. doi:10.1016/j.agee.2009.04.017.
70. Carter, M.R., Peters, R.D., Noronha, C., and Kimpinski, J. 2009b. Influence of 10 years of conservation tillage on some biological properties of a fine sandy loam in the potato phase of two crop rotations in Atlantic Canada. Can. J. Soil. Sci. 89: 391–402. NRC Research Press. doi:10.4141/cjss08054.
71. Carter, M.R., Sanderson, J.B., Ivany, J.A., and White, R.P. 2002. Influence of rotation and tillage on forage maize productivity, weed species, and soil quality of a fine sandy loam in the cool–humid climate of Atlantic Canada. Soil and Tillage Research 67: 85–98. doi:10.1016/S0167-1987(02)00043-0.
72. Carter, N.E., and Brown, N.R. 1973. Seasonal abundance of certain soil arthropods in a fenitrothion-treated red spruce stand. The Canadian Entomologist 105: 1065–1073. doi:10.4039/Ent1051065-8.
73. Chagnon, M., Hébert, C., and Paré, D. 2000a. Community structures of Collembola in sugar maple forests: Relations to humus type and seasonal trends. Pedobiologia 44: 148–174. doi:10.1078/S0031-4056(04)70035-9.
74. Chagnon, M., Paré, D., and Hébert, C. 2000b. Relationships between soil chemistry, microbial biomass and the collembolan fauna of southern Québec sugar maple stands. Écoscience 7: 307–316. Taylor & Francis. doi:10.1080/11956860.2000.11682600.
75. Chagnon, M., Paré, D., Hébert, C., and Camiré, C. 2001. Effects of experimental liming on collembolan communities and soil microbial biomass in a southern Quebec sugar maple (Acer saccharum Marsh.) stand. Applied Soil Ecology 17: 81–90. doi:10.1016/S0929-1393(00)00134-7.
76. Chang, X.C., and Juma, N.G. 1996. Impact of crop rotations on microbial biomass, faunal populations, and plant C and N in a Gray Luvisol (Typic Cryoboralf). Biol Fert Soils 22: 31–39. doi:10.1007/BF00384429.
77. Chen, C., Ejack, L., Chénier, M.R., and Whalen, J.K. 2021. Earthworm functional groups are related to denitrifier activity in riparian soils. Pedosphere 31: 413–422. doi:10.1016/S1002-0160(20)60084-4.
78. Chen, C., Whalen, J.K., and Guo, X. 2014a. Earthworms reduce soil nitrous oxide emissions during drying and rewetting cycles. Soil Biology and Biochemistry 68: 117–124. doi:10.1016/j.soilbio.2013.09.020.
79. Chen, D., Zebarth, B.J., Goyer, C., Comeau, L.-P., Nahar, K., and Dixon, T. 2022. Effect of biofumigation on population densities of Pratylenchus spp. and Verticillium spp. and potato yield in eastern Canada. Am. J. Potato Res. 99: 229–242. doi:10.1007/s12230-022-09875-2.
80. Chen, Y., Amundrud, S.L., and Srivastava, D.S. 2014b. Spatial variance in soil microarthropod communities: Niche, neutrality, or stochasticity? Écoscience 21: 405–418. Taylor & Francis. doi:10.2980/21-(3-4)-3720.
81. Chevalier, R., and Webster, J.M. 2006. Temporary changes in populations of soil organisms after field application of entomopathogenic nematodes. J Nematol 38: 466–473. [Online] Available: https://pmc.ncbi.nlm.nih.gov/articles/PMC2586464/ [2026 May 4].
82. Choi, A., Sackett, T.E., Smith, S.M., and Bellocq, M.I. 2017. Exotic earthworm (Oligochaeta: Lumbricidae) assemblages on a landscape scale in central Canadian woodlands: Importance of region and vegetation type. Can. J. For. Res. 47: 935–945. NRC Research Press. doi:10.1139/cjfr-2016-0337.
83. Chomel, M., Guittonny-Larchevêque, M., DesRochers, A., and Baldy, V. 2016. Effect of mixing herbaceous litter with tree litters on decomposition and N release in boreal plantations. Plant Soil 398: 229–241. doi:10.1007/s11104-015-2648-5.
84. Clapperton, M.J. 1999. Tillage practices, and temperature and moisture interactions affect earthworm populations and species composition. Pedobiologia 43: 658–665. doi:10.1016/S0031-4056(24)00595-X.
85. Clapperton, M.J., Kanashiro, D.A., and Behan-Pelletier, V.M. 2002. Changes in abundance and diversity of microarthropods associated with Fescue Prairie grazing regimes. Pedobiologia 46: 496–511. doi:10.1078/0031-4056-00155.
86. Clapperton, M.J., Lee, N.O., Binet, F., and Conner, R.L. 2001. Earthworms indirectly reduce the effects of take-all (Gaeumannomyces graminis var. tritici) on soft white spring wheat (Triticum aestivum cv. Fielder). Soil Biology and Biochemistry 33: 1531–1538. doi:10.1016/S0038-0717(01)00071-2.
87. Coffey, V., and Otfinowski, R. 2019. Legacies of afforestation on soil nematode community composition, structure, and diversity in a northern Canadian prairie. Plant Soil 435: 437–447. doi:10.1007/s11104-018-03906-w.
88. Conn, K.L., and Lazarovits, G. 1999. Impact of animal manures on verticillium wilt, potato scab, and soil microbial populations. Canadian Journal of Plant Pathology 21: 81–92. Taylor & Francis. doi:10.1080/07060661.1999.10600089.
89. Coors, A., Edwards, M., Lorenz, P., Römbke, J., Schmelz, R.M., Topp, E., Waszak, K., Wilkes, G., and Lapen, D.R. 2016. Biosolids applied to agricultural land: Influence on structural and functional endpoints of soil fauna on a short- and long-term scale. Science of The Total Environment 562: 312–326. doi:10.1016/j.scitotenv.2016.03.226.
90. Corkery, C.A., Nol, E., and Mckinnon, L. 2019. No effects of asynchrony between hatching and peak food availability on chick growth in semipalmated plovers (Charadrius semipalmatus) near Churchill, Manitoba. Polar Biol 42: 593–601. doi:10.1007/s00300-019-02456-w.
91. Cormier, D., and Martel, P. 1997. Effects of soil insecticide treatments on northern corn rootworm, Diabrotica barberi [Coleoptera : Chrysomelidae] populations and on corn yield. phyto 78: 67–73. Société de protection des plantes du Québec (SPPQ)l. doi:10.7202/706121ar.
92. Coupe, M.D., Stacey, J.N., and Cahill, J.F.Jr. 2009. Limited effects of above- and belowground insects on community structure and function in a species-rich grassland. Journal of Vegetation Science 20: 121–129. Wiley. [Online] Available: https://www.jstor.org/stable/40295847 [2026 May 1].
93. Cray, H., Gaudon, J., and Murphy, S. 2021. Introduced earthworms (Lumbricidae) in restored and remnant tallgrass prairies of southern Ontario. The Canadian Field-Naturalist 135: 234–244. doi:10.22621/cfn.v135i3.2721.
94. Crotty, F.V., and Adl, S.M. 2019. Competition and predation in soil fungivorous microarthropods using stable isotope ratio mass spectrometry. Front. Microbiol. 10. Frontiers. doi:10.3389/fmicb.2019.01274.
95. Currie, C.R., Spence, J.R., and Volney, W.J.A. 1996. Biology and life histroy of Epuraea obliquus hatch (Coleoptera: Nitidulidae) on western gall rust. The Canadian Entomologist 128: 177–186. doi:10.4039/Ent128177-2.
96. Currier, H.A., Fremlin, K.M., Elliott, J.E., Drouillard, K.G., and Williams, T.D. 2020. Bioaccumulation and biomagnification of PBDEs in a terrestrial food chain at an urban landfill. Chemosphere 238: 124577. doi:10.1016/j.chemosphere.2019.124577.
97. Dash, M.C., and Cragg, J.B. 1972. Ecology of Enchytraeidae (Oligochaeta) in Canadian Rocky Mountain Soils. Pedobiologia 12: 323–335. doi:10.1016/S0031-4056(23)02055-3.
98. Dauphinais, N., Vandal, M., Gagnon, A.-È., Bélair, G., Véronneau, P.-Y., and Mimee, B. 2018. Development and application of a multiplex qPCR method for the simultaneous detection and quantification of Pratylenchus alleni and P. penetrans in Quebec, Canada. Plant Disease 102: 970–976. Scientific Societies. doi:10.1094/PDIS-08-17-1222-RE.
99. Déchêne, A., and Buddle, C. 2009a. Decomposing logs increase oribatid mite assemblage diversity in mixedwood boreal forest. Biodiversity and Conservation 19: 237–256. doi:10.1007/s10531-009-9719-y.
100. Déchêne, A.D., and Buddle, C.M. 2009b. Effects of experimental forest harvesting on oribatid mite biodiversity. Forest Ecology and Management 258: 1331–1341. doi:10.1016/j.foreco.2009.06.033.
101. Díaz-Aguilar, I., and Quideau, S.A. 2013. Trophic ecology of mesostigmatan and oribatid mites in harvested and control coniferous and deciduous stands of the boreal mixedwood forest determined using 15N stable isotopes. Soil Biology and Biochemistry 67: 147–154. doi:10.1016/j.soilbio.2013.08.019.
102. Díaz-Aguilar, I., Quideau, S.A., Proctor, H.C., Kishchuk, B.E., and Spence, J.R. 2013. Influence of stand composition on predatory mite (Mesostigmata) assemblages from the forest floor in western Canadian boreal mixedwood forests. Forest Ecology and Management 309: 105–114. doi:10.1016/j.foreco.2013.03.029.
103. Diogène, J., Dufour, M., Poirier, G.G., and Nadeau, D. 1997. Extrusion of earthworm coelomocytes: comparison of the cell populations recovered from the species Lumbricus terrestris, Eisenia fetida and Octolasion tyrtaeum. Lab Anim 31: 326–336. doi:10.1258/002367797780596068.
104. Diop, H., Ejack, L., Elmi, A.A., and Whalen, J.K. 2023. Earthworm distribution around tile drainage in agricultural fields of southwest Quebec, Canada. Can. J. Soil. Sci. 103: 377–380. NRC Research Press. doi:10.1139/cjss-2022-0059.
105. Doane, J.F. 1977. Spatial pattern and density oof Ctenicera destructor and Hypolithus bicolor (Coleoptera: Elateridae) in soil in spring wheat. The Canadian Entomologist 109: 807–822. doi:10.4039/Ent109807-6.
106. Doane, J.F., Mukerji, M.K., and Olfert, O. 2000. Sampling distribution and sequential sampling for subterranean stages of orange wheat blossom midge, Sitodiplosis mosellana (Géhin) (Diptera: Cecidomyiidae) in spring wheat. Crop Protection 19: 427–434. doi:10.1016/S0261-2194(00)00035-1.
107. Doane, J.F., and Olfert, O. 2008. Seasonal development of wheat midge, Sitodiplosis mosellana (Géhin) (Diptera: Cecidomyiidae), in Saskatchewan, Canada. Crop Protection 27: 951–958. doi:10.1016/j.cropro.2007.11.016.
108. Doane, J.F., Olfert, O.O., and Mukerji, M.K. 1987. Extraction precision of sieving and brine flotation for removal of wheat midge, Sitodiplosis mosellana (Diptera: Cecidomyiidae), cocoons and larvae from soil. J Econ Entomol 80: 268–271. doi:10.1093/jee/80.1.268.
109. Doblas-Miranda, E., Paquette, A., and Work, T.T. 2014. Intercropping trees’ effect on soil oribatid diversity in agro-ecosystems. Agroforest Syst 88: 671–678. doi:10.1007/s10457-014-9680-y.
110. Doblas-Miranda, E., and Work, T.T. 2015. Localized effects of coarse woody material on soil oribatid communities diminish over 700 years of stand development in black-spruce-feathermoss forests. Forests 6: 914–928. Multidisciplinary Digital Publishing Institute. doi:10.3390/f6040914.
111. Dominique, C.R., Yule, W.N., and Martel, P. 1983. Influence of soil type, soil moisture, and soil surface conditions on oviposition preference of the northern corn rootworm. The Canadian Entomologist 115: 1043–1046. doi:10.4039/Ent1151043-8.
112. Dosdall, L.M., Clayton, G.W., Neil Harker, K., O’Donovan, J.T., and Craig Stevenson, F. 2004. The effects of soil fertility and other agronomic factors on infestations of root maggots in canola. Agronomy Journal 96: 1306–1313. doi:10.2134/agronj2004.1306.
113. Drahun, I., Hunt, K.A., van Herk, W.G., and Cassone, B.J. 2022. Evaluation of standardized bait trapping approaches and climatic factors that influence wireworm catch in the Canadian Prairies. J Econ Entomol 115: 773–782. doi:10.1093/jee/toac037.
114. Drahun, I., Poole, E.A., Hunt, K.A., van Herk, W.G., LeMoine, C.M., and Cassone, B.J. 2023. Seasonal turnover and insights into the overwintering biology of wireworms (Coleoptera: Elateridae) in the Canadian Prairies. Pest Manag Sci 79: 526–536. doi:10.1002/ps.7222.
115. Drahun, I., Rutherford, P.L., van Herk, W.G., and Cassone, B.J. 2024. Wireworm feeding behaviours: The capacity of Hypnoidus bicolor to damage soybean under different environmental conditions. Arthropod-Plant Interactions 18: 129–139. doi:10.1007/s11829-023-10009-6.
116. Drijber, R.A., and McGill, W.B. 1994. Sulfonolipids as a biomarker to monitor the population dynamics of the genera Cytophaga and Flexibacter in soil worked by the earthworm Aporrectodea turgida. Soil Biology and Biochemistry 26: 1395–1403. doi:10.1016/0038-0717(94)90223-2.
117. Drouin, M., Bradley, R., and Lapointe, L. 2016. Linkage between exotic earthworms, understory vegetation and soil properties in sugar maple forests. Forest Ecology and Management 364: 113–121. doi:10.1016/j.foreco.2016.01.010.
118. Duceppe, M.-O., Lafond-Lapalme, J., Palomares-Rius, J.E., Sabeh, M., Blok, V., Moffett, P., and Mimee, B. 2017. Analysis of survival and hatching transcriptomes from potato cyst nematodes, Globodera rostochiensis and G. pallida. Sci Rep 7: 3882. Nature Publishing Group. doi:10.1038/s41598-017-03871-x.
119. Dwyer, E., Larson, D.J., and Thompson, I.D. 1997. Oribatida (Acari) in balsam fir [Abies balsamea (L.)] forests of western Newfoundland. The Canadian Entomologist 129: 151–170. doi:10.4039/Ent129151-1.
120. Dymond, P., Scheu, S., and Parkinson, D. 1997. Density and distribution of Dendrobaena octaedra (Lumbricidae) in aspen and pine forests in the Canadian Rocky Mountains (Alberta). Soil Biology and Biochemistry 29: 265–273. doi:10.1016/S0038-0717(96)00052-1.
121. Edwards, L.M., and Kimpinski, J. 1997. Relationships between soil penetration resistance and soif nematode burden in barley on Prince Edward Island. Biol Fert Soils 24: 13–17. doi:10.1007/BF01420214.
122. Eisenhauer, N., Partsch, S., Parkinson, D., and Scheu, S. 2007. Invasion of a deciduous forest by earthworms: Changes in soil chemistry, microflora, microarthropods and vegetation. Soil Biology and Biochemistry 39: 1099–1110. doi:10.1016/j.soilbio.2006.12.019.
123. Eisenhauer, N., Straube, D., Johnson, E.A., Parkinson, D., and Scheu, S. 2009. Exotic ecosystem engineers change the emergence of plants from the seed bank of a deciduous forest. Ecosystems 12: 1008–1016. doi:10.1007/s10021-009-9275-z.
124. Eisenhauer, N., Yee, K., Johnson, E.A., Maraun, M., Parkinson, D., Straube, D., and Scheu, S. 2011. Positive relationship between herbaceous layer diversity and the performance of soil biota in a temperate forest. Soil Biology and Biochemistry 43: 462–465. doi:10.1016/j.soilbio.2010.10.018.
125. Ejack, L., Kernecker, M.L., Prieto, R., Chen, C., Gul, S., Bradley, R.L., and Whalen, J.K. 2021. Earthworms did not increase long-term nitrous oxide fluxes in perennial forage and riparian buffer ecosystems. Pedobiologia 85–86: 150727. doi:10.1016/j.pedobi.2021.150727.
126. Eriksen-Hamel, N.S., Speratti, A.B., Whalen, J.K., Légère, A., and Madramootoo, C.A. 2009. Earthworm populations and growth rates related to long-term crop residue and tillage management. Soil and Tillage Research 104: 311–316. doi:10.1016/j.still.2009.04.006.
127. Eriksen-Hamel, N.S., and Whalen, J.K. 2006. Growth rates of Aporrectodea caliginosa (Oligochaetae: Lumbricidae) as influenced by soil temperature and moisture in disturbed and undisturbed soil columns. Pedobiologia 50: 207–215. doi:10.1016/j.pedobi.2005.10.008.
128. Eriksen-Hamel, N.S., and Whalen, J.K. 2007a. Impacts of earthworms on soil nutrients and plant growth in soybean and maize agroecosystems. Agriculture, Ecosystems & Environment 120: 442–448. doi:10.1016/j.agee.2006.11.004.
129. Eriksen-Hamel, N.S., and Whalen, J.K. 2007b. Competitive interactions affect the growth of Aporrectodea caliginosa and Lumbricus terrestris (Oligochaeta: Lumbricidae) in single- and mixed-species laboratory cultures. European Journal of Soil Biology 43: 142–150. doi:10.1016/j.ejsobi.2006.11.005.
130. Eriksen-Hamel, N.S., and Whalen, J.K. 2008. Earthworms, soil mineral nitrogen and forage production in grass-based hayfields. Soil Biology and Biochemistry 40: 1004–1010. doi:10.1016/j.soilbio.2007.11.013.
131. Eriksen-Hamel, N.S., and Whalen, J.K. 2009. The “deduction” approach: A non-invasive method for estimating secondary production of earthworm communities. Acta Oecologica 35: 477–484. doi:10.1016/j.actao.2008.11.001.
132. Estevez, B., N’Dayegamiye, A., and Coderre, D. 1996. The effect on earthworm abundance and selected soil properties after 14 years of solid cattle manure and NPKMg fertilizer application. Can. J. Soil. Sci. 76: 351–355. NRC Research Press. doi:10.4141/cjss96-042.
133. Fagan, L.L., Didham, R.K., Winchester, N.N., Behan-Pelletier, V., Clayton, M., Lindquist, E., and Ring, R.A. 2006. An experimental assessment of biodiversity and species turnover in terrestrial vs canopy leaf litter. Oecologia 147: 335–347. doi:10.1007/s00442-005-0262-6.
134. Fitzpatrick, C.R., Mikhailitchenko, A.V., Anstett, D.N., and Johnson, M.T.J. 2018a. The influence of range-wide plant genetic variation on soil invertebrate communities. Ecography 41: 1135–1146. doi:10.1111/ecog.03347.
135. Fitzpatrick, S.M., Wong, W.H.L., Matthews, K., Mathur, S., Elsby, M., Schurmann, K., and Craig, L.N. 2018b. A bucket-type emergence trap for detecting overwintered Dasineura oxycoccana (Diptera: Cecidomyiidae) and its parasitoids in cranberry. flen 101: 695–698. Florida Entomological Society. doi:10.1653/024.101.0410.
136. Fleri, J.R., and Arcese, P. 2021. Predictive mapping to identify refuges for plant communities threatened by earthworm invasion. Ecological Solutions and Evidence 2: e12064. doi:10.1002/2688-8319.12064.
137. Fons, J., and Klinka, K. 1998. Temporal variations of forest floor properties in the Coastal Western Hemlock zone of southern British Columbia. Can. J. For. Res. 28: 582–590. NRC Research Press. doi:10.1139/x98-027.
138. Fons, J., Klinka, K., and Kabzems, R.D. 1998. Humus forms of trembling aspen ecosystems in northeastern British Columbia. Forest Ecology and Management 105: 241–250. doi:10.1016/S0378-1127(97)00290-9.
139. Forge, T., Ehret, D., Messiga, A., and Dorais, M. 2020a. Influences of nitrogen inputs on nematode populations under highbush blueberry. J Nematol 52: 1–14. doi:10.21307/jofnem-2020-056.
140. Forge, T., Kenney, E., Hashimoto, N., Neilsen, D., and Zebarth, B. 2016. Compost and poultry manure as preplant soil amendments for red raspberry: Comparative effects on root lesion nematodes, soil quality and risk of nitrate leaching. Agriculture, Ecosystems & Environment 223: 48–58. doi:10.1016/j.agee.2016.02.024.
141. Forge, T., Munro, P., Wright, H., and Moreau, D. 2022. Plant-parasitic nematodes in Nova Scotia vineyards. phyto 102: 15–20. Société de protection des plantes du Québec (SPPQ). doi:10.7202/1088485ar.
142. Forge, T., and Simard, S. 2001a. Structure of nematode communities in forest soils of southern British Columbia: relationships to nitrogen mineralization and effects of clearcut harvesting and fertilization. Biol Fertil Soils 34: 170–178. doi:10.1007/s003740100390.
143. Forge, T., Smit, R., Neilsen, D., and Neilsen, G. 2020b. Potential impacts of the ring nematode, Mesocriconema xenoplax, on grapevines in British Columbia: A microplot study. Journal of Nematology 52: 1–7. doi:10.21307/jofnem-2020-086.
144. Forge, T.A., Bittman, S., and Kowalenko, C.G. 2005a. Impacts of sustained use of dairy manure slurry and fertilizers on populations of Pratylenchus penetrans under tall fescue. Journal of Nematology 37: 207–213.
145. Forge, T.A., Bittman, S., and Kowalenko, C.G. 2005b. Responses of grassland soil nematodes and protozoa to multi-year and single-year applications of dairy manure slurry and fertilizer. Soil Biology and Biochemistry 37: 1751–1762. doi:10.1016/j.soilbio.2004.11.013.
146. Forge, T.A., Hogue, E., Neilsen, G., and Neilsen, D. 2003. Effects of organic mulches on soil microfauna in the root zone of apple: Implications for nutrient fluxes and functional diversity of the soil food web. Applied Soil Ecology 22: 39–54. doi:10.1016/S0929-1393(02)00111-7.
147. Forge, T.A., Hogue, E.J., Neilsen, G., and Neilsen, D. 2008. Organic mulches alter nematode communities, root growth and fluxes of phosphorus in the root zone of apple. Applied Soil Ecology 39: 15–22. doi:10.1016/j.apsoil.2007.11.004.
148. Forge, T.A., and Kempler, C. 2009. Organic mulches influence population densities of root-lesion nematodes, soil health indicators, and root growth of red raspberry. Canadian Journal of Plant Pathology 31: 241–249. Taylor & Francis. doi:10.1080/07060660909507597.
149. Forge, T.A., Larney, F.J., Kawchuk, L.M., Pearson, D.C., Koch, C., and Blackshaw, R.E. 2015. Crop rotation effects on Pratylenchus neglectus populations in the root zone of irrigated potatoes in southern Alberta. Canadian Journal of Plant Pathology 37: 363–368. Taylor & Francis. doi:10.1080/07060661.2015.1066864.
150. Forge, T.A., and Simard, S.W. 2000. Trophic structure of nematode communities, microbial biomass, and nitrogen mineralization in soils of forests and clearcuts in the southern interior of British Columbia. Can. J. Soil. Sci. 80: 401–410. NRC Research Press. doi:10.4141/S99-112.
151. Forge, T.A., and Simard, S.W. 2001b. Short-term effects of nitrogen and phosphorus fertilizers on nitrogen mineralization and trophic structure of the soil ecosystem in forest clearcuts in the southern interior of British Columbia. Can. J. Soil. Sci. 81: 11–20. NRC Research Press. doi:10.4141/S00-018.
152. Fox, C.A., Fonseca, E.J.A., Miller, J.J., and Tomlin, A.D. 1999. The influence of row position and selected soil attributes on Acarina and Collembola in no-till and conventional continuous corn on a clay loam soil. Applied Soil Ecology 13: 1–8. doi:10.1016/S0929-1393(98)00167-X.
153. Fox, C.A., Miller, J.J., Joschko, M., Drury, C.F., and Reynolds, W.D. 2017. Earthworm population dynamics as a consequence of long-term and recently imposed tillage in a clay loam soil. Can. J. Soil. Sci. 97: 561–579. NRC Research Press. doi:10.1139/cjss-2016-0092.
154. Fugère, M., Bradley, R.L., and Vellend, M. 2016. Exotic earthworms dispersion through protected forest areas and their potential impacts on nitrous oxide production. Biol Invasions 19: 773–783. doi:10.1007/s10530-016-1331-y.
155. Gagnon Koudji, E., Despland, E., Caron, A.-S., and Handa, I.T. 2023. Soil springtail communities are resilient to forest tent caterpillar defoliation in Quebec mixed hardwood forests. Forests 14: 1302. Multidisciplinary Digital Publishing Institute. doi:10.3390/f14071302.
156. Gan, S.H., McTavish, M.J., Bourchier, R.S., and Smith, S.M. 2023. Soil grit influences the effects of earthworms (Lumbricus terrestris) on seed damage. Applied Soil Ecology 185: 104807. doi:10.1016/j.apsoil.2023.104807.
157. Gans, C., and Burr, A.H.J. 1994. Unique locomotory mechanism of Mermis nigrescens, a large nematode that crawls over soil and climbs through vegetation. J Morphol 222: 133–148. doi:10.1002/jmor.1052220203.
158. George, P.B.L., and Lindo, Z. 2015a. Congruence of community structure between taxonomic identification and T-RFLP analyses in free-living soil nematodes. Pedobiologia 58: 113–117. doi:10.1016/j.pedobi.2015.04.003.
159. George, P.B.L., and Lindo, Z. 2015b. Application of body size spectra to nematode trait-index analyses. Soil Biology and Biochemistry 84: 15–20. doi:10.1016/j.soilbio.2015.02.007.
160. Gewurtz, S.B., Tardif, G., Power, M., Backus, S.M., Dove, A., Dubé-Roberge, K., Garron, C., King, M., Lalonde, B., Letcher, R.J., Martin, P.A., McDaniel, T.V., McGoldrick, D.J., Pelletier, M., Small, J., Smyth, S.A., Teslic, S., and Tessier, J. 2021. Bisphenol A in the Canadian environment: A multimedia analysis. Science of The Total Environment 755: 142472. doi:10.1016/j.scitotenv.2020.142472.
161. Gong, X., Li, J., Chang, S.X., Wu, Q., An, Z., Huang, C., Sun, X., Li, S., and Wang, H. 2021. Cattle manure biochar and earthworm interactively affected CO2 and N2O emissions in agricultural and forest soils: Observation of a distinct difference. Front. Environ. Sci. Eng. 16: 39. doi:10.1007/s11783-021-1473-8.
162. Guerra, C.A., Berdugo, M., Eldridge, D.J., Eisenhauer, N., Singh, B.K., Cui, H., Abades, S., Alfaro, F.D., Bamigboye, A.R., Bastida, F., Blanco-Pastor, J.L., de los Ríos, A., Durán, J., Grebenc, T., Illán, J.G., Liu, Y.-R., Makhalanyane, T.P., Mamet, S., Molina-Montenegro, M.A., Moreno, J.L., Mukherjee, A., Nahberger, T.U., Peñaloza-Bojacá, G.F., Plaza, C., Picó, S., Verma, J.P., Rey, A., Rodríguez, A., Tedersoo, L., Teixido, A.L., Torres-Díaz, C., Trivedi, P., Wang, J., Wang, L., Wang, J., Zaady, E., Zhou, X., Zhou, X.-Q., and Delgado-Baquerizo, M. 2022. Global hotspots for soil nature conservation. Nature 610: 693–698. Nature Publishing Group. doi:10.1038/s41586-022-05292-x.
163. Guerrieri, A., Cantera, I., Marta, S., Bonin, A., Carteron, A., Ambrosini, R., Caccianiga, M., Anthelme, F., Azzoni, R.S., Almond, P., Alviz Gazitúa, P., Cauvy-Fraunié, S., Ceballos Lievano, J.L., Chand, P., Chand Sharma, M., Clague, J., Cochachín Rapre, J.A., Compostella, C., Cruz Encarnación, R., Dangles, O., Deline, P., Eger, A., Erokhin, S., Franzetti, A., Gielly, L., Gili, F., Gobbi, M., Hågvar, S., Khedim, N., Meneses, R.I., Peyre, G., Pittino, F., Proietto, A., Rabatel, A., Urseitova, N., Yang, Y., Zaginaev, V., Zerboni, A., Zimmer, A., Taberlet, P., Diolaiuti, G.A., Poulenard, J., Fontaneto, D., Thuiller, W., and Ficetola, G.F. 2024. Local climate modulates the development of soil nematode communities after glacier retreat. Global Change Biology 30: e17057. doi:10.1111/gcb.17057.
164. Guppy, J.C., and Harcourt, D.G. 1990. Seasonal history and behavior of the alfalfa snout beetle, Otiorhynchus ligustici (Coleoptera: Curculionidae), in eastern Ontario. Proceedings of the Entomological Society of Ontario 121: 61–69.
165. Gyug, L.W., Higgins, R.J., Todd, M.A., Meggs, J.M., and Lindgren, B.S. 2014. Dietary dependence of Williamson’s Sapsucker nestlings on ants associated with dead and decaying wood in British Columbia. Can. J. For. Res. 44: 628–637. NRC Research Press. doi:10.1139/cjfr-2013-0380.
166. Hackenberg, C., Muehlkchen, A., Forge, T., and Vrain, T. 2000. Pseudomonas chlororaphis strain Sm3, bacterial antagonist of Pratylenchus penetrans. J Nematol 32: 183–189.
167. Hamou, M.B., Kulhánek, A., Lacombe, S., and Whalen, J.K. 2007. Marking earthworms for release–recapture studies using the trace element rubidium. Applied Soil Ecology 36: 41–45. doi:10.1016/j.apsoil.2006.11.005.
168. Hargreaves, A.L., Whiteside, D.P., and Gilchrist, G. 2011. Concentrations of 17 elements, including mercury, in the tissues, food and abiotic environment of Arctic shorebirds. Science of The Total Environment 409: 3757–3770. doi:10.1016/j.scitotenv.2011.06.043.
169. Harris, M.L., Wilson, L.K., Elliott, J.E., Bishop, C.A., Tomlin, A.D., and Henning, K.V. 2000. Transfer of DDT and metabolites from fruit orchard soils to American robins (Turdus migratorius) twenty years after agricultural use of DDT in Canada. Arch. Environ. Contam. Toxicol. 39: 205–220. doi:10.1007/s002440010098.
170. Hart, M.M., Powell, J.R., Gulden, R.H., Levy-Booth, D.J., Dunfield, K.E., Pauls, K.P., Swanton, C.J., Klironomos, J.N., and Trevors, J.T. 2009. Detection of transgenic cp4 epsps genes in the soil food web. Agron. Sustain. Dev. 29: 497–501. doi:10.1051/agro/2009020.
171. Hawkins, J.W., Lankester, M.W., and Nelson, R.R.A. 1998. Sampling terrestrial gastropods using cardboard sheets. Malacologica 39.
172. Hebert, C.E., Weseloh, D.V., Kot, L., and Glooschenko, V. 1994. Organochlorine contaminants in a terrestrial foodweb on the Niagara Peninsula, Ontario, Canada 1987-89. Arch. Environ. Contam. Toxicol. 26: 356–366. doi:10.1007/BF00203563.
173. van Herk, W., Vernon, R., Moffat, C., and Harding, C. 2008. Response of the Pacific Coast wireworm, Limonius canus, and the dusky wireworm, Agriotes obscurus (Coleoptera: Elateridae), to insecticide-treated wheat seeds in a soil bioassay. phyto 89: 7–19. Société de protection des plantes du Québec (SPPQ). doi:10.7202/000378ar.
174. van Herk, W.G., Vernon, R.S., Labun, T.J., Sevcik, M.H., and Schwinghamer, T.D. 2021. Distribution of pest wireworm (Coleoptera: Elateridae) species in Alberta, Saskatchewan, and Manitoba (Canada). Environ Entomol 50: 663–672. doi:10.1093/ee/nvab006.
175. Holland, T.C., Reynolds, A.G., Bowen, P.A., Bogdanoff, C.P., Marciniak, M., Brown, R.B., and Hart, M.M. 2013. The response of soil biota to water availability in vineyards. Pedobiologia 56: 9–14. doi:10.1016/j.pedobi.2012.08.004.
176. Holliday, N.J., Tomlin, A.D., and Hagley, E.A.C. 1982. Soil fauna of a pest-management apple orchard. Revue d’Écologie et de Biologie du Sol 19: 41–59.
177. Holm, F.A., Zentner, R.P., Thomas, A.G., Sapsford, K., Légère, A., Gossen, B.D., Olfert, O., and Leeson, J.Y. 2006. Agronomic and economic responses to integrated weed management systems and fungicide in a wheat-canola-barley-pea rotation. Can. J. Plant Sci. 86: 1281–1295. NRC Research Press. doi:10.4141/P05-165.
178. Holmstrom, D., Sanderson, K., and Kimpinski, J. 2008a. Effect of tillage regimens on soil erosion, nematodes, and carrot yield in Prince Edward Island. Journal of Soil and Water Conservation 63. doi:10.2489/jswc.63.5.322.
179. Holmstrom, D., Sanderson, K., and Kimpinski, J. 2008b. Effect of tillage regimens on soil erosion, nematodes, and carrot yield in Prince Edward Island. Journal of Soil and Water Conservation 63: 322–328. Taylor & Francis. doi:10.2489/jswc.63.5.322.
180. Huebner, K., Lindo, Z., and Lechowicz, M.J. 2012. Post-fire succession of collembolan communities in a northern hardwood forest. European Journal of Soil Biology 48: 59–65. doi:10.1016/j.ejsobi.2011.10.004.
181. Irshad, U., and Yergeau, E. 2018. Bacterial Subspecies Variation and Nematode Grazing Change P Dynamics in the Wheat Rhizosphere. Front. Microbiol. 9. Frontiers. doi:10.3389/fmicb.2018.01990.
182. Jackson, M., Myrholm, C., Shaw, C., and Ramsfield, T. 2017. Using nested PCR to improve detection of earthworm eDNA in Canada. Soil Biology and Biochemistry 113: 215–218. doi:10.1016/j.soilbio.2017.06.009.
183. Jagdale, G.B., Ball-Coelho, B., Potter, J., Brandle, J., and Roy, R.C. 2000. Rotation crop effects on Pratylenchus penetrans and subsequent crop yields. Can. J. Plant Sci. 80: 543–549. NRC Research Press. doi:10.4141/P99-084.
184. Jayasinghe, B.A.T.D., and Parkinson, D. 2009. Earthworms as the vectors of actinomycetes antagonistic to litter decomposer fungi. Applied Soil Ecology 43: 1–10. doi:10.1016/j.apsoil.2009.06.001.
185. Jegede, O.O., Standen, K.M., Siciliano, S., Lamb, E.G., and Stewart, K.J. 2023. Rocks, lichens, and woody litter influenced the soil invertebrate density in upland tundra heath. PLOS ONE 18: e0282068. Public Library of Science. doi:10.1371/journal.pone.0282068.
186. Jennings, B.W., and Watmough, S.A. 2016. The impact of invasive earthworms on soil respiration and soil carbon within temperate hardwood forests. Ecosystems 19: 942–954. doi:10.1007/s10021-016-9977-y.
187. Jochum, M., Ferlian, O., Thakur, M.P., Ciobanu, M., Klarner, B., Salamon, J.-A., Frelich, L.E., Johnson, E.A., and Eisenhauer, N. 2021. Earthworm invasion causes declines across soil fauna size classes and biodiversity facets in northern North American forests. Oikos 130: 766–780. doi:10.1111/oik.07867.
188. Jochum, M., Thouvenot, L., Ferlian, O., Zeiss, R., Klarner, B., Pruschitzki, U., Johnson, E.A., and Eisenhauer, N. 2022. Aboveground impacts of a belowground invader: How invasive earthworms alter aboveground arthropod communities in a northern North American forest. Biol Lett 18: 20210636. doi:10.1098/rsbl.2021.0636.
189. Johnson, P.W. 1977. Genera of plant parasitic nematodes associated with soybeans on the heavier tebxtured soils of Essex, Kent, and Lambton counties in slouthwestern Ontario. Canadian Plant Disease Survey 57: 19–22.
190. Johnston, H.W., Celetti, M.J., Kimpinski, J., and Platt, H.W. 1994. Fungal pathogens and Pratylenchus penetrans associated with preceding crops of clovers, winter wheat, and annual ryegrass and their influence on succeeding potato crops on Prince Edward Island. American Potato Journal 71: 797–808. doi:10.1007/BF02849375.
191. Jones, A., Fortier, J., Gagnon, D., and Truax, B. 2020. Trading tree growth for soil degradation: Effects at 10 years of black plastic mulch on fine roots, earthworms, organic matter and nitrate in a multi-species riparian buffer. Trees, Forests and People 2: 100032. doi:10.1016/j.tfp.2020.100032.
192. Jorgensen, A., Otani, J., and Evenden, M.L. 2020. Assessment of available tools for monitoring wheat midge (Diptera: Cecidomyiidae). Environ Entomol 49: 627–637. doi:10.1093/ee/nvaa017.
193. Jorna, J., Vandenbrink, B., Hogg, I.D., Wall, D.H., and Adams, B.J. 2023. Metabarcoding inventory of an arctic tundra soil ecosystem reveals highly heterogeneous communities at a small scale. Polar Biol 46: 461–471. doi:10.1007/s00300-023-03131-x.
194. Kabaluk, J., Vernon, R., and Goettel, M. 2007. Mortality and infection of wireworm, Agriotes obscurus [Coleoptera: Elateridae], with inundative field applications of Metarhizium anisopliae. phyto 88: 51–56. Société de protection des plantes du Québec (SPPQ). doi:10.7202/018055ar.
195. Kabaluk, T., Chaigneau, A., and Furlan, L. 2023. Predicting crop damage caused by wireworms and the effect of tillage on trap efficiency. Arthropod-Plant Interactions 17: 517–528. doi:10.1007/s11829-023-09973-w.
196. Kamath, D., Barreto, C., and Lindo, Z. 2022. Nematode contributions to the soil food web trophic structure of two contrasting boreal peatlands in Canada. Pedobiologia 93–94: 150809. doi:10.1016/j.pedobi.2022.150809.
197. Kandel, Y.R., Wise, K.A., Bradley, C.A., Chilvers, M.I., Byrne, A.M., Tenuta, A.U., Faghihi, J., Wiggs, S.N., and Mueller, D.S. 2017. Effect of soybean cyst nematode resistance source and seed treatment on population densities of Heterodera glycines, sudden death syndrome, and yield of soybean. Plant Disease 101: 2137–2143. Scientific Societies. doi:10.1094/PDIS-12-16-1832-RE.
198. Kaneko, N., McLean, M.A., and Parkinson, D. 1995. Grazing preference of Onychiurus subtenuis (Collembola) and Oppiella nova (Oribatei) for fungal species inoculated on pine needles. Pedobiologia 39: 538–546. doi:10.1016/S0031-4056(24)00221-X.
199. Kaufman, C.A., Bennett, J.R., Koch, I., and Reimer, K.J. 2007. Lead Bioaccessibility in Food Web Intermediates and the Influence on Ecological Risk Characterization. Environ. Sci. Technol. 41: 5902–5907. American Chemical Society. doi:10.1021/es062443u.
200. Kernecker, M., Whalen, J.K., and Bradley, R.L. 2015. Agricultural management and flooding shape habitats for non-native earthworms in southern Quebec, Canada. Applied Soil Ecology 96: 240–250. doi:10.1016/j.apsoil.2015.08.011.
201. Kesic, R., Elliott, J.E., Fremlin, K.M., Gauthier, L., Drouillard, K.G., and Bishop, C.A. 2021. Continuing persistence and biomagnification of DDT and metabolites in northern temperate fruit orchard avian food chains. Environ Toxicol Chem 40: 3379–3391. doi:10.1002/etc.5220.
202. Kevan, P.G., Forbes, B.C., Kevan, S.M., and Behan-Pelletier, V. 1995. Vehicle tracks on high Arctic tundra: Their effects on the soil, vegetation, and soil arthropods. Journal of Applied Ecology 32: 655–667. [British Ecological Society, Wiley]. doi:10.2307/2404660.
203. Kimpinski, J., Arsenault, W.J., and Sanderson, J.B. 1997. Fosthiazate for suppression of Pratylenchus penetrans in Potato on Prince Edward Island. J Nematol 29: 685–689.
204. Kimpinski, J., Gallant, C.E., Henry, R., Macleod, J.A., Sanderson, J.B., and Sturz, A.V. 2003. Effect of compost and manure soil amendments on nematodes and on yields of potato and barley: A 7-year study. J Nematol 35: 289–293. [Online] Available: https://pmc.ncbi.nlm.nih.gov/articles/PMC2620642/ [2026 May 1].
205. Kimpinski, J., and Johnston, H.W. 1995. Effects of aldicarb and fungicides on Pratylenchus penetrans populations, root rot and net blotch severity on barley. phyto 76: 9–16. Société de protection des plantes du Québec (SPPQ)l. doi:10.7202/706080ar.
206. Kimpinski, J., Johnston, H.W., and Sanderson, J.B. 1998a. Effects of carbathiin and thiram with propiconazole and aldicarb on grain yield, and on incidence of root rot, leaf blotch, and root lesion nematode in spring wheat. Canadian Journal of Plant Pathology 20: 201–205. Taylor & Francis. doi:10.1080/07060669809500428.
207. Kimpinski, J., Platt, H.W., Perley, S., and Walsh, J.R. 1998b. Pratylenchus spp. and Verticillium spp. in New Brunswick potato fields. Am. J. Pot Res 75: 87–91. doi:10.1007/BF02883882.
208. Kimpinski, J., Plumas, G., and Macdonald, M.C. 1993. Occurrence of the clover cyst nematode, Heterodera trifolii, in Prince Edward Island soils. J Nematol 25: 876–879.
209. Kimpinski, J., and Sanderson, K. 2004. Effects of crop rotations on carrot yield and on the nematodes Pratylenchus penetrans and Meloidogyne hapla. phyto 85: 13–17. Société de protection des plantes du Québec (SPPQ). doi:10.7202/008901ar.
210. Kimpinski, J., and Smith, E.M. 1988. Nematodes in potato soils in New Brunswick. Canadian Plant Disease Survey 68: 147–148.
211. Kimpinski, J., and Sturz, A.V. 1996. Population growth of a rhabditid nematode on plant growth promoting bacteria from potato tubers and rhizosphere soil. J Nematol 28: 682–686.
212. Klein, A., Cameron, E.K., Heimburger, B., Eisenhauer, N., Scheu, S., and Schaefer, I. 2017. Changes in the genetic structure of an invasive earthworm species (Lumbricus terrestris, Lumbricidae) along an urban – rural gradient in North America. Applied Soil Ecology 120: 265–272. doi:10.1016/j.apsoil.2017.08.009.
213. Klironomos, J.N., Bednarczuk, E.M., and Neville, J. 1999. Reproductive significance of feeding on saprobic and arbuscular mycorrhizal fungi by the collembolan, Folsomia candida. Functional Ecology 13: 756–761. doi:10.1046/j.1365-2435.1999.00379.x.
214. Klironomos, J.N., and Kendrick, B. 1995a. Relationships among microarthropods, fungi, and their environment. Plant Soil 170: 183–197. doi:10.1007/BF02183066.
215. Klironomos, J.N., and Kendrick, W.B. 1995b. Stimulative effects of arthropods on endomycorrhizas of sugar maple in the presence of decaying litter. Functional Ecology 9: 528–536. [British Ecological Society, Wiley]. doi:10.2307/2390019.
216. Klironomos, J.N., and Kendrick, W.B. 1996. Palatability of microfungi to soil arthropods in relation to the functioning of arbuscular mycorrhizae. Biol Fertil Soils 21: 43–52. doi:10.1007/BF00335992.
217. Klironomos, J.N., and Moutoglis, P. 1999. Colonization of nonmycorrhizal plants by mycorrhizal neighbours as influenced by the collembolan, Folsomia candida. Biol Fertil Soils 29: 277–281. doi:10.1007/s003740050553.
218. Klironomos, J.N., and Ursic, M. 1998. Density-dependent grazing on the extraradical hyphal network of the arbuscular mycorrhizal fungus, Glomus intraradices, by the collembolan, Folsomia candida. Biol Fertil Soils 26: 250–253. doi:10.1007/s003740050375.
219. Klironomos, J.N., Widden, P., and Deslandes, I. 1992. Feeding preferences of the collembolan Folsomia candida in relation to microfungal successions on decaying litter. Soil Biology and Biochemistry 24: 685–692. doi:10.1016/0038-0717(92)90047-2.
220. Knight, C.C., Pauls, K.P., Sears, M.K., and Schaafsma, A.W. 2005. Oviposition site selected by the western corn rootworm (Diabrotica virgifera virgifera LeConte) in southern Ontario strip plots. Can. J. Plant Sci. 85: 949–954. NRC Research Press. doi:10.4141/P04-159.
221. Kranabetter, J.M., and Chapman, B.K. 1999. Effects of forest soil compaction and organic matter removal on leaf litter decomposition in central British Columbia. Can. J. Soil. Sci. 79: 543–550. NRC Research Press. doi:10.4141/S98-081.
222. Kreutzweiser, D.P., Good, K.P., Chartrand, D.T., Scarr, T.A., Holmes, S.B., and Thompson, D.G. 2008. Effects on litter-dwelling earthworms and microbial decomposition of soil-applied imidacloprid for control of wood-boring insects. Pest Manag Sci 64: 112–118. doi:10.1002/ps.1478.
223. Kuchta, S., Neilsen, D., Zebarth, B.J., Forge, T., and Nichol, C. 2021. Nitrogen, irrigation, and alley management affects raspberry crop response and soil nitrogen and root-lesion nematode dynamics. Soil Science Society of America Journal 85: 1139–1156. doi:10.1002/saj2.20190.
224. Kunelius, H.T., Carter, M.R., Kimpinski, J., and Sanderson, J.B. 1988. Effect of seeding method on alfalfa and red clover establishment and growth, soil physical condition and nematode populations. Soil and Tillage Research 12: 163–175. doi:10.1016/0167-1987(88)90039-6.
225. Labrie, G., Gagnon, A.-È., Vanasse, A., Latraverse, A., and Tremblay, G. 2020. Impacts of neonicotinoid seed treatments on soil-dwelling pest populations and agronomic parameters in corn and soybean in Quebec (Canada). PLOS ONE 15: e0229136. Public Library of Science. doi:10.1371/journal.pone.0229136.
226. Laganière, J., Paré, D., and Bradley, R.L. 2009. Linking the abundance of aspen with soil faunal communities and rates of belowground processes within single stands of mixed aspen–black spruce. Applied Soil Ecology 41: 19–28. doi:10.1016/j.apsoil.2008.08.005.
227. Laigle, I., Moretti, M., Rousseau, L., Gravel, D., Venier, L., Handa, I.T., Messier, C., Morris, D., Hazlett, P., Fleming, R., Webster, K., Shipley, B., and Aubin, I. 2021. Direct and indirect effects of forest anthropogenic disturbance on above and below ground communities and litter decomposition. Ecosystems 24: 1716–1737. doi:10.1007/s10021-021-00613-z.
228. Lapied, E., Nahmani, J., and Rousseau, G.X. 2009. Influence of texture and amendments on soil properties and earthworm communities. Applied Soil Ecology 43: 241–249. doi:10.1016/j.apsoil.2009.08.004.
229. Lazarovits, G., Conn, K.L., and Potter, J. 1999. Reduction of potato scab, verticillium wilt, and nematodes by soymeal and meat and bone meal in two Ontario potato fields. Canadian Journal of Plant Pathology 21: 345–353. Taylor & Francis. doi:10.1080/07060669909501170.
230. LeFait, A., Gailey, J., and Kernaghan, G. 2019. Fungal species selection during ectomycorrhizal grazing by Collembola. Symbiosis 78: 87–95. doi:10.1007/s13199-018-00596-x.
231. Leger, R.G., and Millette, G.J.F. 1977. Properties of four Quebec soils after incubation with five species of earthworms. Can. J. Soil. Sci. 57: 165–172. NRC Research Press. doi:10.4141/cjss77-021.
232. Lejoly, J., Quideau, S., and Laganière, J. 2021. Invasive earthworms affect soil morphological features and carbon stocks in boreal forests. Geoderma 404: 115262. doi:10.1016/j.geoderma.2021.115262.
233. Liang, Q., and Thomson, A.J. 1994. Habitat–abundance relationships of the earthworm Eisenia rosea (Savigny) (Lumbricidae), using principal component regression analysis. Can. J. Zool. 72: 1354–1361. NRC Research Press. doi:10.1139/z94-178.
234. Lindo, Z. 2015. Warming favours small-bodied organisms through enhanced reproduction and compositional shifts in belowground systems. Soil Biology and Biochemistry 91: 271–278. doi:10.1016/j.soilbio.2015.09.003.
235. Lindo, Z. 2020. Transoceanic dispersal of terrestrial species by debris rafting. Ecography 43: 1364–1372. doi:10.1111/ecog.05155.
236. Lindo, Z., and N. Winchester, N. 2008. Scale dependent diversity patterns in arboreal and terrestrial oribatid mite (Acari: Oribatida) communities. Ecography 31: 53–60. doi:10.1111/j.2007.0906-7590.05320.x.
237. Lindo, Z., and Stevenson, S.K. 2007. Diversity and distribution of oribatid mites (Acari: Oribatida) associated with arboreal and terrestrial habitats in interior cedar-hemlock forests, British Columbia, Canada. Northwest Science 81: 305–315. doi:10.3955/0029-344X-81.4.305.
238. Lindo, Z., and Visser, S. 2003. Microbial biomass, nitrogen and phosphorus mineralization, and mesofauna in boreal conifer and deciduous forest floors following partial and clear-cut harvesting. Can. J. For. Res. 33: 1610–1620. NRC Research Press. doi:10.1139/x03-080.
239. Lindo, Z., and Visser, S. 2004. Forest floor microarthropod abundance and oribatid mite (Acari: Oribatida) composition following partial and clear-cut harvesting in the mixedwood boreal forest. Can. J. For. Res. 34: 998–1006. NRC Research Press. doi:10.1139/x03-284.
240. Lindo, Z., and Winchester, N.N. 2006. A comparison of microarthropod assemblages with emphasis on oribatid mites in canopy suspended soils and forest floors associated with ancient western redcedar trees. Pedobiologia 50: 31–41. doi:10.1016/j.pedobi.2005.09.002.
241. Lindo, Z., and Winchester, N.N. 2007. Oribatid mite communities and foliar litter decomposition in canopy suspended soils and forest floor habitats of western redcedar forests, Vancouver Island, Canada. Soil Biology and Biochemistry 39: 2957–2966. doi:10.1016/j.soilbio.2007.06.009.
242. Lindo, Z., and Winchester, N.N. 2009. Spatial and environmental factors contributing to patterns in arboreal and terrestrial oribatid mite diversity across spatial scales. Oecologia 160: 817–825. doi:10.1007/s00442-009-1348-3.
243. Little, L.R., and Maun, M.A. 1997. Relationships among plant-parasitic nematodes, mycorrhizal fungi and the dominant vegetation of a sand dune system. Écoscience 4: 67–74. Taylor & Francis. doi:10.1080/11956860.1997.11682378.
244. Lumley, L.M., Azeria, E.T., Giacobbo, V.A., and Cobb, T.P. 2023. Effects of natural land cover, anthropogenic disturbance, space, and climate on oribatid mite communities in Canada’s oil sands region. Diversity 15: 469. Multidisciplinary Digital Publishing Institute. doi:10.3390/d15040469.
245. Lupardus, R.C., Battigelli, J.P., Janz, A., and Lumley, L.M. 2021. Can soil invertebrates indicate soil biological quality on well pads reclaimed back to cultivated lands? Soil and Tillage Research 213: 105082. doi:10.1016/j.still.2021.105082.
246. Lupwayi, N.Z., Floate, K.D., Schwinghamer, T.D., Hao, X., and Kanashiro, D.A. 2023. Trickle-down effects of the cattle parasiticide ivermectin on soil microfauna on a prairie grassland. Applied Soil Ecology 190: 105021. doi:10.1016/j.apsoil.2023.105021.
247. Lyons, D.B. 1999. Phenology of the native parasitoid Sinophorus megalodontis (Hymenoptera: Ichneumonidae) relative to its introduced host, the pine false webworm (Hymenoptera: Pamphiliidae). The Canadian Entomologist 131: 787–800. doi:10.4039/Ent131787-6.
248. Lyse Benoit, D., Vincent, C., and Chouinard, G. 2006. Management of weeds, apple sawfly (Hoplocampa testudinea Klug) and plum curcuclio (Conotrachelus nenuphar Herbst) with cellulose sheeting. Crop Protection 25: 331–337. doi:10.1016/j.cropro.2005.05.006.
249. Ma, B.L., Meloche, F., and Wei, L. 2009. Agronomic assessment of Bt trait and seed or soil-applied insecticides on the control of corn rootworm and yield. Field Crops Research 111: 189–196. doi:10.1016/j.fcr.2008.12.006.
250. Macherius, A., Lapen, D.R., Reemtsma, T., Römbke, J., Topp, E., and Coors, A. 2014. Triclocarban, triclosan and its transformation product methyl triclosan in native earthworm species four years after a commercial-scale biosolids application. Science of The Total Environment 472: 235–238. doi:10.1016/j.scitotenv.2013.10.113.
251. MacPherson, T., and Bouman, O.T. 2013. Concordance of N bio-indicators: Dandelion and earthworms in a rotational pasture. Ecological Indicators 24: 633–635. doi:10.1016/j.ecolind.2012.08.025.
252. Madani, M., Tenuta, M., and Castillo, P. 2024. The occurrence of cyst nematodes in agricultural fields of Southern Manitoba. Canadian Journal of Plant Pathology 46: 306–317. Taylor & Francis. doi:10.1080/07060661.2023.2293128.
253. Mahran, A., Conn, K.L., Tenuta, M., Lazarovits, G., and Daayf, F. 2008. Effectiveness of liquid hog manure and acidiﬁcation to kill Pratylenchus spp. in soil. Journal of Nematology 40: 266–275.
254. Mahran, A., Tenuta, M., Lumactud, R.A., and Daayf, F. 2009. Response of a soil nematode community to liquid hog manure and its acidification. Applied Soil Ecology 43: 75–82. doi:10.1016/j.apsoil.2009.06.003.
255. Mahran, A., Tenuta, M., Shinners-Carenelly, T., Mundo-Ocampo, M., and Daayf, F. 2010. Prevalence and species identification of Pratylenchus spp. in Manitoba potato fields and host suitability of ‘Russet Burbank.’ Canadian Journal of Plant Pathology 32: 272–282. Taylor & Francis. doi:10.1080/07060661.2010.484218.
256. Malloch, B., Kernaghan, G., and Walker, A.K. 2022. Temporal diversity patterns for fungi and mites associated with decaying Sporobolus pumilus (Spartina patens) in the Minas Basin, Nova Scotia. FACETS 7: 654–673. Canadian Science Publishing. doi:10.1139/facets-2021-0175.
257. Malloch, B., Tatsumi, S., Seibold, S., Cadotte, M.W., and MacIvor, J.S. 2020. Urbanization and plant invasion alter the structure of litter microarthropod communities. Journal of Animal Ecology 89: 2496–2507. doi:10.1111/1365-2656.13311.
258. Marks, C.F., Saidak, W.J., and Johnson, P.W. 1973. Effects of soil management on numbers of the root-lesion nematode pratylenchus penetrans in soils of Ontario peach orchards. Can. J. Plant Sci. 53: 181–185. NRC Research Press. doi:10.4141/cjps73-033.
259. Marshall, C.B., and Lynch, D.H. 2020. Soil microbial and macrofauna dynamics under different green manure termination methods. Applied Soil Ecology 148: 103505. doi:10.1016/j.apsoil.2020.103505.
260. Marshall, V.G. 1974. Seasonal and vertical distribution of soil fauna in a thinned and urea-fertilized Douglas fir forest. Can. J. Soil. Sci. 54: 491–500. NRC Research Press. doi:10.4141/cjss74-062.
261. Marshall, V.G., and Barclay, H.J. 1998. Soil nematodes in successional stages of Douglas-fir forests. Northwest Sceince 71: 98–100.
262. Massicotte, R., Robidoux, P.Y., Sauvé, S., Flipo, D., Fournier, M., and Trottier, B. 2003. Immune response of earthworms (Lumbricus terrestris, Eisenia andrei and Aporrectodea tuberculata) following in situ soil exposure to atmospheric deposition from a cement factory. J. Environ. Monit. 5: 774–779. The Royal Society of Chemistry. doi:10.1039/B301956J.
263. McAdams, B.N., Quideau, S.A., Swallow, M.J.B., and Lumley, L.M. 2018. Oribatid mite recovery along a chronosequence of afforested boreal sites following oil sands mining. Forest Ecology and Management 422: 281–293. doi:10.1016/j.foreco.2018.04.034.
264. McCabe, D.C., Protz, R., and Tomlin, A.D. 1991. Faunal effects on the distribution of gamma emitting radionuclides in four forested soils. Water Air Soil Pollut 57: 521–532. doi:10.1007/BF00282916.
265. McElligott, P.E.K., and Lewis, D.J. 1994. Relative efficiencies of wet and dry extraction techniques for sampling aquatic macroinvertebrates in a Subarctic peatland. The Memoirs of the Entomological Society of Canada 126: 285–289. doi:10.4039/entm126169285-1.
266. McElligott, P.E.K., and Lewis, D.J. 1996a. Development of larvae of Chrysops nigripes Zetterstedt and Chrysops probably zinzalus Philip (Diptera: Tabanidae) in a subarctic Labrador peatland. Can. J. Zool. 74: 1370–1375. NRC Research Press. doi:10.1139/z96-150.
267. McElligott, P.E.K., and Lewis, D.J. 1996b. Distribution and abundance of immature Tabanidae (Diptera) in a subarctic Labrador peatland. Can. J. Zool. 74: 1364–1369. NRC Research Press. doi:10.1139/z96-149.
268. McKeown, A.W., Cerkauskas, R.F., Potter, J.W., and Driel, L. van 1998. Long-term evaluation of cover crop and strip-tillage on tomato yield, foliar diseases and nematode populations. Can. J. Plant Sci. 78: 341–348. NRC Research Press. doi:10.4141/P97-090.
269. McKeown, A.W., and Potter, J.W. 2001. Yield of ‘Superior’ potatoes (Solanum tuberosum) and dynamics of root-lesion nematode (Pratylenchus penetrans) populations following “nematode suppressive” cover crops and fumigation. phyto 82: 13–23. Société de protection des plantes du Québec (SPPQ)l. doi:10.7202/706211ar.
270. Mckercher, R.B., Tollefson, T.S., and Willard, J.R. 1979. Biomass and phosphorus contents of some soil invertebrates. Soil Biology and Biochemistry 11: 387–391. doi:10.1016/0038-0717(79)90052-X.
271. McLean, M.A., Kaneko, N., and Parkinson, D. 1996a. Does selective grazing by mites and collembola affect litter fungal community structure? Pedobiologia 40: 97–105. doi:10.1016/S0031-4056(24)00343-3.
272. McLean, M.A., Kolodka, D.U., and Parkinson, D. 1996b. Survival and growth of Dendrobaena octaedra (Savigny) in pine forest floor materials. Pedobiologia 40: 281–288. doi:10.1016/S0031-4056(24)00371-8.
273. McLean, M.A., and Parkinson, D. 1997. Soil impacts of the epigeic earthworm Dendrobaena octaedra on organic matter and microbial activity in lodgepole pine forest. Can. J. For. Res. 27: 1907–1913. NRC Research Press. doi:10.1139/x97-165.
274. McLean, M.A., and Parkinson, D. 1998a. Impacts of the epigeic earthworm Dendrobaena octaedra on oribatid mite community diversity and microarthropod abundances in pine forest floor: A mesocosm study. Applied Soil Ecology 7: 125–136. doi:10.1016/S0929-1393(97)00041-3.
275. McLean, M.A., and Parkinson, D. 1998b. Impacts of the epigeic earthworm Dendrobaena octaedra on microfungal community structure in pine forest floor: A mesocosm study. Applied Soil Ecology 8: 61–75. doi:10.1016/S0929-1393(97)00067-X.
276. McLean, M.A., and Parkinson, D. 2000. Introduction of the epigeic earthworm Dendrobaena octaedra changes the oribatid community and microarthropod abundances in a pine forest. Soil Biology and Biochemistry 32: 1671–1681. doi:10.1016/S0038-0717(00)00083-3.
277. McTavish, M., Rossi, A., Bourchier, R., and Smith, S. 2022. Quantifying seeds egested by field-collected earthworms: a dynamic and overlooked pool in forest soil seed banks. The Canadian Field-Naturalist 136: 262–267. doi:10.22621/cfn.v136i3.2873.
278. McTavish, M.J., Gorgolewski, A., Murphy, S.D., and Basiliko, N. 2020. Field and laboratory responses of earthworms to use of wood ash as a forest soil amendment. Forest Ecology and Management 474: 118376. doi:10.1016/j.foreco.2020.118376.
279. Meehan, M.L., Barreto, C., Turnbull, M.S., Bradley, R.L., Bellenger, J.-P., Darnajoux, R., and Lindo, Z. 2020. Response of soil fauna to simulated global change factors depends on ambient climate conditions. Pedobiologia 83: 150672. doi:10.1016/j.pedobi.2020.150672.
280. Meehan, M.L., Caruso, T., and Lindo, Z. 2021. Short-term intensive warming shifts predator communities (Parasitiformes: Mesostigmata) in boreal forest soils. Pedobiologia 87–88: 150742. doi:10.1016/j.pedobi.2021.150742.
281. Meehan, M.L., Song, Z., Lumley, L.M., Cobb, T.P., and Proctor, H. 2019. Soil mites as bioindicators of disturbance in the boreal forest in northern Alberta, Canada: Testing taxonomic sufficiency at multiple taxonomic levels. Ecological Indicators 102: 349–365. doi:10.1016/j.ecolind.2019.02.043.
282. Meehan, M.L., Song, Z., and Proctor, H. 2018. Roles of environmental and spatial factors in structuring assemblages of forest-floor Mesostigmata in the boreal region of northern Alberta, Canada. International Journal of Acarology 44: 300–309. Taylor & Francis. doi:10.1080/01647954.2018.1520297.
283. Miller, J.J., Battigelli, J.P., Beasley, B.W., and Drury, C.F. 2017. Response of soil mesofauna to long-term application of feedlot manure on irrigated cropland. Journal of Environmental Quality 46: 185–192. doi:10.2134/jeq2016.08.0318.
284. Miller, J.J., Battigelli, J.P., and Willms, W.D. 2014. Grazing protection influences soil mesofauna in ungrazed and grazed riparian and upland pastures. Rangeland Ecology & Management 67: 429–434. doi:10.2111/REM-D-14-00004.1.
285. Miller, J.J., Owen, M.L., Battigelli, J.P., Drury, C.F., and Chanasyk, D.S. 2020. Short-term legacy effects of feedlot manure application on soil mesofauna. Journal of Environmental Quality 49: 1730–1737. doi:10.1002/jeq2.20152.
286. Miller, J.J., Owen, M.L., Drury, C.F., and Chanasyk, D.S. 2019. Short-term legacy effects of feedlot manure amendments on earthworm abundance in a clay loam soil. Can. J. Soil. Sci. 99: 447–457. NRC Research Press. doi:10.1139/cjss-2019-0022.
287. Mills, A.A.S., and Adl, M.S. 2011. Changes in nematode abundances and body length in response to management intensive grazing in a low-input temperate pasture. Soil Biology and Biochemistry 43: 150–158. doi:10.1016/j.soilbio.2010.09.027.
288. Mills, A.A.S., Price, G.W., and Fillmore, S.A.E. 2020. Responses of nematode, bacterial, and fungal populations to high frequency applications and increasing rates of biosolids in an agricultural soil. Applied Soil Ecology 148: 103481. doi:10.1016/j.apsoil.2019.103481.
289. Mimee, B., Dauphinais, N., and Bélair, G. 2015. Life cycle of the golden cyst nematode, Globodera rostochiensis, in Quebec, Canada. J Nematol 47: 290–295.
290. Mimee, B., Dauphinais, N., and Bélair, G. 2019. “Piler Dirt” survey for the sampling and detection of potato cyst nematodes. Plant Disease 103: 2065–2069. Scientific Societies. doi:10.1094/PDIS-12-18-2188-RE.
291. Mitchell, M.J. 1977. Population dynamics of oribatid mites Acari Cryptostigmata in an aspen woodland soil. Pedobiologia 17: 305–319. doi:https://doi.org/10.1016/S0031-4056(23)00188-9.
292. Mitchell, M.J. 1978. Vertical and horizontal distributions of oribatid mites (Acari: Cryptostigmata) in an aspen woodland soil. Ecology 59: 516–525. Ecological Society of America. doi:10.2307/1936582.
293. Mitchell, M.J. 1979. Energetics of oribatid mites (Acari: Cryptostigmata) in an aspen woodland soil. Pedobiologia 19: 89–98. doi:10.1016/S0031-4056(23)02217-5.
294. Mitchell, M.J., and Parkinson, D. 1976. Fungal feeding or oribatid mites (Acari: Cryptostigmata) in an aspen woodland soil. Ecology 57: 302–312. doi:10.2307/1934818.
295. Mrácek, Z., Kindlmann, P., and Webster, J.M. 2005. Steinernema afﬁne (Nematoda: Steinernematidae), a new record for North America and its distribution relative to other entomopathogenic nematodes in British Columbia. Nematology 7: 495–501.
296. Mrácek, Z., and Webster, J.M. 1993. Survey of heterorhabditidae and steinernematidae (rhabditida, nematoda) in western Canada. J Nematol 25: 710–717.
297. Mulock, B.S., Ellis, C.R., and Whitfield, G.H. 1995. Evaluation of an oviposition trap for monitoring egg populations of Diabrotica spp. (Coleoptera: Chrysomelidae) in field corn. The Canadian Entomologist 127: 839–849. doi:10.4039/Ent127839-6.
298. Munawar, M., Castillo, P., and Yevtushenko, D.P. 2022a. Description of Filenchus species from agroecosystem of southern Alberta, Canada. Agronomy 12: 690. Multidisciplinary Digital Publishing Institute. doi:10.3390/agronomy12030690.
299. Munawar, M., Rahman, A.U., Castillo, P., and Yevtushenko, D.P. 2022b. Morphological and molecular characterization of Nothotylenchus medians and N. similis (Nematoda: Anguinidae) from southern Alberta, Canada. Horticulturae 8: 74. Multidisciplinary Digital Publishing Institute. doi:10.3390/horticulturae8010074.
300. Munro, P., Forge, T.A., Jones, M.D., and Nelson, L.M. 2020. Soil biota from newly established orchards are more beneficial to early growth of cherry trees than biota from older orchards. Applied Soil Ecology 155: 103658. doi:10.1016/j.apsoil.2020.103658.
301. Murray, P., Ge, Y., and Hendershot, W.H. 2000. Evaluating three trace metal contaminated sites: Afield and laboratory investigation. Environ Pollut 107: 127–135. Barking, Essex : 1987. doi:10.1016/s0269-7491(99)00120-7.
302. Nelson, K.L., Lynch, D.H., and Boiteau, G. 2009. Assessment of changes in soil health throughout organic potato rotation sequences. Agriculture, Ecosystems & Environment 131: 220–228. doi:10.1016/j.agee.2009.01.014.
303. Newton, J.S., Glasier, J., Maw, H.E.L., Proctor, H.C., and Foottit, R.G. 2011. Ants and subterranean Sternorrhyncha in a native grassland in east-central Alberta, Canada. The Canadian Entomologist 143: 518–523. doi:10.4039/n11-034.
304. Newton, J.S., and Proctor, H.C. 2013. A fresh look at weight-estimation models for soil mites (Acari). International Journal of Acarology 39: 72–85. Taylor & Francis. doi:10.1080/01647954.2012.744351.
305. Niwranski, K., Kevan, P.G., and Fjellberg, A. 2002. Effects of vehicle disturbance and soil compaction on Arctic collembolan abundance and diversity on Igloolik Island, Nunavut, Canada. European Journal of Soil Biology 38: 193–196. doi:10.1016/S1164-5563(02)01142-1.
306. Nour, S.M., Lawrence, J.R., Zhu, H., Swerhone, G.D.W., Welsh, M., Welacky, T.W., and Topp, E. 2003. Bacteria associated with cysts of the soybean cyst nematode (Heterodera glycines). Applied and Environmental Microbiology 69: 607–615. American Society for Microbiology. doi:10.1128/AEM.69.1.607-615.2003.
307. Nunes, K.A., Fitzpatrick, C.R., and Kotanen, P.M. 2019. Soil biota composition and the performance of a noxious weed across its invaded range. Ecography 42: 1671–1681. doi:10.1111/ecog.04562.
308. Nunes, K.A., and Kotanen, P.M. 2018. Comparative impacts of aboveground and belowground enemies on an invasive thistle. Ecology and Evolution 8: 1430–1440. doi:10.1002/ece3.3751.
309. Nyiraneza, J., Chen, D., Fraser, T., and Comeau, L.-P. 2021. Improving soil quality and potato productivity with manure and high-residue cover crops in eastern Canada. Plants 10: 1436. Multidisciplinary Digital Publishing Institute. doi:10.3390/plants10071436.
310. Olthof, T.H., and Townshend, J.L. 1991. Effect of oxamyl treatment of potato seed pieces on Pratylenchus penetrans and yield. J Nematol 23: 699–705.
311. Osler, G.H.R., Harrison, L., Kanashiro, D.K., and Clapperton, M.J. 2008. Soil microarthropod assemblages under different arable crop rotations in Alberta, Canada. Applied Soil Ecology 38: 71–78. doi:10.1016/j.apsoil.2007.09.003.
312. Otfinowski, R., and Coffey, V. 2020. Can root traits predict communities of soil nematodes in restored northern prairies? Plant Soil 453: 459–471. doi:10.1007/s11104-020-04624-y.
313. Panesar, T.S., Marshall, V.G., and Barclay, H.J. 2000. The impact of clearcutting and partial harvesting systems on population dynamics of soil nematodes in coastal Douglas-fir forests. Pedobiologia 44: 641–665. doi:10.1078/S0031-4056(04)70079-7.
314. Panesar, T.S., Marshall, V.G., and Barclay, H.J. 2001. Abundance and diversity of soil nematodes in chronosequences of coastal Douglas-fir forests on Vancouver Island, British Columbia. Pedobiologia 45: 193–212. doi:10.1078/0031-4056-00080.
315. Pang, A., Rutter, A., Gainer, A., Haack, E., and Zeeb, B. 2023. Assessment of heavily weathered petroleum hydrocarbon-impacted soils to native soil invertebrates from a Canadian subarctic site. J Soils Sediments 23: 2096–2105. doi:10.1007/s11368-023-03466-2.
316. Paquin, P., and Coderre, D. 1997a. Deforestation and fire impact on edaphic insect larvae and other macroarthropods. Environ Entomol 26: 21–30. doi:10.1093/ee/26.1.21.
317. Paquin, P., and Coderre, D. 1997b. Changes in soil macroarthropod communities in relation to forest maturation through three successional stages in the Canadian boreal forest. Oecologia 112: 104–111. doi:10.1007/s004420050289.
318. Parizadeh, M., Kembel, S.W., and Mimee, B. 2023a. Neonicotinoid seed treatments influence soil nematode taxonomic composition and the soil microbial cooccurrence networks. Phytobiomes Journal 7: 478–490. Scientific Societies. doi:10.1094/PBIOMES-12-22-0105-R.
319. Parizadeh, M., Kembel, S.W., and Mimee, B. 2023b. Neonicotinoid seed treatments influence soil nematode taxonomic composition and the soil microbial cooccurrence networks. Phytobiomes Journal 7: 478–490. Scientific Societies. doi:10.1094/PBIOMES-12-22-0105-R.
320. Pentinsaari, M., Blagoev, G.A., Hogg, I.D., Levesque-Beaudin, V., Perez, K., Sobel, C.N., Vandenbrink, B., and Borisenko, A. 2020. A DNA barcoding survey of an Arctic arthropod community: Implications for future monitoring. Insects 11: 46. Multidisciplinary Digital Publishing Institute. doi:10.3390/insects11010046.
321. Perreault, J.M., Eriksen-Hamel, N.S., and Whalen, J.K. 2007. Temporal and spatial dynamics of earthworm surface casting in a temperate soybean agroecosystem. Applied Soil Ecology 37: 10–17. doi:10.1016/j.apsoil.2007.03.007.
322. Perreault, J.M., and Whalen, J.K. 2006. Earthworm burrowing in laboratory microcosms as influenced by soil temperature and moisture. Pedobiologia 50: 397–403. doi:10.1016/j.pedobi.2006.07.003.
323. Pettit, T., Faulkner, K.J., Buchkowski, R.W., Kamath, D., and Lindo, Z. 2023. Changes in peatland soil fauna biomass alter food web structure and function under warming and hydrological changes. European Journal of Soil Biology 117: 103509. doi:10.1016/j.ejsobi.2023.103509.
324. Poprawski, T.J., and Yule, W.N. 1992. Acari associated with Phyllophaga anxia (Leconte) (Coleoptera: Scarabaeidae) in southern Quebec and eastern Ontario. The Canadian Entomologist 124: 397–403. doi:10.4039/Ent124397-2.
325. Porco, D., Skarżyński, D., Decaëns, T., Hebert, P.D.N., and Deharveng, L. 2014. Barcoding the Collembola of Churchill: a molecular taxonomic reassessment of species diversity in a sub-Arctic area. Molecular Ecology Resources 14: 249–261. doi:10.1111/1755-0998.12172.
326. Porter, T.M., Morris, D.M., Basiliko, N., Hajibabaei, M., Doucet, D., Bowman, S., Emilson, E.J.S., Emilson, C.E., Chartrand, D., Wainio-Keizer, K., Séguin, A., and Venier, L. 2019. Variations in terrestrial arthropod DNA metabarcoding methods recovers robust beta diversity but variable richness and site indicators. Sci Rep 9: 18218. Nature Publishing Group. doi:10.1038/s41598-019-54532-0.
327. Porter, T.M., Smenderovac, E., Morris, D., and Venier, L. 2023. All boreal forest successional stages needed to maintain the full suite of soil biodiversity, community composition, and function following wildfire. Sci Rep 13: 7978. Nature Publishing Group. doi:10.1038/s41598-023-30732-7.
328. Potter, J.W., and McKeown, A.W. 2002. Fluctuations of populations of the pin nematode Paratylenchus projectus under selected potato management practices. phyto 83: 147–155. Société de protection des plantes du Québec (SPPQ)l. doi:10.7202/706238ar.
329. Prescott, C.E. 1996. Influence of forest floor type on rates of litter decomposition in microcosms. Soil Biology and Biochemistry 28: 1319–1325. doi:10.1016/S0038-0717(96)00132-0.
330. Price, G.W., and Gordon, A.M. 1998. Spatial and temporal distribution of earthworms in a temperate intercropping system in southern Ontario, Canada. Agroforestry Systems 44: 141–149. doi:10.1023/A:1006213603150.
331. Price, G.W., and Voroney, R.P. 2008. Response to annual applications of de-inked papermill biosolids by field earthworms on three agricultural soils. Applied Soil Ecology 38: 230–238. doi:10.1016/j.apsoil.2007.10.014.
332. Puvanendran, D.C., Larson, D.J., and Thompson, I.D. 1997. Collembola (Arthropoda) of balsam fir [Abies balsamea (L.)] forests of western Newfoundland. The Canadian Entomologist 129: 505–517. doi:10.4039/Ent129505-3.
333. Rafael, O., Victory, C., and Ryan, N. 2023. Dominant plants mediate effects of grazing on soil nematode traits in a wet meadow grassland. Applied Soil Ecology 191: 105047. doi:10.1016/j.apsoil.2023.105047.
334. Raymond-Léonard, L.J., Gravel, D., and Handa, I.T. 2019. A novel set of traits to describe Collembola mouthparts: taking a bite out of the broad chewing mandible classification. Soil Biology and Biochemistry 138: 107608. doi:10.1016/j.soilbio.2019.107608.
335. Raymond-Léonard, L.J., Gravel, D., Reich, P.B., and Handa, I.T. 2018. Springtail community structure is influenced by functional traits but not biogeographic origin of leaf litter in soils of novel forest ecosystems. Proc Biol Sci 285: 20180647. doi:10.1098/rspb.2018.0647.
336. Reeleder, R.D., Miller, J.J., Ball Coelho, B.R., and Roy, R.C. 2006. Impacts of tillage, cover crop, and nitrogen on populations of earthworms, microarthropods, and soil fungi in a cultivated fragile soil. Applied Soil Ecology 33: 243–257. doi:10.1016/j.apsoil.2005.10.006.
337. Rees, H.W., Chow, T.L., and Gregorich, E.G. 2007. Spatial and temporal trends in soil properties and crop yield at a site under intensive up- and down-slope potato production in northwestern New Brunswick. Can. J. Soil. Sci. 87: 383–398. NRC Research Press. doi:10.4141/CJSS07017.
338. Rees, H.W., Chow, T.L., and Gregorich, E.G. 2008. Soil and crop responses to long-term potato production at a benchmark site in northwestern New Brunswick. Can. J. Soil. Sci. 88: 409–422. NRC Research Press. doi:10.4141/CJSS07086.
339. Rees, H.W., Chow, T.L., Zebarth, B., Xing, Z., Toner, P., Lavoie, J., and Daigle, J.-L. 2014. Impact of supplemental poultry manure application on potato yield and soil properties on a loam soil in north-western New Brunswick. Can. J. Soil. Sci. 94: 49–65. NRC Research Press. doi:10.4141/cjss2013-009.
340. Renkema, J., Nyrop, J., Difonzo, C., Sears, M., and Schaafsma, A. 2006. Control decision rule for European chafer (Coleoptera: Scarabaeidae) larvae in field corn. Journal of economic entomology 99: 76–84. doi:10.1603/0022-0493(2006)099%5B0076:CDRFEC%5D2.0.CO;2.
341. Reynolds, L.B., Potter, J.W., and Ball-Coelho, B.R. 2000. Crop rotation with Tagetes sp. is an alternative to chemical fumigation for control of root-lesion nematodes. Agronomy Journal 92: 957–966. doi:10.2134/agronj2000.925957x.
342. Rivest, M., Whalen, J.K., and Rivest, D. 2020. Variation of soil microbial and earthworm communities along an agricultural transect with tree windbreak. Agroforest Syst 94: 1639–1649. doi:10.1007/s10457-019-00476-3.
343. Robidoux, P.Y., Svendsen, C., Sarrazin, M., Thiboutot, S., Ampleman, G., Hawari, J., Weeks, J.M., and Sunahara, G.I. 2005. Assessment of a 2,4,6-trinitrotoluene-contaminated site using Aporrectodea rosea and Eisenia andrei in mesocosms. Arch Environ Contam Toxicol 48: 56–67. doi:10.1007/s00244-004-0217-7.
344. Rochefort, S., Therrien, F., Shetlar, D.J., and Brodeur, J. 2006. Species diversity and seasonal abundance of Collembola in turfgrass ecosystems of North America. Pedobiologia 50: 61–68. doi:10.1016/j.pedobi.2005.10.007.
345. Rousseau, L., Venier, L., Aubin, I., Gendreau-Berthiaume, B., Moretti, M., Salmon, S., and Handa, I.T. 2019. Woody biomass removal in harvested boreal forest leads to a partial functional homogenization of soil mesofaunal communities relative to unharvested forest. Soil Biology and Biochemistry 133: 129–136. doi:10.1016/j.soilbio.2019.02.021.
346. Rousseau, L., Venier, L., Fleming, R., Hazlett, P., Morris, D., and Handa, I.T. 2018a. Long-term effects of biomass removal on soil mesofaunal communities in northeastern Ontario (Canada) jack pine (Pinus banksiana) stands. Forest Ecology and Management 421: 72–83. doi:10.1016/j.foreco.2018.02.017.
347. Rousseau, L., Venier, L., Hazlett, P., Fleming, R., Morris, D., and Handa, I.T. 2018b. Forest floor mesofauna communities respond to a gradient of biomass removal and soil disturbance in a boreal jack pine (Pinus banksiana) stand of northeastern Ontario (Canada). Forest Ecology and Management 407: 155–165. doi:10.1016/j.foreco.2017.08.054.
348. Rutherford, P.M., and Juma, N.G. 1989. Dynamics of microbial biomass and soil fauna in two contrasting soils cropped to barley (Hordeum vulgare L.). Biol Fert Soils 8: 144–153. doi:10.1007/BF00257758.
349. Sackett, T.E., Smith, S.M., and Basiliko, N. 2012. Exotic earthworm distribution in a mixed-use northern temperate forest region: Influence of disturbance type, development age, and soils. Can. J. For. Res. 42: 375–381. NRC Research Press. doi:10.1139/x11-195.
350. Sackett, T.E., Smith, S.M., and Basiliko, N. 2013. Indirect and direct effects of exotic earthworms on soil nutrient and carbon pools in North American temperate forests. Soil Biology and Biochemistry 57: 459–467. doi:10.1016/j.soilbio.2012.08.015.
351. Saguez, J., Latraverse, A., De Almeida, J., van Herk, W.G., Vernon, R.S., Légaré, J.-P., Moisan-De Serres, J., Fréchette, M., and Labrie, G. 2017. Wireworm in Quebec field crops: Specific community composition in North America. Environ Entomol 46: 814–825. doi:10.1093/ee/nvx116.
352. Sampedro, L., Jeannotte, R., and Whalen, J.K. 2006. Trophic transfer of fatty acids from gut microbiota to the earthworm Lumbricus terrestris L. Soil Biology and Biochemistry 38: 2188–2198. doi:10.1016/j.soilbio.2006.02.001.
353. Sanderson, J.B., MacLeod, J.A., and Kimpinski, J. 1999. Glyphosate application and timing of tillage of red clover affects potato response to N, soil N profile, and root and soil nematodes. Can. J. Soil. Sci. 79: 65–72. NRC Research Press. doi:10.4141/S98-028.
354. Scheffczyk, A., Floate, K.D., Blanckenhorn, W.U., Düring, R.-A., Klockner, A., Lahr, J., Lumaret, J.-P., Salamon, J.-A., Tixier, T., Wohde, M., and Römbke, J. 2016. Nontarget effects of ivermectin residues on earthworms and springtails dwelling beneath dung of treated cattle in four countries. Environ Toxicol Chem 35: 1959–1969. doi:10.1002/etc.3306.
355. Scheu, S., and Parkinson, D. 1994. Effects of earthworms on nutrient dynamics, carbon turnover and microorganisms in soils from cool temperate forests of the Canadian Rocky Mountains — laboratory studies. Applied Soil Ecology 1: 113–125. doi:10.1016/0929-1393(94)90031-0.
356. Scheuhammer, A.M., Bond, D.E., Burgess, N.M., and Rodrigue, J. 2003. Lead and stable lead isotope ratios in soil, earthworms, and bones of American woodcock (Scolopax minor) from eastern Canada. Environ Toxicol Chem 22: 2585–2591. doi:10.1897/02-361.
357. Scott, I.M., Gagnon, N., Lesage, L., Philogène, B.J.R., and Arnason, J.T. 2005. Efficacy of botanical insecticides from Piper species (Piperaceae) extracts for control of Ruropean chafer (Coleoptera: Scarabaeidae). J Econ Entomol 98: 845–855. doi:10.1603/0022-0493-98.3.845.
358. Scott, T., Bradley, R.L., and Bourgault, P. 2024. Non-native earthworms increase the abundance and diet quality of a common woodland salamander in its northern range. Biol Invasions 26: 187–200. doi:10.1007/s10530-023-03168-3.
359. Simard, L., Belair, G., Powers, T., Tremblay, N., and Dionne, J. 2008. Incidence and population density of plant-parasitic nematodes on golf courses in Ontario and Québec, Canada. Journal of Nematology 40: 241–251.
360. Simard, L., Bélair, G., Stock, S.P., Mauléon, H., and Dionne, J. 2007a. Natural occurrence of entomopathogenic nematodes (Rhabditida: Steinernematidae) on golf courses in eastern Canada. Nematol 9: 325–332. doi:10.1163/156854107781352070.
361. Simard, L., Brodeur, J., and Dionne, J. 2007b. Distribution, abundance, and seasonal ecology of Listronotus maculicollis (Coleoptera: Curculionidae) on golf courses in Quebec, Canada. Journal of economic entomology 100: 1344–52. doi:10.1603/0022-0493(2007)100%5B1344:DAASEO%5D2.0.CO;2.
362. Smenderovac, E., Hoage, J., Porter, T.M., Emilson, C., Fleming, R., Basiliko, N., Hajibabei, M., Morris, D., and Venier, L. 2023. Boreal forest soil biotic communities are affected by harvesting, site preparation with no additional effects of higher biomass removal 5 years post-harvest. Forest Ecology and Management 528: 120636. doi:10.1016/j.foreco.2022.120636.
363. Smith, C.A.S., Tomlin, A.D., Miller, J.J., Moore, L.V., Tynen, M.J., and Coates, K.A. 1990. Large enchytraeid (Annelida: Oligochaeta) worms and associated fauna from unglaciated soils of the northern Yukon, Canada. Geoderma 47: 17–32. doi:10.1016/0016-7061(90)90045-B.
364. Smith, J.L., Baute, T.S., and Schaafsma, A.W. 2020. Quantifying early-season pest injury and yield protection of insecticide seed treatments in corn and soybean production in Ontario, Canada. J Econ Entomol 113: 2197–2212. doi:10.1093/jee/toaa132.
365. Smith, J.L., Baute, T.S., Sebright, M.M., Schaafsma, A.W., and DiFonzo, C.D. 2018. Establishment of Striacosta albicosta (Lepidoptera: Noctuidae) as a primary pest of corn in the Great Lakes region. J Econ Entomol 111: 1732–1744. doi:10.1093/jee/toy138.
366. Speratti, A.B., and Whalen, J.K. 2008. Carbon dioxide and nitrous oxide fluxes from soil as influenced by anecic and endogeic earthworms. Applied Soil Ecology 38: 27–33. doi:10.1016/j.apsoil.2007.08.009.
367. St. John, M.G., Bagatto, G., Behan-Pelletier, V., Lindquist, E.E., Shorthouse, J.D., and Smith, I.M. 2002. Mite (Acari) colonization of vegetated mine tailings near Sudbury, Ontario, Canada. Plant and Soil 245: 295–305. doi:10.1023/A:1020453912401.
368. Stock, P.S., Mrácek, Z., and Webster, J.M. 2000. Morphological variation between allopatric populations of Steinernema kraussei (Steiner, 1923) (Rhabditida: Steinernematidae). Nematology 2: 143–152. doi:10.1163/156854100509033.
369. Straube, D., Johnson, E.A., Parkinson, D., Scheu, S., and Eisenhauer, N. 2009. Nonlinearity of effects of invasive ecosystem engineers on abiotic soil properties and soil biota. Oikos 118: 885–896. doi:10.1111/j.1600-0706.2009.17405.x.
370. Sturz, A.V., and Kimpinski, J. 1999. Effects of fosthiazate and aldicarb on populations of plant-growth-promoting bacteria, root-lesion nematodes and bacteria-feeding nematodes in the root zone of potatoes. Plant Pathology 48: 26–32. doi:10.1046/j.1365-3059.1999.00305.x.
371. Sutherland, J.R., and Sluggett, L.J. 1973. Time, temperature, and soil moisture effects on Xiphinema bakeri nematode survival in fallow soil. Phytopathology 64: 507–513.
372. Sylvain, Z.A., and Buddle, C.M. 2010. Effects of forest stand type on oribatid mite (Acari: Oribatida) assemblages in a southwestern Quebec forest. Pedobiologia 53: 321–325. doi:10.1016/j.pedobi.2010.03.001.
373. Sylvain, Z.A., and Mosseler, A. 2017. Use of shrub willows (Salix spp.) to develop soil communities during coal mine restoration. Can. J. For. Res. 47: 1687–1694. NRC Research Press. doi:10.1139/cjfr-2017-0196.
374. Tan, C.S., Drury, C.F., Soultani, M., van Wesenbeeck, I.J., Ng, H.Y.F., Gaynor, J.D., and Welacky, T.W. 1998. Effect of controlled drainage and tillage on soil structure and tile drainage nitrate loss at the field scale. Water Science and Technology 38: 103–110. doi:10.1016/S0273-1223(98)00503-4.
375. Tanney, J.B., and Hutchison, L.J. 2011. Encapsulation and immobilization of a mycophagous nematode by two Sphaerobolus species. Botany 89: 745–751. NRC Research Press. doi:10.1139/b11-060.
376. Therrien, F., Chagnon, M., and Hébert, C. 1999. Biodiversity of Collembola in sugar maple (Aceraceae) forests. Can Entomol 131: 613–628. doi:10.4039/Ent131613-5.
377. Thompson, A.R., and Sans, W.W. 1974. Effects of soil insecticides in southwestern Ontario on non-target invertebrates: Earthworms in pasture. Environ Entomol 3: 305–308. doi:10.1093/ee/3.2.305.
378. Thomson, A.J., and Davies, D.M. 1974. Mapping methods for studying soil factors and earthworm distribution. Oikos 25: 199–203. [Nordic Society Oikos, Wiley]. doi:10.2307/3543642.
379. Tomlin, A.D. 1977. Pipeline construction - impact on soil micro- and mesofauna (Arthropoda and Annelida) in Ontario. Proceedings of the Entomological Society of Ontario 108: 13–17.
380. Tomlin, A.D. 1980. Effects on soil fauna of the fungicide, benomyl, used to control earthworm populations around an airport. Protection Ecology 2: 325–330.
381. Tomlin, A.D., Mccabe, D., and Protz, R. 1992. Species composition and seasonal variation of earthworms and their effect on soil properties in southern Ontario, Canada. Soil Biology and Biochemistry 24: 1451–1457. doi:10.1016/0038-0717(92)90132-H.
382. Tomlin, A.D., and Miller, J.J. 1987. Composition of the soil fauna in forested and grassy plots at Delhi, Ontario. Can. J. Zool. 65: 3048–3055. NRC Research Press. doi:10.1139/z87-462.
383. Tomlin, A.D., and Miller, J.J. 1988. Impact of ring-billed gull (Larus delawarensis Ord.) foraging on earthworm populations of southwestern Ontario agricultural soils. Agriculture, Ecosystems & Environment 20: 165–173. doi:10.1016/0167-8809(88)90108-9.
384. Tomlin, A.D., Protz, R., Martin, R.R., McCabe, D.C., and Lagace, R.J. 1993. Relationships amongst organic matter content, heavy metal concentrations, earthworm activity, and soil microfabric on a sewage sludge disposal site. Geoderma 57: 89–103. doi:10.1016/0016-7061(93)90149-F.
385. Tomlin, A.D., Tolman, J.H., and Thorn, G.D. 1980. Suppression of earthworm (Lumbricus terrestris) populations around an airport by soil application of the fungicide, Benomyl. Protection Ecology 2: 319–323.
386. Tousignant, S., and Coderre, D. 1992. Niche partitioning by soil mites in a recent hardwood plantation in southern Québec, Canada. Pedobiologia 36: 287–294.
387. Tu, C.M., Marks, C.F., Johnson, P.W., and Elliot, J.M. 1995a. Effects of nematicides on populations of soil microflora and Pratylenchus penetrans, soil enzymes and flue-cured tobacco. J. of Env. Sc. & Hlth., Part B 30: 95–111. doi:10.1080/03601239509372929.
388. Tu, C.M., Marks’, C.F., Johnson, P.W., Gayed, S.K., and Elliot, J.M. 1995b. Effects of pesticides on soil enzymatic activities, Pratylenchus penetrans populations, black root rot, and growth of flue‐cured tobacco. Journal of Environmental Science and Health, Part B: Pesticides, Food Contaminants, and Agricultural Wastes 30: 141–162. Taylor & Francis. doi:10.1080/03601239509372931.
389. Turnbull, M.S., and Lindo, Z. 2015. Combined effects of abiotic factors on Collembola communities reveal precipitation may act as a disturbance. Soil Biology and Biochemistry 82: 36–43. doi:10.1016/j.soilbio.2014.12.007.
390. Turney, S., Altshuler, I., Whyte, L.G., and Buddle, C.M. 2018. Macroinvertebrate and soil prokaryote communities in the forest–tundra ecotone of the Subarctic Yukon. Polar Biol 41: 1619–1633. doi:10.1007/s00300-018-2330-5.
391. Ulmer, B.J., and Dosdall, L.M. 2006a. Spring emergence biology of the cabbage seedpod weevil (Coleoptera: Curculionidae). Ann Entomol Soc Am 99: 64–69. doi:10.1603/0013-8746(2006)099%5B0064:SEBOTC%5D2.0.CO;2.
392. Ulmer, B.J., and Dosdall, L.M. 2006b. Emergence of overwintered and new generation adults of the crucifer flea beetle, Phyllotreta cruciferae (Goeze) (Coleoptera: Chrysomelidae). Crop Protection 25: 23–30. doi:10.1016/j.cropro.2005.03.003.
393. Utkhede, R.S., Vrain, T.C., and Yorston, J.M. 1992. Effects of nematodes, fungi and bacteria on the growth of young apple trees grown in apple replant disease soil. Plant Soil 139: 1–6. doi:10.1007/BF00012835.
394. VandenBygaart, A.J., Protz, R., Tomlin, A.D., and Miller, J.J. 1998. 137Cs as an indicator of earthworm activity in soils. Applied Soil Ecology 9: 167–173. doi:10.1016/S0929-1393(98)00071-7.
395. VandenBygaart, A.J., Protz, R., Tomlin, A.D., and Miller, J.J. 1999. Tillage system effects on near-surface soil morphology: observations from the landscape to micro-scale in silt loam soils of southwestern Ontario. Soil and Tillage Research 51: 139–149. doi:10.1016/S0167-1987(99)00050-1.
396. Vernon, R.S., Van Herk, W.G., Clodius, M., and Harding, C. 2013. Further studies on wireworm management in Canada: damage protection versus wireworm mortality in potatoes. J Econ Entomol 106: 786–799. doi:10.1603/ec12180.
397. Vrain, T., DeYoung, R., Hall, J., and Freyman, S. 1996. Cover crops resistant to root-lesion nematodes in raspberry. HortSci 31: 1195–1198. doi:10.21273/HORTSCI.31.7.1195.
398. Vrain, T.C., Forge, T.A., and Young, R.D. 1997. Population dynamics of Pratylenchus penetrans parasitizing raspberry. Fundam. appl. Nematol. 20: 29–36.
399. Wall, D.H., Bradford, M.A., St. John, M.G., Trofymow, J.A., Behan-Pelletier, V., Bignell, D.E., Dangerfield, J.M., Parton, W.J., Rusek, J., Voigt, W., Wolters, V., Gardel, H.Z., Ayuke, F.O., Bashford, R., Beljakova, O.I., Bohlen, P.J., Brauman, A., Flemming, S., Henschel, J.R., Johnson, D.L., Jones, T.H., Kovarova, M., Kranabetter, J.M., Kutny, L., Lin, K.-C., Maryati, M., Masse, D., Pokarzhevskii, A., Rahman, H., Sabará, M.G., Salamon, J.-A., Swift, M.J., Varela, A., Vasconcelos, H.L., White, D., and Zou, X. 2008. Global decomposition experiment shows soil animal impacts on decomposition are climate-dependent. Global Change Biology 14: 2661–2677. doi:10.1111/j.1365-2486.2008.01672.x.
400. Watmough, S., and Meadows, M. 2014. Do earthworms have a greater influence on nitrogen dynamics than atmospheric nitrogen deposition? Ecosystems 17. doi:10.1007/s10021-014-9794-0.
401. Watson, T.T., Forge, T.A., and Nelson, L.M. 2018a. Pseudomonads contribute to regulation of Pratylenchus penetrans (Nematoda) populations on apple. Can. J. Microbiol. 64: 775–785. NRC Research Press. doi:10.1139/cjm-2018-0040.
402. Watson, T.T., Nelson, L.M., and Forge, T.A. 2018b. Preplant soil incorporation of compost to mitigate replant disease: Soil biological factors associated with plant growth promotion in orchard soil. Compost Science & Utilization 26: 286–296. Taylor & Francis. doi:10.1080/1065657X.2018.1540953.
403. Watson, T.T., Nelson, L.M., Neilsen, D., Neilsen, G.H., and Forge, T.A. 2017. Soil amendments influence Pratylenchus penetrans populations, beneficial rhizosphere microorganisms, and growth of newly planted sweet cherry. Applied Soil Ecology 117–118: 212–220. doi:10.1016/j.apsoil.2017.04.014.
404. Welke, S.E., and Parkinson, D. 2003. Effect of Aporrectodea trapezoides activity on seedling growth of Pseudotsuga menziesii, nutrient dynamics and microbial activity in different forest soils. Forest Ecology and Management 173: 169–186. doi:10.1016/S0378-1127(02)00010-5.
405. Whalen, J., and Costa, C. 2003. Linking spatio-temporal dynamics of earthworm populations to nutrient cycling in temperate agricultural and forest systems. Pedobiologia 47: 801–806. doi:10.1078/0031-4056-00262.
406. Whalen, J., Sampedro, L., and Waheed, T. 2004. Quantifying surface and subsurface cast production by earthworms under controlled laboratory conditions. Biology and Fertility of Soils 39: 287–291. doi:10.1007/s00374-003-0715-1.
407. Whalen, J.K. 2004. Spatial and temporal distribution of earthworm patches in corn field, hayfield and forest systems of southwestern Quebec, Canada. Applied Soil Ecology 27: 143–151. doi:10.1016/j.apsoil.2004.04.004.
408. Whalen, J.K., Benslim, H., Elmi, A.A., and Husk, B.R. 2021. Earthworm populations are stable in temperate agricultural soils receiving wood-based biochar. Pedosphere 31: 398–404. doi:10.1016/S1002-0160(20)60080-7.
409. Whalen, J.K., Han, L., and Dutilleul, P. 2015. Burrow refilling behavior of Aporrectodea turgida (Eisen) and Lumbricus terrestris L. as revealed by X-ray computed tomography scanning: Graphical and quantitative analyses. Can. J. Soil. Sci. 95: 231–235. NRC Research Press. doi:10.4141/cjss-2014-040.
410. Whalen, J.K., and Janzen, H.H. 2002. Labeling earthworms uniformly with 13C and 15N: implications for monitoring nutrient fluxes. Soil Biology and Biochemistry 34: 1913–1918. doi:10.1016/S0038-0717(02)00207-9.
411. Wilkinson, A.T.S., Finlayson, D.G., and Campbell, C.J. 1977. Soil incorporation of insecticides for control of wireworms in potato land in British Columbia. Journal of Economic Entomology 70: 755–758.
412. Willard, J.R., and Petrovich, M.S. 1975. Efficiency of the centrifugal-flotation method for extraction of nematodes from clay soils. Plant Dis. Reptr. 59: 314–318.
413. Winchester, N., and Ring, R. 1996. Northern temperate coastal Sitka spruce forests with special emphasis on canopies: Studying arthropods in an unexplored frontier. Northwest Science 70: 94–103. [Online] Available: https://www.semanticscholar.org/paper/Northern-temperate-coastal-Sitka-spruce-forests-on-Winchester-Ring/e6ab26b5fd786630ae1dc27214298253baba8bd9 [2026 May 1].
414. Winter, J.P., Voroney, R.P., and Ainsworth, D.A. 1990. Soil microarthropods in long-term no-tillage and conventional tillage corn production. Can. J. Soil. Sci. 70: 641–653. NRC Research Press. doi:10.4141/cjss90-067.
415. Wironen, M., and Moore, T.R. 2006. Exotic earthworm invasion increases soil carbon and nitrogen in an old-growth forest in southern Quebec. Can. J. For. Res. 36: 845–854. NRC Research Press. doi:10.1139/x06-016.
416. Young, E.H., Abedin, J., and Unc, A. 2023. Soil nematode trophic structure and biochar addition in recently converted boreal lands. Soil Res 61: 456–467. doi:10.1071/SR22228.
417. Young, E.H., Vallotton, J.D., Kedir, A.J., Medaiyese, A.O., Goyer, C., Comeau, L.-P., and Unc, A. 2022. The impacts of rock pulverization on soil quality and functional soil nematode and respiration properties of boreal lands converted from forest to agricultural use. Can. J. Soil. Sci. 102: 977–990. NRC Research Press. doi:10.1139/cjss-2022-0007.
418. Young, M.R., Behan-Pelletier, V.M., and Hebert, P.D.N. 2012. Revealing the hyperdiverse mite fauna of Subarctic Canada through DNA barcoding. PLOS ONE 7: e48755. Public Library of Science. doi:10.1371/journal.pone.0048755.
419. Young, M.R., and Hebert, P.D.N. 2022. Unearthing soil arthropod diversity through DNA metabarcoding. PeerJ 10: e12845. PeerJ Inc. doi:10.7717/peerj.12845.
420. Young, M.R., Proctor, H.C., deWaard, J.R., and Hebert, P.D.N. 2019. DNA barcodes expose unexpected diversity in Canadian mites. Mol Ecol 28: 5347–5359. doi:10.1111/mec.15292.
421. Yu, Q., Potter, J.W., and Gilby, G. 1998. Plant-parasitic nematodes associated with turfgrass in golf courses in southern Ontario. Canadian Journal of Plant Pathology 20: 304–307. Taylor & Francis. doi:10.1080/07060669809500397.
422. Yu, Q., Tsao, R., Chiba, M., and Potter, J. 2007. Oriental mustard bran reduces Pratylenchus penetrans on sweet corn. Canadian Journal of Plant Pathology 29: 421–426. Taylor & Francis. doi:10.1080/07060660709507488.
423. Yu, Q., Ye, W., and Powers, T. 2016. Morphological and molecular characterization of Gracilacus wuae n. sp. (Nematoda: Criconematoidea) associated with cow parsnip (Heracleum maximum) in Ontario, Canada. J Nematol 48: 203–213. doi:10.21307/jofnem-2017-028.
424. Yu, Q., Ye, W., Sun, F., and Miller, S. 2010. Characterization of Globodera rostochiensis (Tylenchida: Heteroderidae) associated with potato in Quebec, Canada. Canadian Journal of Plant Pathology 32: 264–271. Taylor & Francis. doi:10.1080/07060661003740322.
425. Yurgel, S.N., Douglas, G.M., Comeau, A.M., Mammoliti, M., Dusault, A., Percival, D., and Langille, M.G.I. 2017. Variation in bacterial and eukaryotic communities associated with natural and managed wild blueberry habitats. Phytobiomes Journal 1: 102–113. Scientific Societies. doi:10.1094/PBIOMES-03-17-0012-R.
426. Zaitlin, B., Hayashi, M., and Clapperton, J. 2007. Distribution of northern pocket gopher burrows, and effects on earthworms and infiltration in a prairie landscape in Alberta, Canada. Applied Soil Ecology 37: 88–94. doi:10.1016/j.apsoil.2007.04.003.
427. Zasada, I., Peetz, A., and Forge, T. 2017. Pratylenchus species associated with blueberry (Vaccinium spp.) and weed species in the Pacific North-west of North America. Canadian Journal of Plant Pathology 39: 497–502. Taylor & Francis. doi:10.1080/07060661.2017.1367724.
