## Supplementary figures and images for "A systematic map of soil invertebrate biodiversity in Canada: Identifying spatial and taxonomic knowledge gaps"

### supplementary files_S2_ROSES flow diagram

**ROSES Flow Diagram for Systematic Maps. Version 1.0**

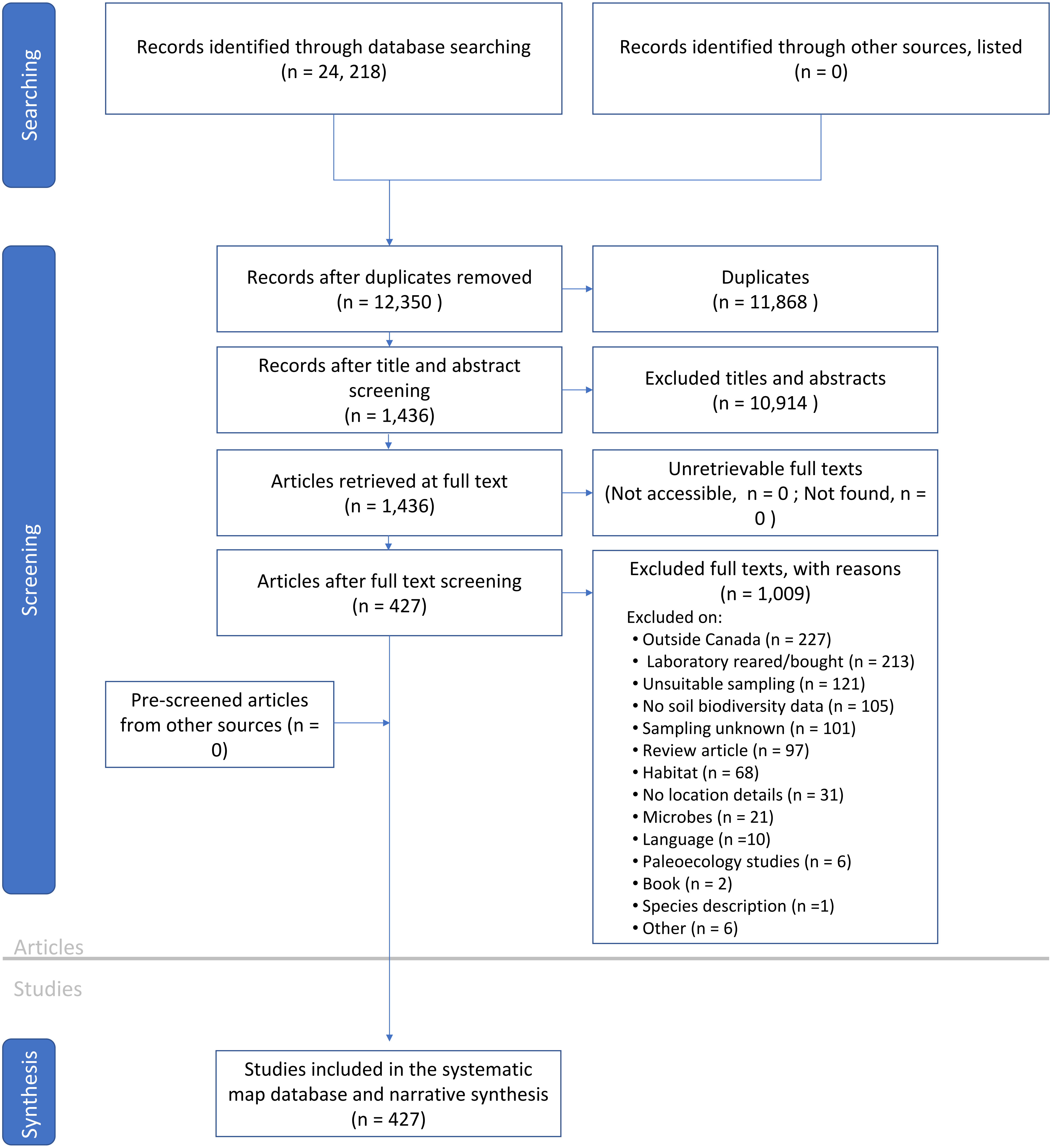
